# PiP-plex: A Particle-in-Particle System for Multiplexed Quantification of Proteins Secreted by Single Cells

**DOI:** 10.1101/2025.03.26.645608

**Authors:** Félix Lussier, Byeong-Ui Moon, Mojra Janta-Polczynski, Yonatan Morocz, Fabian Svahn, Molly Shen, Ljuboje Lukic, Lidija Malic, Teodor Veres, Andy Ng, David Juncker

## Abstract

Cell signaling is modulated by the secretion of various proteins, which can be used to infer a cell’s phenotype. However, these proteins cannot be readily detected in multiplex by commonly used methods at the single cell level. Here, we present PiP-plex, a particles-in-particle (PiPs) system comprising (i) fluorescence intensity barcoded microparticles (BMPs) co-entrapped with (ii) a single cell inside an alginate hydrogel particle for multiplex protein secretion analysis by confocal microscopy. We show that developed PiPs maintained >90 % cellular viability and allowed live cells retrieval. A seven-plex fluorescent barcoding and concomitant sandwich immunoassay in PiPs were implemented with limits of detection ranging from 0.8 pg mL^-1^ to 2 ng mL^-1^ depending on the protein. PiP-plex assays were benchmarked with bulk immunoassays and found to rival or outperform them. We applied PiP-plex to analyze protein secreted by THP-1 cells upon exposure to lipopolysaccharide and detected varying cell responses, with a significant increase in MIP-1α, TNF-α and IL-17A. Multivariate analysis revealed that the majority of stimulated cells secreted either MIP-1α or IL-17A, while other cytokines were typically co-secreted. Using PiP-plex, we analyzed ∼750 THP-1 cells, showcasing its potential for characterizing cells and cell-based therapeutics for cancer immunotherapies.

## Introduction

Cell signaling plays a central role in health and disease, as cells secrete proteins and vesicles to communicate and orchestrate collective behavior. For instance, immune cells encountering pathogenic material or stimulated by neighboring cells can expand into a heterogeneous repertoire of functionally distinct types to modulate the immune response. These diverse cellular phenotypes are reflected in the composition and abundance of secreted cytokines and chemokines,^[1–3]^ key modulators of immune functions. Hence, cell secretion contains promising information for fundamental cell biology studies, as well as for diagnostic and therapeutic monitoring. However, commonly employed bulk-type measurements only record ensemble averages and fail to capture single cell heterogeneity known to drive systemic responses.^[4]^ So-called polyfunctional cells that co-secrete multiple proteins have been identified as major contributors to the efficacy of cell immunotherapy in the treatment of hematological malignancies.^[5]^ There is a desire to identify polyfunctional cells as an indicator of therapeutic potential, as well as to isolate and proliferate them to produce more effective cell therapies and conduct further analysis.^[6, 7^] However, current single cell assays used to identify polyfunctional cells tend to be destructive, and thus cells are not retrievable for proliferation. There is thus an unmet need to detect multiple proteins secreted from a single cell, while retaining its viability and to permit subsequent isolation and clonal expansion.

Quantification of single cell protein secretions is typically performed by segregating cells in microfluidic chambers,^[1, 2^] water-in-oil (W/O) droplets,^[8–14]^ or hydrogels.^[15–18]^ Within the confined space, secreted proteins are captured on arrays of capture antibodies (cAb),^[19]^ or on nano-^[9–11]^ to micrometer-sized^[12, 20, 21^] beads and assayed by sandwich immunoassay using fluorescently labeled detection antibodies (dAbs). In the case of droplets, microfluidics are used to manipulate and sort W/O droplets based on assay results recorded as the fluorescence signal of droplets and cells.^[22, 23^] However, sandwich immunoassays performed in W/O droplets co-encapsulate labeled dAb that contribute to a high background, which limits the sensitivity and multiplexing capacity. Moreover, meticulous optimization of droplet size for optimal cell viability, which depends on the cell type, may be required.^[24]^ Many limitations were mitigated by technologies employing hydrogels to trap cells for immunoassays. Cells are either directly entrapped inside the hydrogel matrix during gel formation using microfluidic droplet generators,^[8, 25^] or trapped within a templated cavity (e.g. nanovials^[17, 26, 27^]). Due to the porosity of hydrogels and the open geometry of nanovials, assay reagents for single cell immunoassays can be easily added and washed while maintaining cell viability. Additionally, hydrogels are compatible with flow cytometry (FCM) for high throughput quantification and sorting of selected cell population.

Droplets and nanovials can be combined with downstream sequencing to correlate protein secretion with gene expression using DNA-barcoded antibodies.^[15, 16^] In this case, multiplex secretion profiles can be accomplished, but cells are not retrievable. While multiplexed fluorescent readouts of secreted proteins can preserve cell viability, they suffer from spectral overlap between fluorescent dyes and weak signals due to low concentration. Hence, the level of multiplexing remains low, whereas only two^[28]^- or tri-plex^[8]^ assays have been reported to date. Spatial segregation of secreted antibodies inside a microfluidic chamber patterned with antibody arrays can increase multiplexing and is now commercially available as a single cell barcoded chip.^[2]^ However, cells are not retrievable for further analysis.

We recently reported the development of intensity barcoded microparticles (BMPs) enabling high multiplex sandwich immunoassay (MSA) for quantification of proteins by FCM (see Supplementary Note 1).^[29, 30^] By using DNA oligos as building blocks, double-stranded DNA (dsDNA) oligos harboring both a 5’ biotin and a 3’ fluorescent dye modifications were bound to streptavidin (SA)-coated microparticles in the presence of biotinylated cAbs to serve as an immunoassay support. Using this approach, we demonstrated that the amount of labeled dsDNA supplied to the microparticles was stoichiometrically recapitulated on the surface of the particles, enabling the generation of >500 distinct multicolor BMPs and used for bulk immunoassays.

In this work, we introduce a novel particle-in-particle (PiPs) system for single cell encapsulation and multiplex quantification of their secreted proteins (referred to as PiP-plex) (Figure 1A). To generate PiPs, we co-encapsulate single cells with a panel of BMPs, each functionalized with an antibody against a specific cytokine, into a selectively permeable alginate hydrogel particle using droplet microfluidics (Figure 1B). The BMPs entrapped in PiPs are used as probes to measure multiple cytokines secreted by single cells using sandwich immunoassays (Figure 1C), and thus assess the functional diversity of single cells. The use of permeable alginate permits diffusive perfusion and rinsing of nutrients and assay reagents, allowing for sensitive, single cell assays while maintaining cell viability and enabling downstream retrieval and expansion. To analyze the PiPs, we developed a processing pipeline that includes imaging of PiPs and BMPs using confocal laser scanning microscopy (CLSM), 3D segmentation of BMPs, barcode decoding, and quantification of secreted cytokines. Due to the transparency of alginate hydrogels, PiPs lack a visible physical boundary, making it difficult to determine which cells and which BMPs are co-encapsulated. We leveraged the limited diffusivity of large molecules through alginate hydrogel to localize each PiP using microscopy. By supplementing fluorescent molecules larger than 2MDa to the buffer suspension containing PiPs, we increased the background fluorescence, thereby enhancing the contrast between the PiPs and the surrounding medium. This enabled automated image segmentation with the alginate hydrogel particles appearing as dark spheres. Next, cells and BMPs were assigned to individual PiPs based on whether their XY coordinates fell within the boundaries of a specific PiP. Their secretion levels were then quantified using the corresponding fluorescence assay signal. Using PiP-plex, we quantified up to seven cytokines secreted by single cells to assess their functional diversity (Figure 1D). As a proof-of-concept, we assessed the functional diversity of THP-1 macrophage (Mφ)-like cells upon stimulation by lipopolysaccharide (LPS), identifying multiple distinct cell phenotypes while preserving cell viability.

**Figure 1:**
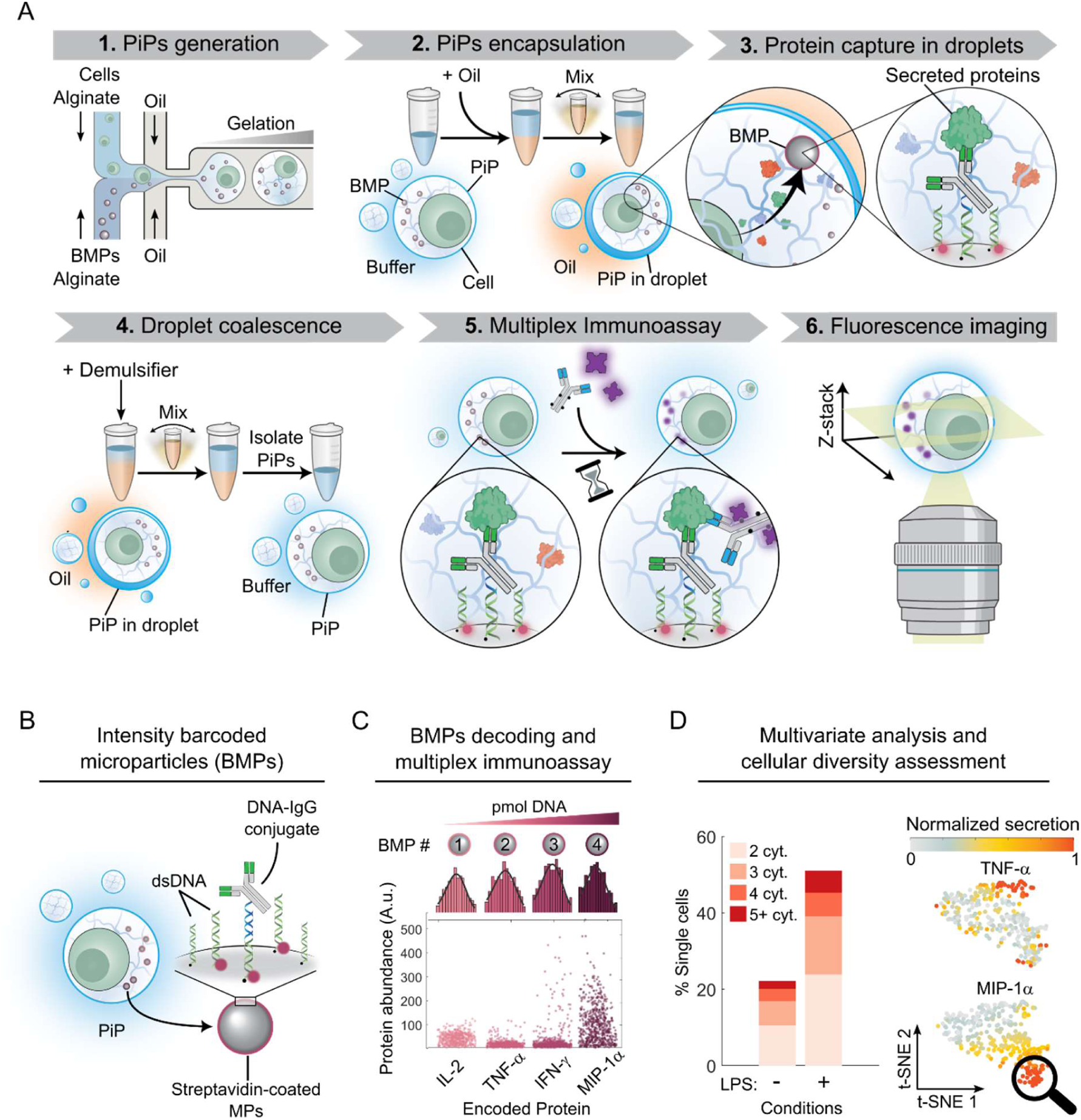
Multiplexed quantification of proteins secreted by a single cell with PiP-plex. (**A**) Schematic representation of the PiP-plex pipeline including six key steps: **1.** Co-encapsulation of barcoded microparticles (BMPs) and a single cell and soluble alginate in W/O droplets. On-chip crosslinking of alginate to form particles-in-particle (PiPs) hydrogel entrapping a single cell and multiple BMPs. **2.** PiPs are isolated, washed, and resuspended in buffer prior being encapsulated in *de novo* droplets. **3.** The encapsulated PiPs enable the capture of secreted proteins by single cell onto BMP harboring a capture antibody. **4.** Following incubation, droplets are coalesced to isolate and resuspend the PiPs in buffer. **5.** PiPs are sequentially incubated with assay reagents for multiplex immunoassays. **6.** Viable cells are labeled, and PiPs are contrast-enhanced by supplementing a fluorescent dye unable to diffuse through the hydrogel matrix. PiPs are imaged by fluorescence confocal microscopy. (**B**) A generalized construct of BMPs composed of biotinylated and fluorescently labeled dsDNA and DNA-IgG conjugate bound onto streptavidin-coated microparticles (MPs). (**C**) BMPs are barcoded by increasing the amount of labeled dsDNA on their surface and used for multiplex detection of secreted proteins by decoding the BMPs and measuring the detection antibody signal on each BMP entrapped in PiPs. (**D**) Data generated by PiP-plex enables the assessment of cellular diversity, such as the number of individual cytokines secreted by single cells and multivariate analysis for further exploration.

## Results and discussion

### BMPs can be automatically localized and decoded by CLSM

Beads are commonly analyzed using FCM, which allows for highly multiplex protein quantification by individually interrogating beads labeled with fluorescent dyes. However, when using beads to analyze protein secreted by single cells, several considerations need to be addressed. First, beads must be barcoded and functionalized with capture antibodies to enable highly sensitive and selective immunoassays. Second, multiple distinct beads must be co-isolated with a single cell to capture secreted proteins while preventing cross-contamination from neighbouring cells. Third, it is essential to trace the captured proteins to their cell of origin. To address these requirements, we encapsulated multiple BMPs, each functionalized with a different antibody, within a permeable hydrogel particle, referred to as PiPs, using droplet-based microfluidic (Figure 1A). Herein, microfluidics enables the co-encapsulation of a single cell with multiple BMPs into discreate volumes, while the hydrogel matrix spatially immobilized both the cell and BMPs within the W/O droplets. The resulting hydrogels are permeable, allowing them to support downstream sandwich immunoassay and can be readily isolated and manipulated for further analysis.

However, FCM cannot be used to analyze PiPs containing multiple BMPs as FCM interrogates entire PiPs rather than individual beads. Consequently, multiplexing would be impaired due to the spectral overlap of barcoding dyes and immunoassay reporters. To overcome this limitation, an image-based method was required to spatially resolve individual BMPs within each PiP. While imaging FCM offers some spatial resolution, its limited *Z-axis* resolution hinders accurate detection of BMPs that are stochastically distributed in three-dimensional space. Conversely, CLSM features high spatial resolution in all three dimensions (*XYZ)*. We hence employed CLSM to analyze BMPs within PiPs and explored its potential to quantify fluorescence signals from individual beads, analogous to FCM.

We generated single-color BMPs using double-stranded encoder oligos (dsEO_i_) harboring a single fluorescent dye encoder. Each encoder dye *i* was conjugated to the 3’ end of a 21-nucleotides (nt) DNA oligo (EO*_i_*) and annealed to a complementary 5’ biotinylated oligo (SO) to produce dsEO_i_. We also incorporated a non-fluorescent encoder oligo (EO_0_) to balance and conserve the overall amount of dsEOs to be bound onto the particles (Figure S1A; Supplementary Note 1). During the annealing of SO and EOs, a small excess of EOs was provided to ensure complete hybridization of the biotinylated oligo. Then, mixtures of dsEO_i_ and dsEO_0_ along with biotinylated IgG were bound onto streptavidin (SA)-coated microparticles to form BMPs (Figure S1B). We anticipate a quantitative conversion of the SA-coated microparticles into BMPs given the high affinity between the biotinylated dsEOs and SA. The excess and non-hybridized EOs were removed through repeated cycles of magnetic aggregation and resuspension. The recapitulation of the dsEOs composition from solution to the particle surface enabled precise encoding of fluorescence intensity on the BMPs. The encoded fluorescence intensities were then used for protein encoding by incorporating biotinylated IgG during BMP synthesis, enabling subsequent protein quantification through immunoassay.

To evaluate the suitability of CLSM for single particle detection, we used both conventional FCM dyes (FAM, Cy3 and Cy5) and microscopy-favored dyes (ATTO488, ATTO550 and AF647), known for their higher irradiance and photostability, to generate BMPs, while employing similar excitation/emission configurations for analysis (Figure S2A). For each single-color BMP, four standards were generated by varying the dsEO_i_:dsEO_0_ ratio. In all cases, we fixed the total amount of double-stranded oligos (dsEO_0_ + dsEO_i_) to 90 pmol, based on our previous work.^[29]^ BMPs were electrostatically immobilized on a poly-L-lysine (PLL) coated coverslip to minimize Brownian motion and drift during imaging (Figure 2A). Upon fluorescence CLSM imaging, we noted a highly homogeneous background signal at various excitation wavelengths, particularly at Ex/Em of 561/590 nm. We attributed this background to light scattering due to the presence of a magnetic core within SA-coated Dynabeads® (Figure S2B). Microscopy images of non-paramagnetic beads composed of the same polymeric material and affinity protein (i.e. SA-coated polystyrene) showed minimal background/scattering of the beads when imaged under the same conditions, confirming our hypothesis (Figure S2C). We used this intrinsic, homogeneous scattering signal to localize each BMP centroid by applying a 3D spherical Hough transform directly to the 3D data acquired by z-stacking of the 561/590 nm channel (Figure 2B).^[31]^ We then generated and applied a 3.0 μm diameter spherical mask to the uniformly-sized BMPs (∼2.8 μm, size distribution < 3 %), with each mask centered on the detected spheroid, followed by image segmentation using a 3D watershed algorithm. The resulting masks were used to integrate the mean fluorescence intensity (MFI) in 3D of each BMP for each fluorescence channel. The usage of the intrinsic 561/590 nm signal to segment all individual BMP overcomes the challenges related to the segmentation of the BMPs exhibiting low fluorescence intensity barcodes with insufficient signal intensity for reliable segmentation. BMPs generated with dyes employing the 561/590 nm Ex/Em configuration (e.g., ATTO550 and Cy3) did not interfere with the segmentation. Our results showed that BMPs’ fluorescence signal increased compared to the background signal of BMPs functionalized with only dsEO_0_, highlighted by the monotonic increase of the signal-to-background ratio (Figure 2C). Also, single particle fluorescence signals measured by CLSM were consistent with the ones obtained by FCM.

**Figure 2:**
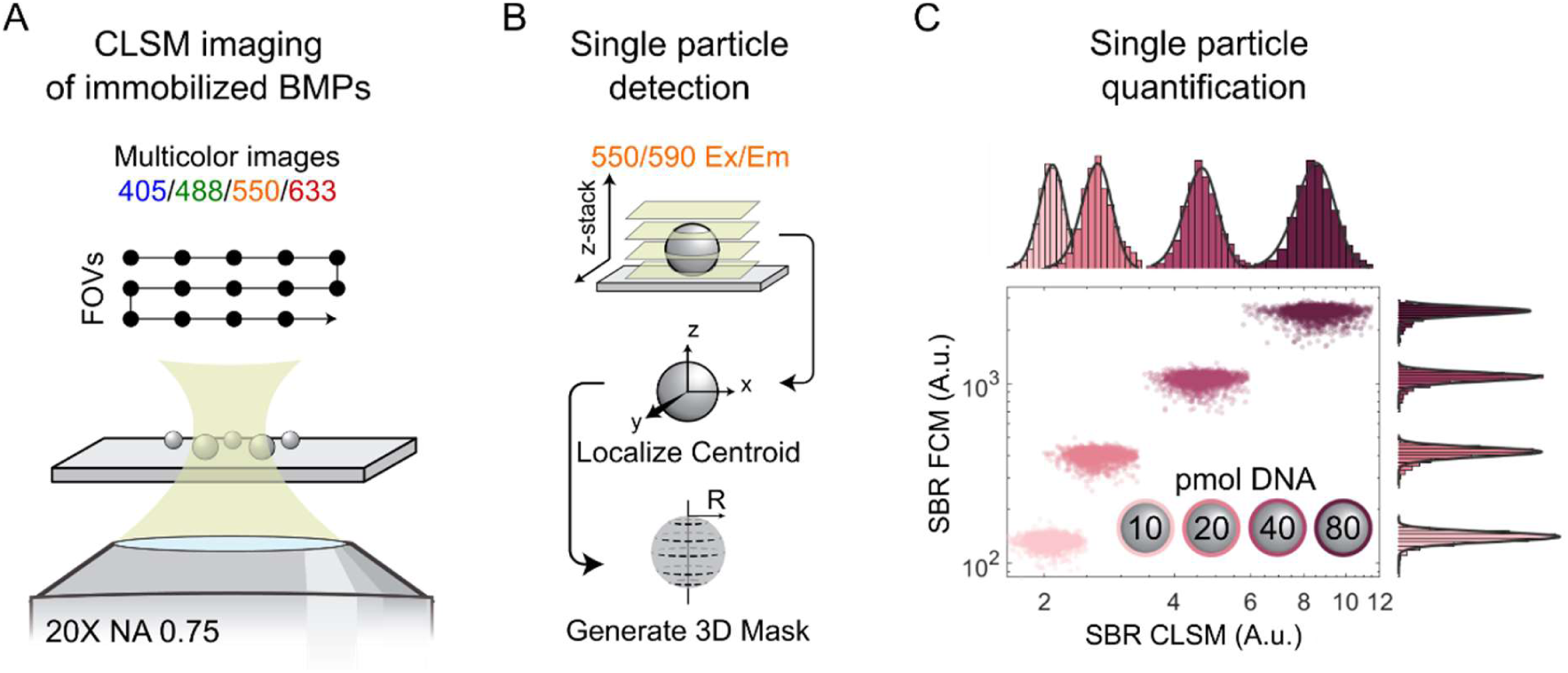
CLSM segmentation and decoding of BMPs directly in 3D. (**A**) Multicolor imaging of immobilized BMPs by CLSM. Multiple fields of view (FOVs) are acquired, each including multicolor *Z*-stack. (**B**) Single particle detection by CLSM. Each BMP centroid is directly localized in 3D via a spherical Hough transform on the 561/590 nm Ex/Em Z-stack images. Spherical masks with a radius *R* of 1.5 μm are centered on each BMP centroid and applied to assess the volumetric fluorescence intensity of BMPs in each fluorescence channel. (**C**) Signal-to-background ratio (SBR) of individual, single-color BMPs (AF647) detected by CLSM in 3D, and FCM (*n >* 5000 BMPs per barcode).

Results showed that for both FCM and CLSM, the ATTO dyes and AF647 exhibited higher brightness and monotonic increase in signal upon increasing the dsEO_i_:dsEO_0_ ratio, whereas we noted a deviation in linearity at high dsEO_i_:dsEO_0_ ratio for Cy3 and Cy5, attributed to self-quenching of the dyes (Figure S3). For all tested dyes, the BMP signals in FCM and CLSM were correlated with Pearson’s *r* values ranging from 0.84 to 0.99 (Figure S4). The coefficients of variation (%CV) were smaller for CLSM compared to FCM across all dyes, and AF647 showed the lowest variation (∼6% CV; Figure S5A-B). We attributed this observation to the favorable properties of AF647, including its high irradiance, superior photostability compared to other dyes, and minimal self-quenching at high concentrations.^[32]^ Additionally, the hydrophilicity of AF647 may reduce nonspecific interactions with the SA-coated beads, hence minimizing %CV. Taken together, these results confirm that CLSM is capable of detecting and discriminating fluorescence signals of individual BMPs.

### Novel BMPs construct improves single-color barcoding capacity and enables reliable antibody encoding for multiplex sandwich immunoassay

Antibody pairs are commonly supplied as a pristine antibody to be immobilized on a solid support for analyte capture (cAb), and a biotinylated Ab intended for detection (dAb), which facilitates the subsequent binding of fluorescently or enzymatically labeled proteins for signal generation. In our previous efforts to generate BMPs for MSAs, we reversed the dAb and cAb by immobilizing biotinylated IgG on the surface of the MPs (Figure S1), while pristine IgGs were used as dAbs, requiring a different strategy to generate the assay signal.^[29]^ The co-immobilization of biotinylated cAb and dsEOs for barcoding introduced a binding competition between cAb and dsEOs that was not previously considered. Such competition could skew either cAb density and/or dsEOs density, affecting assay performance and barcoding accuracy.

To reduce binding competition, we introduced a new strategy that immobilizes only dsEOs, which are used for both barcoding and subsequent antibody immobilization by hybridization of a cAb-DNA conjugate (Figure 3). In conjunction with our previously used SO, we introduced a 5’-biotinylated oligos composed of 42 nt, referred to as capture oligo (CO), possessing both the complementary sequence to the EO_i_ and an additional 21 nt sequence (Figure 3A). Both CO and SO were annealed to their respective EO_i_ to generate pairs of dsEO_i,_ referred to as dsCO_i_ and dsSO_i_ respectively. Again, we kept the total amount of dsDNA at 90 pmol for all BMPs, where the usage of dsCO_i_ and dsSO_i_ now enabled a precise and predictable control of the surface density of anchoring points on the BMPs. We fixed the total amount of dsCOs and dsSOs to 20 and 70 pmol, respectively, based on previous investigation using similar DNA-based construct for MSA.^[30]^ We observed that removing the biotinylated cAb during the binding of dsEOs to the MPs resulted in an overall increase in the number of dsEOs bound to the MPs’ surface (Figure S6). We generated single-color BMPs (AF647) using both our novel construct (cAb_oligo_) and the original co-adsorption method with biotinylated antibody (cAb_biotin_; Figure S6A), and measured their fluorescence using FCM and CLSM (Figure S6B-C). We noted an increased fluorescence signal for the cAb_oligo_ construct compared to cAb_biotin_ while preserving high correlation between the two methods (Pearson’s *r =* 0.99; Figure S6D). All AF647 BMPs generated with the cAb_oligo_ construct had a lower %CV when compared to their cAb_biotin_ counterparts (Figure S7) that we atrributed to the reduction in binding competition between the biotinylated reagents and the beads’ surface for the cAb_oligo_ constructs.

**Figure 3:**
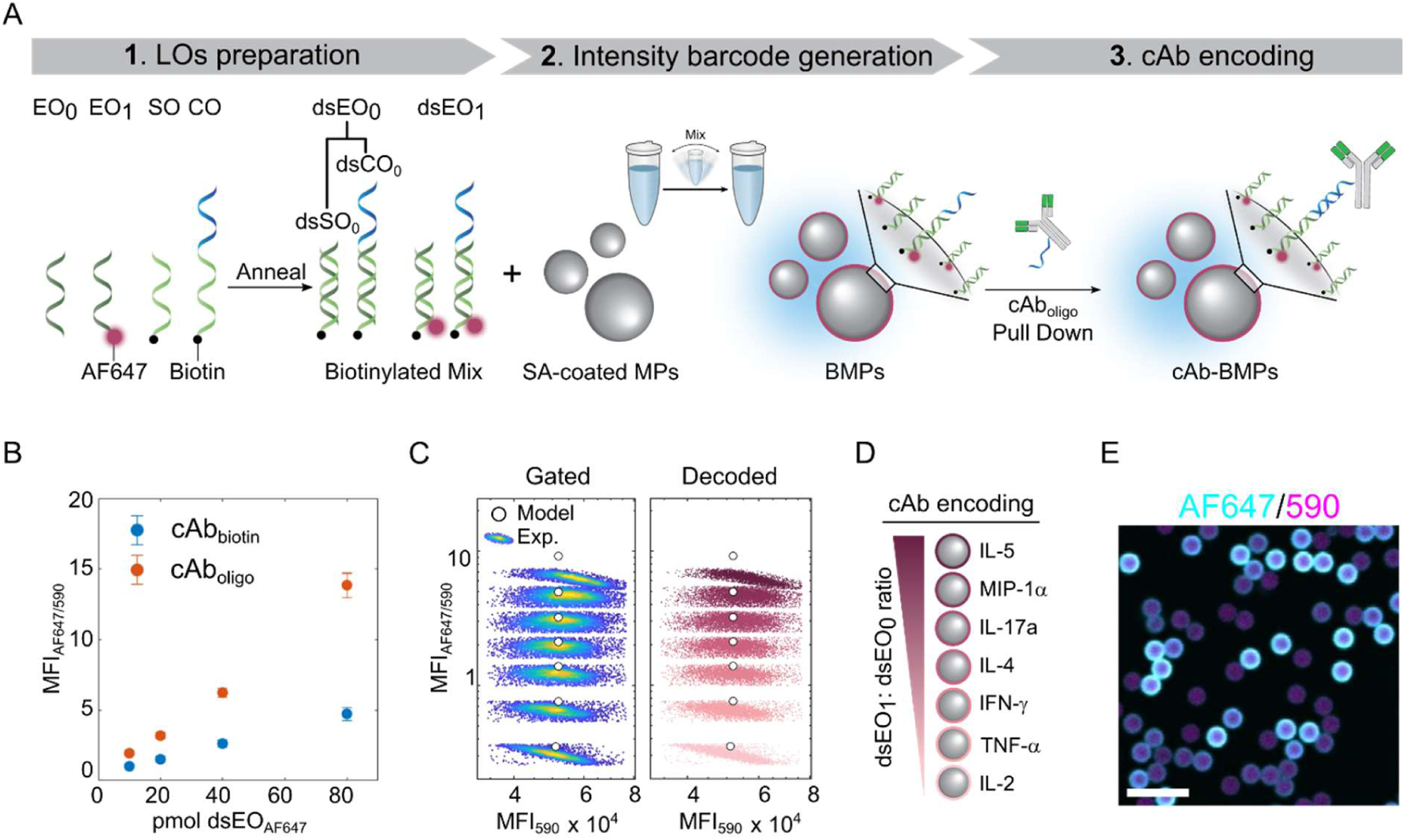
BMPs with high barcoding-capacity based on a cAb_oligo_ for multiplexed protein immunoassays. (**A**) Overview schematic of the three-steps in cAb-BMP assembly: **1.** Encoder oligos corresponding to unlabeled (EO_o_) and fluorescently labeled oligonucleotides (EO_1_) are each annealed individually in the presence of a spacer oligo (SO) and a capture oligo (CO) to generate pairs of biotinylated double-stranded encoder oligo (dsEO_0_ and dsEO_1_). Each dsEO_i_ is a mixture of double-stranded spacer (dsSO_i_) and capture (dsCO_i_) annealed with their corresponding *i* encoder. For each dsEO_i_, dsSO_i_ and dsCO_i_ were combined at a 7:2 ratio. **2.** SA-coated MPs are then rapidly mixed with a mixture of dsEO_0_/dsEO_1_ at a specified ratio to generate the BMPs, where the dsEO_s_ are captured onto the MPs due to the high SA-biotin affinity. By varying the ratio of dsEO_1_:dsEO_0_, while maintaining the overall amount of dsEOs on the surface of the MPs constant (dsEO_1_ + dsEO_0_ = 90 pmol), the distinct fluorescence intensity barcodes can be generated. **3.** The cAb_oligo_ conjugates are pulled down by incubating the cAb_oligo_ in presence of BMPs to capture the antibodies onto the BMPs for antibody encoding and downstream MSA. (**B**) Calibration curves of single color (AF647) cAb-BMPs measured by CLSM, normalized by the scattering of the MPs at 590 nm. BMPs were synthesized either via co-adsorption of biotinylated cAb onto the BMPs (cAb_biotin_) or through cAb_oligo_ pull-down using the novel BMP construct (cAb_oligo_). Mean ± S.D. (*N* = 3, *n* ≥ 5000 BMPs) are presented. (**C**) Single-color (AF647) cAb-BMPs detected in multiplex by CLSM. Poorly segmented BMPs are gated-out based on their MFI_590_. Individual BMPs are decoded using a gaussian mixture model, initiated by the predicted intensity of the BMPs (solid white circles). (**D**) Antibody encoding of the distinct BMPs, generated at various dsEO_1_:dsEO_0_ ratios. (**E**) CLSM image of pooled single-color cAb-BMPs of various dsEO_i_:dsEO_0_ ratio. Scale Bar: 10 μm.

Interestingly, after normalizing the AF647 signal of individual BMPs by their scattering signal (561/590 nm), we observed a significant reduction in variance (Figure 3B) and %CV for both cAb_oligo_ and cAb_biotin_ when measured by CLSM compared to FCM (Figure S7). We hypothesized that normalizing fluorescence intensity by the scattering of the beads could correct two key sources of variability. First, it could account for the discrepancies between the sampling planes in the *Z-*stack and the actual 3D positions of individual beads, as scattering reflects their spatial placement. Second, normalization could reduce variability in fluorescence intensity caused by bead polydispersity, since scattering correlates with bead size and structure. The scattering-based normalization was particularly effective for AF647 for two reasons. First, the pseudo scattering signal at 590 nm was immune to bleed-through from AF647. The signal measured at 590 nm was affected by the bleed-through of dyes in the 488 channels (FAM/ATTO488), thus skewing the results. Secondly, AF647 already exhibited the lowest %CV for barcoding compared to other dyes, such as Cy5, as discussed above.

Due to the quantitative recapitulation of the dsEOs proportion in solution on the particles’ surface,^[29]^ we modeled a series of dsEO_i_:dsEO_0_ ratios *in silico,* exhibiting minimal overlap in the BMPs intensity distribution for both FCM and CLSM (Supplementary Note 2). The proportion of dsEO_AF647_ per barcode was modeled based on the measured %CV of AF647 BMPs, with barcodes separated by ∼3.5 times their expected standard deviation. We hence expected to include over 99% of a given barcode population from a normal distribution corresponding to a tolerance of less than 1 %. Using these assumptions, we predicted the generation of 3 and 5 well-separated barcodes when using cAb_biotin_ and cAb_oligo_ structure, respectively, for BMPs measured by CLSM. When measured by FCM, we predicted 5 barcodes for both constructs, corresponding to the same barcoding capacity for single-color BMPs previously reported (Figure S8).^[29]^ However, by using the AF647/590 ratio instead of the AF647 intensity to reduce the experimental %CV, we predicted the generation of up to 7 distinct barcodes with minimal overlap for the cAb_oligo_ construct when measured by CLSM (Figure S8), corresponding to a 1.5- to 2-fold improvement in single-color barcoding capacity. We validated experimentally the generation of these 7 distinct single-color barcodes and used their predicted intensity value measured by CLSM to automatically decode BMPs by a gaussian mixture model (GMM; Figure 3C). We noted this improvement in barcoding capacity solely in the case of single-color barcode measured by CLSM and generated with the novel cAb_oligo_ construct. The 7 BMPs could then be used for antibody encoding, whereas each distinct BMP served to immobilize cAb-DNA conjugate (Figure 3D). This approach enabled the encoding of a specific cAb to a given intensity value, and direct decoding by GMM upon imaging of pooled cAb-BMPs (Figure 3E).

Next, we assessed the analytical performance of our new BMPs structure as a bead-based support for the sandwich immunoassay. We generated a cAb_oligo_ by conjugating a 30-nt oligo sequence (Supplementary Table S1) to a mouse anti-goat cAb (Supplementary Table S2). Following the removal of unconjugated oligos, the cAb_oligo_ was anchored on the surface of BMPs via pull-down. We confirmed the successful hybridization of our cAb_oligo_ conjugate onto all BMPs by CLSM using a BV421-labeled goat anti-mouse dAb (Figure S9). We also measured uniform BV421 labeling across all BMPs, confirming that the hybridization of cAb_oligo_ is independent of the dsEO_AF647_ proportion used for barcoding. We then assayed a single cytokine (IL-5) using an anti-IL-5 cAb_oligo_ and benchmarked our performance against FCM (Figure S10). We used a biotinylated anti-IL-5 dAb in conjunction with labeled SA to generate assay signal (Figure S10A). Similar to the fluorescent barcoding read-out, we also used BMPs’ scattering signal at 561/590 nm to normalize the CLSM volumetric assay signal. We observed comparable limits of detection (LOD) of 84 and 33 pg mL^-1^ for recombinant IL-5 in buffer using FCM and CLSM, respectively, with comparable dynamic ranges (Figure S10B). Our results indicate that the assay performance was preserved between FCM and CLSM for our cAb_oligo_-based BMP.

We then compared the performance for of our novel BMP construct (cAb_oligo_) with the previous cAb_biotin_ BMP construct in a sandwich immunoassay for IFN-γ detection in buffer. Both BMP types were tested using the same antibody pair. In the cAb_biotin_ construct, biotinylated IgG was immobilized on the beads for antigen capture, while the pristine IgG was used for detection. Conversely, the cAb_oligo_ construct used the pristine IgG conjugated to a DNA anchor, which was hybridized onto the BMPs for antigen capture, while the biotinylated IgG served as the detection antibody (Figure S11A). The assay signal was generated using either BV421-labeled secondary Abs or BV421-labeled SA. We observed a LOD of 47 and 109 pg mL^-1^ for recombinant IFN-γ using the cAb_oligo_ and cAb_biotin_, respectively, representing a 2-fold improvement with cAb_oligo_. We hypothesize that this enhancement is partly due to the DNA-based anchoring strategy used in cAb_olig._ After hybridization to the beads, the DNA linker conserved a 9-nts single stranded sequence, which may allow the cAb to adopt a more favorable orientation for efficient antigen binding. Additionally, DNA-mediated anchoring may reduce local steric hindrance, further improving capture efficiency. The use of SA as opposed to a secondary Ab may also contribute to the improved assay performance given SA’s higher avidity, specificity and affinity compared to conventional antibodies. However, we noted an increase in fluorescence background signals when using labeled SA compared to labeled secondary Ab (Figure S11B). This background was attributed to multivalent binding of biotinylated dAb to unoccupied SA binding sites on the beads in absence of antigen (Figure S12A-B). Pre-blocking the residual SA binding sites with free biotin prior to immunoassay significantly reduced this nonspecific background signal (Figure S12B), without affecting the intensity of the fluorescent barcodes (Figure S12C).

We then applied our cAb_oligo_ assembly to assess the multiplexing capability of our construct. We generated a seven-plex protein panel comprising interleukin-2 (IL-2), IL-4, IL-17A, IL-5, tumor necrosis factor-α (TNFα), interferon-γ (IFN-γ), and chemokine (C-C motif) ligand-3 (CCL3/MIP-1α). These cytokines were selected for their role as immune stimulatory, effector, regulatory and inflammatory signaling molecules, which are highly relevant for single cell secretion profiling.^[5]^ We conjugated the same 30 nt sequence to all the different cAbs (Supplementary Table S2) with a coupling yield ranging from 60-85% across our antibody panel to produce various cAb_oligo_ conjugates. These conjugates were subsequently anchored onto specific BMPs (Figure 3D). After appropriate blocking with biotin to prevent nonspecific interactions (Figure S13), we assessed potential cross-reactivity of our antibody pairs on the cAb-BMPs (Figure S14).^[33]^ We confirmed low cross-reactivity at both low (1 ng mL^-1^; Figure S14A, C) and high (100 ng mL^-1^; Figure S14B, D) antigen concentrations, confirming the specificity of our antibody pairs and validating the suitability our approach for MSA targeting the selected protein panel.

Taken together, our new construct supported the generation of 7 single-color BMPs, only achievable by CLSM. In addition, our construct can be used for sandwich immunoassay through the hybridization of cAb_oligo_ conjugates onto BMPs for protein encoding without affecting fluorescence intensity barcodes. Our construct demonstrated improved sensitivity for the sandwich immunoassay of recombinant IFN-γ compared to the previously reported cAb_biotin_ construct. The use of biotinylated dAb and labeled SA to generate assay signal now leverages the use of commercially optimized antibody pairs (e.g. ELISA), facilitating protein panel development. Finally, the efficient use of a single fluorescence channel dedicated to barcoding enables the allocation of other channels for different purposes (e.g. cell labeling).

### Droplet microfluidic generation of permeable alginate hydrogels/PiPs by competitive ligand exchange

To perform MSA at the single cell level, individual cells must be isolated in an enclosed compartments together with multiple BMPs, allowing the secreted cytokines to be captured by the cAb-functionalized BMPs. Additionally, the spatial association between cells and BMPs must be maintained throughout the assay to preserve single cell resolution. Common single cell immunoassays isolate cells in W/O droplets in presence of all assay reagents (e.g., labeled dAbs). This negatively impacts assay performance due to high background signals. In addition, the enclosed confinement of droplets can adversely affect cell viability.

To overcome these issues while preserving cell-BMPs association, we applied W/O droplets to template alginate hydrogels, which upon resuspension in buffer, acted as permeable scaffolds for supplying and removing assay reagents. First, we generated 50 μm (∼65 pL volume) W/O droplets containing soluble alginate (1% w/v) using a flow-focusing microfluidic droplet generator (Figure S15).^[34]^ We selected alginate for its optical transparency and ease of gelation by ionic cross-linking in the presence of a divalent cation. To ensure homogeneous cross-linking, we used a competitive ligand exchange crosslinking process employing Ca-EDTA and Zn-EDDA (Figure S15).^[35]^ Briefly, upon co-encapsulation of Ca-EDTA/alginate and Zn-EDDA/alginate solutions at pH 7.0, the higher affinity of EDTA toward Zn^2+^ ions induces the complexation of Zn^2+^ by EDTA and the concomitant release of Ca^2+^ ions within the droplets. Since alginate has a higher affinity for Ca^2+^ ions compared to EDDA, alginate is ionically cross-linked into hydrogels templated by the current droplet geometry. Since our CLSM lacks a differential interference contrast, localization of transparent alginate particles relied on fluorescence. However, directly labelling the alginate would interfere with imaging and segmentation of cells and BMPs within the PIP, as well as with barcode decoding and quantification of fluorescent binding signals. Hence, we chose to fluorescently dope the surrounding medium instead, utilizing size-exclusion to restrict diffusion of the fluorescent dye into the alginate particles, which then appeared as dark spheres. We identified FITC-labeled dextran with a molecular weight >2MDa as effectively excluded from diffusing into the hydrogel (Figure S16). This approach allowed us to clearly recognize alginate PiPs, image within them, and automatically segment the alginate particles, cells, and BMPs, while accurately decoding and quantifying assay fluorescence on the BMPs.

To assess whether alginate particles remained permeable to assay reagents (e.g., dAbs and labeled SA), we encapsulated antibody-coated MPs in droplets with soluble alginate to generate PiPs. We immobilized the PiPs in a microchannel and perfused them with AF488-labeled dAb and BV421-labeled SA (Supplementary Note 3; Figure S17A). Biotinylated antibodies on the surface of MPs allowed direct binding of SA, while the dAb, a chicken anti-goat Ab, recognized the goat anti-mouse Ab on the MPs (Figure S17B). Upon constant flow of AF488-dAb and BV421-SA, we observed selective binding of both dAb and SA on the MPs with minimal background within the hydrogels (Figure S17C-D). We calculated a characteristic diffusion time *t* of 3.6 and 10.1 min for AF488-dAb and BV421-SA, respectively, in concordance with their respective molecular weights of ∼150 kDa and 340 kDA (Figure S17E). These results validate the selective permeability of alginate/PiPs for immunoassay reagents, but not for molecules with ≥ 2MDa.

### Probabilities and statistics of multiple BMPs encapsulation in PiPs and multiple PiPs in droplets

Following the generation of PiPs, we evaluated the probability of assaying our complete cytokine panel within individual PiPs for downstream single cell MSA. We encapsulated our seven BMPs into PiPs by droplet-based microfluidics at a concentration of 30 000 BMPs μL^-1^ per barcode (total of 210 000 BMPs μL^-1^; Figure S18A). While the Poisson statistics predict ∼14 BMPs (λ = 13.85) on average per PiP (Figure S18B), we measured on average 12 BMPs per PiP experimentally. This discrepancy between prediction and experimental results was attributed to aggregation and sedimentation of BMPs over time during the encapsulation process, skewing the encapsulation rate.

Considering the stochastic nature of BMPs encapsulation, one cannot expect to encapsulate only one of each of the seven target-specific BMPs used in our seven-plex assay. However, since we encapsulated ∼12 BMPs per PiP, we found that on average ∼35 % of PiPs contained the complete seven-plex BMPs (Figure S18C). We also calculated the probability of co-encapsulating multiple BMPs with the same barcode (i.e., encoded antibody, corresponding to a technical replicate) in a PiP. By considering the concentration of single barcode (30 000 BMPs μL^-1^) for prediction, our results were in good agreement with Poisson statistics, with the probabilities of encapsulating 0, 1, 2 or 3 replicates of the same barcode being 22, 25, 19 and 6 %, respectively (Figure S18D).

To perform MSA at the single cell level, the generated PiPs need to support downstream manipulation such as multiple cycles of centrifugation and resuspension without significant degradation. Hence, PiPs, which are loaded with ∼12 BMPs each consisting of a dense paramagnetic bead, needed to exhibit sufficient mechanical stability. We observed that PiPs formulated at pH 7.0 offered optimal mechanical stability, aligned with previously reported results by Bassett *et al*.^[36]^ In an attempt to minimize PiPs manipulation, we assessed if the pH and optimal alginate cross-linking conditions (42 mM Ca-EDTA/Zn-EDDA, in MOPS buffer) would enable direct immunoassay without loss in assay performance. We supplemented 100 ng mL^-1^ of recombinant IL-17A along with IL-17A encoded BMPs during PiPs generation by droplet-based microfluidics, and kept the produced hydrogels in their templating droplets. Here, target capture and alginate gelation would occur simultaneously within the same droplet.

Assay results showed a reduction in signal that we attributed to reduced capture efficiency of IL-17A by the cAb-BMPs at pH 7.0. However, when formulated at pH 7.4 to maximize the affinity of the cAb-BMPs, the PiPs showed an important reduction in mechanical stability and failed to sustain consecutive cycles of centrifugation and resuspension. To improve assay signal and preserve PiPs integrity, we formulated PiPs at pH 7.0, and adjusted the pH to 7.4 prior performing protein capture (Figure S19). We re-entrapped PiPs into *de novo* W/O droplets, similar to templated emulsion^[37]^, to maintain single cell resolution during antigen capture. To entrap PiPs in droplets, the washed PiPs were concentrated by centrifugation, and re-emulsified into W/O droplets by adding fluorinated oil supplemented with fluorosurfactant and rapidly pipetting up and down (Figure S20A). The concentrated aqueous suspension of PiPs is completely dispersed into the fluorinated oil due to their immiscibility. The emulsion remained stable due to the presence of fluorosurfactant that prevented droplet coalescence. CLSM imaging confirmed that most droplets contained a single PiP (Figure S20B), with the droplet diameter matching the size of a single PiP of ∼50 μm diameter (Figure S20C). Larger droplets showed the concomitant encapsulation of multiple PiPs per droplet (Figure S20C). Overall, more than 90 % of droplets contained a single PiP, minimizing potential crosstalk between PiPs in downstream single cell assays (Figure S20D). We also measured negligible leakage of biomolecules between mixed W/O droplets following production, or after 24 h incubation at 37 °C. Approximately 1 in 2000 droplets exhibited leakage (Figure S21), consistent with the leakage observed in droplet-based RNA sequencing technologies.^[38]^ Our results confirm that PiPs generated at pH 7.0 can be manipulated without risking to rupture the hydrogel. Moreover, PiPs can be effectively re-encapsulated into droplets, where droplets had negligeable crosstalk in terms of leakage and exhibited >90% yield of encapsulation of single PiPs per droplet. These results showcase the potential of PiPs to conserve single cell information.

### PiP-plex versus bulk BMPs

To assess the impact of the alginate hydrogel on sandwich immunoassay performance when using PiP, we performed MSA by incubating separately PiPs and pooled cAb-BMPs with a mixture of antigens in buffer, followed by a cocktail of dAbs, and BV421-labeled SA. For MSA in bulk, pooled cAb-BMPs were first immobilized electrostatically using a PLL-coated well plate prior to incubation with BV421-SA. We observed that the addition of SA led to aggregation of cAb-BMPs in bulk, in an antibody dependent manner, which impaired overall imaging and decoding of BMPs. As reported earlier, we attributed this phenomenon to the valency of the biotinylation of the antibodies. Immobilization of BMPs prior to labeling by SA eliminated this issue, which was not needed with PiPs whom intrinsically trap BMPs in 3D.

Depending on the antigen, PiPs exhibited slightly improved LOD compared to pooled cAb-BMPs in buffer. For instance, in the case of IL-4, a 50-fold improvement was observed, while for MIP-1α a 4-fold decrease was obtained (Figure 4). Since the LOD was calculated as three times the standard deviation of the blank above the background signal, reducing assay variability in blank samples directly improves the LOD. In our MSA, we obtained %CV of 101 % for the blank samples using bulk BMPs compared to 39 % when using PiPs (Figure S22). At low concentration, the assays showed higher variation between replicates. The %CV was reduced to <10 % at concentrations above 100 pg mL^-1^ (Figure S22). The LODs of PiPs ranged from 0.8 to 2000 pg mL^-1^ on PiPs and from 0.4 to 1000 pg mL^-1^ on cAb-BMPs. Although there was an improvement in LODs, we observed a reduced slope (reduction in the steepness) of the binding curves (i.e., a decrease in analytical sensitivity) for MSA performed in PiPs compared to bulk, which could be useful for single cell assays. The reduced slope led to an increase in the dynamic range of all cytokines assayed in PiPs compared to bulk BMPs. A lower slope can be beneficial in single cell assays, where diluting samples to fit within the assay’s narrow dynamic range is not practical. Consequently, a wider dynamic range enabled by a shallower slope allows for more accurate quantification across the diverse secretion levels inherent to cellular heterogeneity.

**Figure 4:**
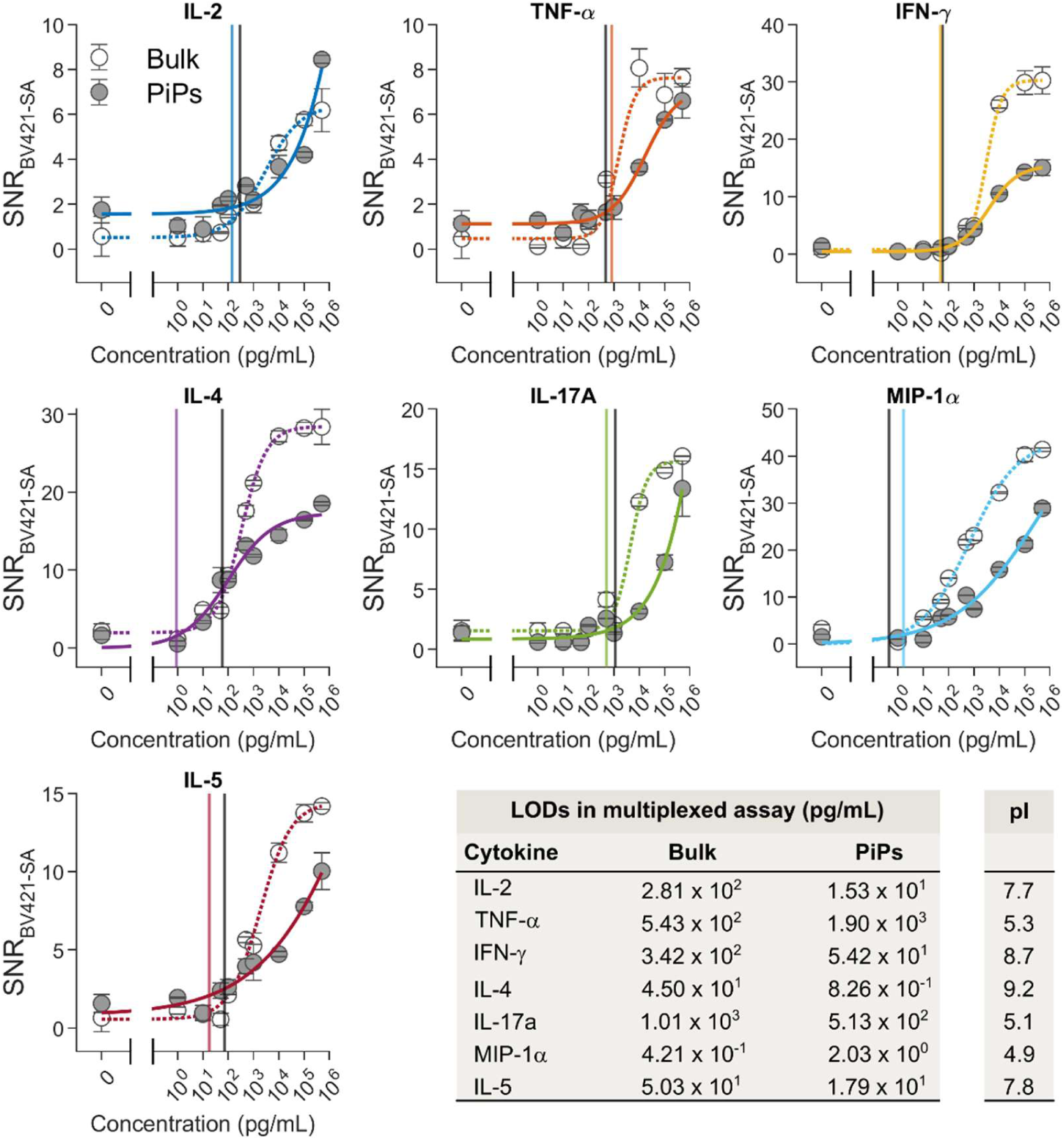
Multiplex assay binding curves acquired by CLSM for the 7-plex cytokine panel (IL-2, TNF-α, IFN-γ, IL-4, IL-17A, MIP-1α and IL-5) performed using the combined 7 cAb-BMPs in bulk (dashed lines) or formulated as PiPs (solid lines). Mean ± S.D. (*n_PIPs_* = 5*, n_bulk_ =* 5) are presented. The plots show a fit corresponding to a four-point logistic regression. In all cases, the pooled cAb-BMPs concentration was adjusted to 100 cAb-BMPs per microliter per samples. Calculated LODs for the different cytokines measured in bulk and PiPs with the corresponding isoelectric point (pI) are presented. Vertical lines indicate calculated LODs for the bulk cAb-BMPs (black lines) and PiPs (color lines).

At high concentrations of IFN-γ, IL-4, IL-5 and MIP-1α, a reduction in assay signals was observed in PiPs compared to bulk. While it is known that confinement within the hydrogel network can lead to improved protein association and increased specificity, the choice of hydrogel material is critical, as some materials may be prone to non-specific protein binding due to electrostatic interactions or inherent binding affinities. We evaluated whether the isoelectric point of the assayed cytokines could account for improved or reduced assay performance in PiPs formulated with alginate compared to bulk BMPs, but found no generalized causal effect (Figure 4). IFN-γ, IL-4 and IL-5 possessing isoelectric points of 8.7, 9.2 and 7.8 respectively, showed the highest reduction in assay signal at high concentration rationalized by their excess of positive residue at pH 7.4 used for the assay. However, MIP-1α with an isoelectric point of 4.9, also showed reduction in assay signal.

From a physiological standpoint, cytokines are known to bind electrostatically to negatively charged glycosaminoglycan components in the extracellular matrix.^[39–41]^ In fact, IFN-γ,^[42]^ IL-4,^[43]^ IL-5^[44]^ and MIP-1α^[45]^ can all electrostatically bind to glycosaminoglycans. Alginate contains the negatively charged residue guluronate, an enantiomer to iduronate, which is an important constituent of glycosaminoglycan and central to the generation of their negatively charged clusters. Guluronate may reduce specific binding due in part to its different stereochemistry, but it still contributes to the generation of negatively charged clusters, which are associated to the non-specific binding of cytokines to alginate and the elevated background, ultimately reducing the measured SNRs at high protein concentrations. Despite this, alginate still provides key advantages compared to other materials such as ease of gelation, potential for simple chemical modification, ease of degradation through enzymatic reaction or scavenging of cross-linking ions for cell retrieval, and enhanced MSA performance via confinement and molecular crowding. Overall, the calculated LODs are compatible with protein expression levels of single cells, with LOD <1 ng mL^-1^,^[15]^ while empowering multiplexing.

### PiP-plex of cytokines secreted by single macrophages-like cells

We tested PiP-plex with the well-established THP-1 human leukemia monocytes commonly used to test single cell proteomic platforms.^[2]^ First, THP-1 cells were differentiated into macrophage-like (Mφ) cells by 48 h incubation with 25 nM Phorbol 12-myristate 13-acetate (PMA).^[46]^ Differentiation was confirmed by the expected change in morphology and transition from suspended to adherent cells (Figure S23). Mφ THP-1 cells were encapsulated with BMPs using droplet-based microfluidics as described above to form PiPs. Cell concentration was selected to maximize single cell encapsulation based on Poisson distribution, with ∼45 % of PiPs containing a single cell (Figure S24). Using calcein-AM staining, we documented >95 % cell viability (Figure S25), which decreased to 90 % upon re-emulsification and incubation between 3 – 6 h. We further observed significant cell death for long term incubation in W/O droplet with viability decreasing to ∼30 % after 18 h, which can be attributed to the depletion of nutrients and accumulation of waste (Figure S25). We selected 6 h as the optimal incubation time to balance maximizing protein secretion while maintaining cell viability. It is noteworthy that dead cell stains, such as ethidium homodimer-1, which label available double stranded DNA, homogeneously labeled the BMPs and affected the measurement of the signal in the 561/590 nm channel. This homogeneous staining did not significantly affect decoding of the BMPs, but led to an increase in %CV (data not shown). Hence, we did not stain dead cells for further analysis, and we focused solely on viable cells.

Having confirmed that cells remained viable in PiPs, subsequently we assessed whether they could be recovered from the alginate hydrogels. THP-1 cells were co-encapsulated with BMPs in alginate-based PiPs, and the hydrogel was subsequently dissolved using a mixture of trypsin-EDTA and 2% w/v citrate to competitively chelate Ca^2+^ ions responsible for ionic crosslinking of the alginate.^[47]^ Following dissolution of alginate, BMPs were removed by magnetic separation, and cells were collected by centrifugation (Figure S26A). This process preserved >95% cell viability (Figure S26B-C), demonstrating that cells can be efficiently and gently retrieved from alginate-based PiPs.

We next evaluated the cytokine secretion of single Mφ THP-1 cells following LPS stimulation known to stimulate cytokine secretion by activating Toll-like receptor 4 (TLR4) signaling.^[48]^ Firstly, Mφ THP-1 cells were co-encapsulated with cAb-BMPs in PiPs, washed and resuspended in medium supplemented with 10% FBS, 1% Pen/Strep, 25 mM HEPES and 2 mM CaCl_2,_ for both pH equilibration (pH 7.4) and nutrient supply. Next, we exposed PiPs to 100 ng mL^-1^ LPS and re-encapsulated them into W/O droplet to stimulate protein secretion and enable cytokine capture by the cAb-BMPs. Following incubation, we assayed the captured cytokines by MSA and used CLSM for assay readout. Here, PiPs were loaded into an imaging well plate and localized by supplementing 2MDa FITC-dextran to the surrounding medium, which acted as a negative stain to enhance the contrast of the PiPs’ boundary, while the cells were stained with calcein-AM (Figure 5A) allowing precise determination of their centroid using the resulting fluorescence signal. Since both FITC and calcein are imaged using the same Ex/Em condition, we purposely oversaturated calcein signals generated inside the cells’ cytosol and applied variable thresholding on *Z*-projected images to sequentially segment PiPs and cells (Figure 5B). Using image analysis, we extracted the contour of each PiPs using a low pass threshold (Figure 5C; PiP segmentation) and extracted the centroid of each cell with a high pass threshold (Figure 5C; Cell segmentation). We used the *XY* coordinate of each cell centroid to assess if a cell was entrapped in a hydrogel using the extracted boundaries of PiPs. This step allowed the identification of cell-loaded PiPs (Figure 5C; Cell-loaded PiPs). Similarly, BMPs were segmented based on their scattering signals (561/590 nm) and assigned to their corresponding PiP by comparing their *XY* coordinates to the hydrogel boundaries (Figure 5C; BMP association). BMPs and cells whose *XY* coordinates fell outside the boundaries of a PiP were excluded from further analysis. Retained BMPs were decoded based on their intensity barcode in the AF647 channel, while proteins secreted from a single cell were measured via the signals generated by the BV421 labeled SA (Figure 5D).

**Figure 5:**
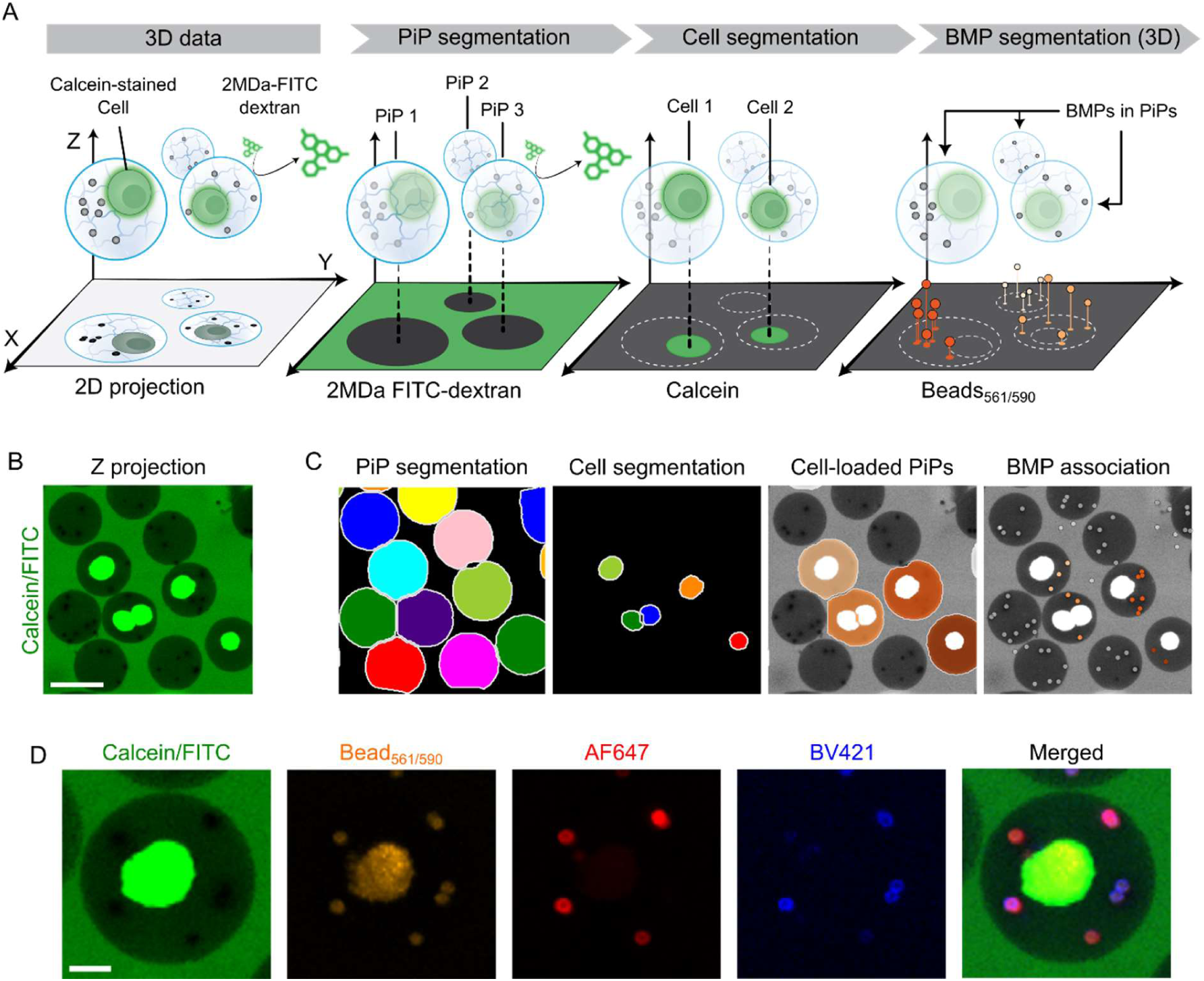
Segmentation of PiPs, viable cells and BMPs, followed by fluorescence quantification of barcodes and binding signals for PiP-plex assays. (**A**) Schematic overview of PiP segmentation and BMP association. PiPs immobilized on a poly-lysine coated coverslip and encapsulated cells are sequentially segmented from the Z-projection using a single fluorescence channel (FITC/Calcein). BMPs are segmented directly in 3D space based on their scattering signal at 590 nm, and associated with a specific PiP through their *XY* coordinates and PiP contour. (**B**) Z-projection image used for segmentation. Viable THP-1 cells are labeled by calcein-AM, which diffuses through the PiPs and across the cell membrane, while a high molecular weight FITC-labeled dextran (2MDa) is excluded from the PiPs, enhancing the signal of the surrounding buffer. Scale bar: 50 μm. (**C**) Segmentation of image (B) with superposed masks. PiPs are initially segmented by applying a low pass threshold, followed by cell segmentation with a high pass threshold. Cell-loaded PiPs are identified by comparing the cell’s coordinate to the PiPs’ contour. BMPs are then associated with a specific PiP based on their coordinates. (**D**) CLSM images of a single PiP at different fluorescence channels. The Calcein/FITC channel was used to locate the cell and the PiP. The 561/590 channel was used for BMP segmentation. The AF647 fluorescence was used to assess intensity barcodes and associate them with specific proteins. Finally, BV421 fluorescence channel was used to record the binding signal. Scale bar: 10 μm.

Using our image processing pipeline, we identified the number of BMPs entrapped within each PiP, along with their corresponding encoded cytokine. We focused our analysis on PiPs containing at least 5 distinct BMPs. To avoid bias in multivariate analysis, missing protein measurements (due to the absence of their corresponding BMPs) were imputed. Specifically, missing values were replaced with the average signal for that protein, calculated from all BMPs of the same type within samples under the same experimental condition. For PiPs containing multiple BMPs targeting the same protein, we summed the BV421 fluorescence signals to account for the distribution of captured proteins across these replicate beads. Assay signals were then background corrected by subtracting the average signal of the corresponding barcoded BMPs exposed to the same assay conditions in the absence of cytokine. We believe that this approach more accurately reflects the concentration of secreted protein at the single cell level, as it accounts for signal dilution across multiple beads. In contrast, failing to consider this dilution could lead to an underestimation of protein levels. Conversely, if a protein-encoded BMP was absent from a PiP (e.g. BMP not encapsulated), that protein could not be assayed.

The microfluidic traps designed to immobilize PiPs in a microchannel with a step height reduction used in permeability testing (see Supplementary Note 3, Figure S17) were not employed for cell assays. Instead, we opted for an open well strategy for imaging, which unlike our current traps, allowed PIPs to arrange in a monolayer format without clogging. As such, spatial overlap of PiPs was minimized, allowing accurate co-localization of cells and BMPs in PiPs using the Z-projection of the FITC-dextran emission channel. In addition, the use of commercial imaging well-plate facilitated platform dissemination and simplified sample manipulations.

Using PiP-plex, we measured an increased secretion of MIP-1α, TNF-α, IL-17A and IL-2 in Mφ THP-1 cells stimulated by LPS compared to the unstimulated controls (Figure 6A) which is in good agreement with secretion profile reported by Ma et al.^[2]^ We validated our findings by bulk measurements of cytokines in the supernatant from LPS-stimulated and unstimulated control cells cultured in well plates in the absence of alginate. The measurements were performed using antibody microarrays with the same antibody pairs employed for PiP-plex analysis. While perfect consistency is not expected due to differences in assay formats (i.e., single cells vs. bulk measurements, different LOD cut-offs, potential statistical sampling biases, and different materials and their impact on cytokine secretion), we anticipate replicating key findings. For both PiP-plex and antibody microarrays, only assay signals surpassing the LOD for each cytokine were considered in the assessment of statistical significance. Both platforms detected highly significant differences in secretion of MIP-1α and TNF-α following LPS stimulation (Figure S27-28 vs Figure 6A). IL-17A was also highly differentially secreted when assayed in PiP-plex, but since it did not reach the LOD on the microarray, the secretion differences could not be measured. Interestingly, IL-5 exhibited significant differential secretion in bulk, but not with PiP-plex (Figure 6A, Figure S27), possibly because most single cell measurements were below the LOD. For IFN-γ secretion, which was detectable in the PiP-plex assay but did not show significant differences, values above the LOD could not be measured in the bulk assays.

**Figure 6:**
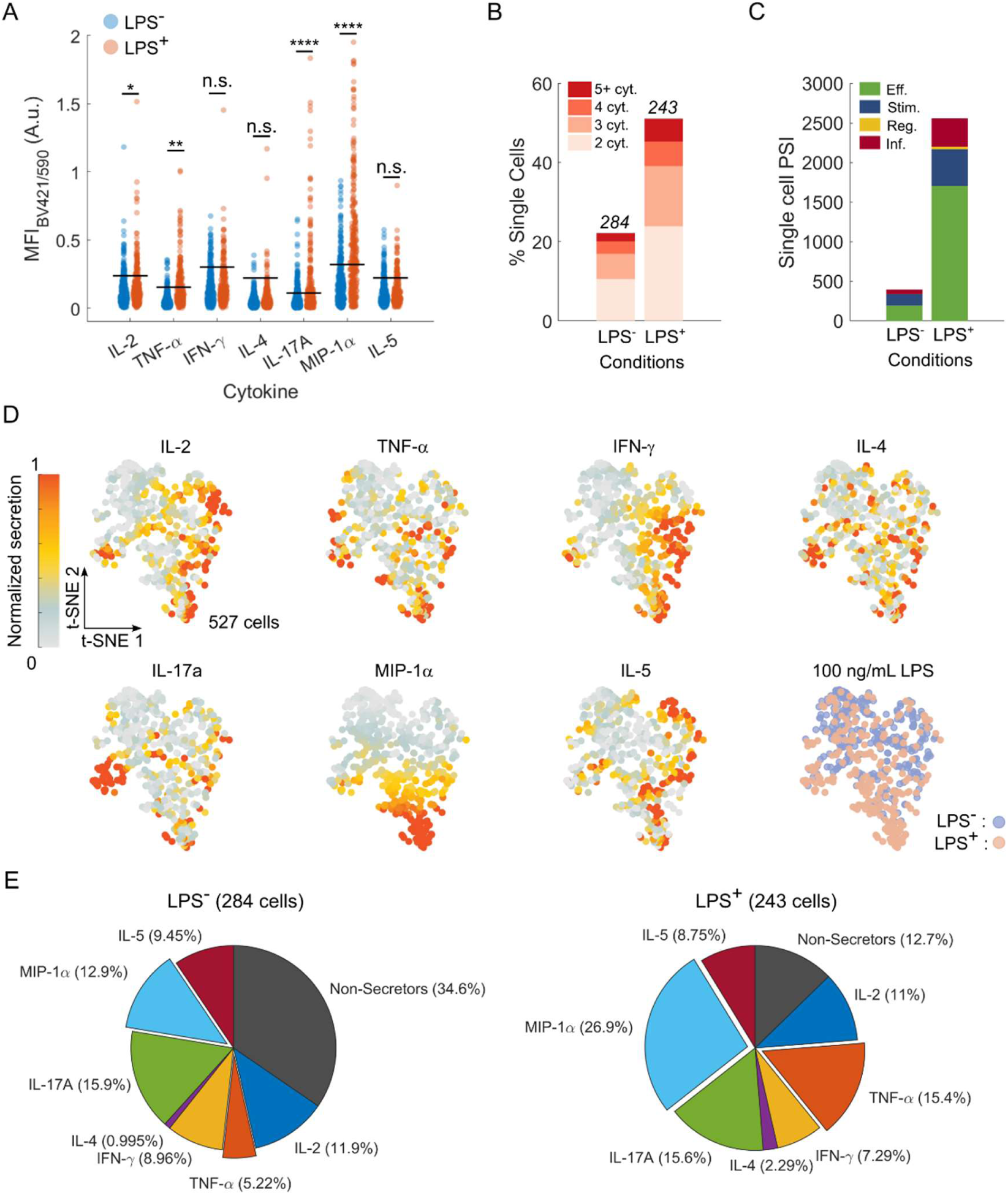
Single cell quantification of secreted protein by THP-1 Mφ cells via PiP-plex. (**A**) Scatter plots showing BMP signal measurements from THP-1 Mφ cells in PiPs upon stimulation with 100 ng mL^-1^ LPS. ns: not significant; * *P* < 0.05, ** *P* < 0.01, **** *P* < 0.0001 by Mann-Whitney two-tailed test. Black lines represent the assay’s LOD, calculated as three times the standard deviation above the background of PiPs containing no cells. Comparison of cytokine secretion of non-stimulated and stimulated THP-1 Mφ cells represented in the form of (**B**) single cell polyfunctionality shown as a proportion (%) of cells that secreted 2, 3, 4 or more than 5 cytokines above the assay LOD for the cells entrapped in PiPs containing at least 5 different cAbs-BMPs. The values above the bar correspond to the total amount of cells analyzed per condition, including non-secretors and single-cytokine secreting cells. (**C**) Polyfunctional strength index (PSI, with Eff: Effector; Stim: Stimulatory; Reg: Regulatory; Inf: Inflammatory) from cells described in (**B**). (**D**) T-distributed stochastic neighbor embedding (t-SNE) of polyfunctional cells (527 cells) according to their secretion of the seven cytokines and (**E**) the distribution of cytokine secretions with exploded views of TNF-α and MIP-1α to highlight the fold changes in secreting cell populations.

We evaluated whether the random spatial distribution of BMPs entrapped within alginate affects the capture efficiency of secreted proteins and impacts assay performance. Our results showed that the average distance between BMPs and cells was ∼7.5 ± 5.8 μm (Figure S29) with 95% of the BMPs located within 19 μm from a cell. Among all BMPs, 4.1% of exhibited above the average LOD across all cytokines (MFI_BV421/590_ of 0.22), while only 0.3% of these signal-positive BMPs were located farther than 19 μm from a cell. These findings suggest that assay signal intensity is largely independent of the cell-to-BMPs distance, as the vast majority of BMPs reside in close proximity to their associated cells. Based on our diffusion studies (Figure S17), the random spatial distribution of BMPs within PiPs does not appear to affect the binding of dAbs or SA, as evidenced by the low signal variance across all BMPs.

We assessed the polyfunctionality of 527 viable THP-1 cells by enumerating cells that secreted more than one cytokine above the assay LOD (Figure 6B). Without LPS stimulation, approximately 10 %, 6 %, 3 % and 2 % of 284 cells secreted 2, 3, 4 or more than 5 different cytokines, respectively. In contrast, upon LPS stimulation approximately 23 %, 15 %, 6 % and 5 % of the 243 cells secreted 2, 3, 4 or more than 5 different cytokines, respectively. In both cases, cytokines are part of various immunological programs, acting as effector, regulatory, stimulatory or inflammatory molecules. Our results demonstrate that following LPS stimulation, a larger proportion of cells were engaged in different immunological programs and secreted cytokines above the assay LOD, as quantified by the polyfunctional strength index (PSI; Figure 6C). Alongside cellular polyfunctionality, we evaluated the proportion of non-secreting cells and those secreting a single cytokine, based on whether the assay signal was below or above LOD, respectively. We noted an increase in the number of secreting cells upon stimulation (Figure 6D). Specifically, TNF-α and MIP-1α secretion increased 2- to 3-fold following LPS treatment, while the proportion of cells secreting other cytokines remained unchanged. For cells secreting at least 2 distinct cytokines, the cytokine contribution and individual profiles showed notable differences upon LPS stimulation compared to control (Figure S30). Secretion profiles indicated both a greater number of polyfunctional cells and an increase in the number of cytokines secreted by the LPS-treated cells, which was reflected as an increase in PSI. Overall, PiP-plex enabled robust assessment of cellular polyfunctionality, while retaining cellular viability throughout the immunoassay and downstream recovery.

## Discussion

We introduced a novel strategy to encode and anchor cAbs onto BMPs using dsDNA. By leveraging dsDNA as a modular building block, we generated up to seven distinct single-color BMPs with our novel construct. This was enabled by the direct transfer of dsDNA concentrations in solutions onto BMP surface and the elimination of binding competition among biotinylated reagents (Figure 3). Subsequent anchoring of DNA-antibody conjugates enabled downstream sandwich immunoassays. We observed an enhanced protein binding efficiency which we ascribe to the immobilization of antibodies via a DNA-linker that provided flexibility for gyration, a more favourable orientation, and reduced steric hindrance for protein capture compared to physisorbed antibodies. Our strategy accommodated various antibody panels for tailored immunoassays utilizing commercially available and validated antibody pairs for ELISA. Our novel construct also supported the use of various fluorescent dyes for assay signal generation, including small organic and large polymeric dyes such as Brilliant Violet® which are readily available in SA conjugate format. The synthesis of cAb-BMPs strictly relied on affinity interactions between hybridized DNA strands, and biotin-streptavidin binding. While this approach was sufficient for our single cell immunoassay, we noted an increase in %CV in barcode intensity (reflected as broader intensity distribution) when BMPs were pooled and stored for more than 7 days (data not shown). This signal broadening was attributed to gradual de-hybridization of DNA strands from the bead surface over time. We thus limited the use of cAb-BMPs to within one week of preparation. Another important consideration is the susceptibility of cAb-BMPs to degradation by DNases. This is particularly relevant for single cell assay that may require long incubation times with live, biologically active cells. We did not observe any degradation of the cAb-BMPs (e.g. spreading of barcode intensity) over the course of our experiments (∼6 h). We hypothesize that the use of hydrogels, which restrict cellular motility and spatially immobilize both cells and cAb-BMPs, may minimize undesired cell-bead interactions and signaling events, such as those from macrophage-like THP-1 cells, that could otherwise trigger DNase activity.

PiP-plex is based on the entrapment of cells and BMPs within hydrogel particles, enabling spatially indexed multiplexed immunoassay in 3D. We developed an image processing pipeline enabling the segmentation of individual BMPs in 3D by CLSM which emulated single particle detection acquired by FCM. Our approach proved to be successful for both immobilized BMPs onto coverslips and BMPs formulated into 50-μm-diameter PiPs. Our pipeline circumvented the limitations in *Z-axis* resolution of FCM while the CLSM provided 3D images of each PiP including the encapsulated cells, BMPs and quantitative fluorescence signals. Our multiplexing capability surpasses other well established single cell secretomic platforms based on optofluidic technology^[49, 50^], and confined microenvironment such as droplets^[9, 11^] or nanovials^[26]^ by 2- to 3-fold, all while maintaining cellular viability and supporting cell retrieval. Multiplexing could be further increased in PiP-plex by introducing additional fluorophores (i.e. dsEO_i_ ) for barcoding, such as ATTO750, as previously developed.^[29]^ However, the formulation of PiPs used here follows Poisson statistics, meaning that as multiplexing increases, the probability of generating PiPs with the full panel of cAb-BMPs decreases. PiP-plex is not inherently limited to DNA-based beads or to alginate as the hydrogel material; it could, in principle, be applied to other barcoded beads platforms, such as xMAP beads from Luminex. However, our BMP construct enables a rapid and flexible tuning of both barcoding and detection fluorophores to maximize compatibility with available microscopes, while xMAP is locked into red and far-red fluorophores. Conversely, our BMPs are compatible with most standard microscope setups by employing commercially available dyes covalently linked to DNA. Moreover, the smaller size of our BMPs (∼2.8 μm) allows for a higher theoretical bead loading per 50 μm hydrogel particle compared to the xMAP beads, which typically range from ∼5.6 to 6.5 μm in diameter.

PiP-plex achieved an experimental throughput of ∼1000 cell h^-1^, rivaling other microfluidic chip-based methods^[2]^, but using a hydrogel PiP instead of a solid-state chip. We analyzed 1441 cells in total and ∼6000 BMPs in less than 90 minutes. Currently, this throughput is limited by the acquisition rate of our microscope and unfavorable Poisson statistics. For instance, we acquire 4-color *Z*-stack images (320 x 320 μm^2^) composed of 35 slices in approximately 25 s. On average, each of these *Z*-stacks comprises 14 PiPs, whereas 6 of them are containing a single cell. In addition, ∼35 % of imaged PiPs contain the 7-plex panel. Taken together, there is 2 PiPs on average per field of view that encapsulate both a single cell and the full 7-plex panel. Overall, these statistics led to a total of 527 cells to be considered for downstream analysis by our analytical pipeline. Further improvements are possible. Encapsulation approaches using inertial microfluidics that surpass Poisson statics could improve the odds of simultaneous encapsulation of the full 7-plex panel with a single cell.^[51, 52^] Compression traps that squeeze PiPs into thin hydrogel layers could increase imaging speed by reducing the number of Z-stacks.

Overall, the use of alginate as a hydrogel for our PiPs enhanced MSA performance of PiP-plex resulting in better LODs and wider dynamic range compared to assays performed with bulk cAb-BMPs (Figure 4). This improvement was attributed to the confinement effect of the hydrogel that increased both the apparent local protein concentrations and the collision frequency between molecules, as discussed by Moorthy *et al*.^[53]^ Moreover, confinement favors protein association over dissociation due to favorable entropic contribution via molecular crowding.^[54]^ We speculate that alginate, a natural polysaccharide, can bind some cytokines, due in part, but not exclusively, to its intrinsic isoelectric point. Cytokine binding to alginate was more pronounced for IFN-γ, IL-4, MIP-1α and IL-5, which are all known for binding to glycosaminoglycan in the extracellular matrix. This rationale is consistent with the increased assay background and lower assay sensitivity (as defined by the slope of the curve in its linear region) of PiPs compared to bulk cAb-BMPs (Figure 4). Since assay signals were corrected by subtracting local background signals near each BMPs, an effective reduction in SNR was observed for PiP-plex compared to bulk BMPs. To further improve PiP-plex assay performance, sacrificial proteins, which do not affect cellular secretion, could be employed to passivate alginate and reduce undesired cytokine binding. Although the concept of PiPs entrapping beads in 3D for multiplexed single cell immunoassays within a hydrogel particle is powerful, the use of natural biopolymers to form the hydrogel matrix can inherently influence cellular secretion. Alternatives hydrogels to alginate include polyethylene glycol and agarose (widely used for single cell assays^[8, 55^]), but at the expense of live cell retrieval for expansion and further analysis. Agarose, while offering high mechanical stability and excellent diffusivity for antibody and proteins, was also reported to affect cytokine secretion.^[56]^ Nonetheless, different hydrogel composition could be considered for PiP-plex depending on the application, but careful validation of assay performance and antibody pairs, particularly in the context of MSA of secreted proteins at the single cell level, where protein concentrations are low and require antibodies with high affinity, specificity and minimal cross-reactivity will be required ^[33]^ as well as consideration of hydrogel-cell interactions.

As a proof-of-concept, PiP-plex was used to quantify proteins secreted by differentiated THP-1 cells (Figure 6). Since cytokine capture occurs within the alginate network, we confirmed that the spatial heterogeneity of BMPs, which are randomly distributed in 3D, did not affect the capture efficiency of cytokines secreted by single cell (Figure S29). We attributed this to the high confinement provided by the ∼65 pL droplets during incubation, which likely minimized the formation of concentration gradients. In contrast to previously reported microfluidic chips employing ∼3 nL chambers^[1, 2^], where signal variability was observed due to such gradients, PiP-plex appeared largely unaffected, consistent with a nearly 50-fold reduction in assay volume. Using PiP-plex, we measured an increased secretion of TNF-α, IL-17A, IL-2 and MIP-1α upon LPS stimulation compared to the unstimulated control (Figure 6A). We noted a good agreement between single cells (PiP-plex) vs bulk (antibody microarrays) measurements for the proteins that showed high differences in secretion and were well-above the LOD on both platforms (e.g. TNF-α and MIP-1α). IL-17A was highly secreted as measured by PiP-plex, but below the LOD on antibody microarrays, while IL-5 was only differentially secreted when measured by microarrays. We believe that the confinement in a 65 pL volume W/O droplet helped increase the local concentration of IL-17A, exceeding the LOD in PiP-plex. Alginate saccharides were reported to suppress expression of IL-5 and IL-4 associated factor in T helper 1, and 2 cells,^[57]^ which could account for the absence of significance in IL-5 signal in PiP-plex compared to the antibody microarray. Besides the non-specific binding of cytokines mentioned above, alginate is also known to induce immunological responses on cells via TLR4^[58]^ which is also activated by LPS. Furthermore, Ca^2+^-crosslinked alginate hydrogels can leach Ca^2+^ ions, which may trigger an inflammatory response from immune cells, as reported by Chan *et al*. in dendritic cells.^[59]^ These inherent aspects of alginate could be responsible for the activation of LPS^-^cells and increased the basal cytokine secretion that we observed for our untreated control. Basal cell activation could be optimized in the future by adjusting the alginate composition or gelation conditions. In the future, chemical modification of alginate backbone with immunomodulatory peptide could be explored.^[60]^

We assessed the functional diversity of PMA-differentiated THP-1 cells using PiP-plex while preserving cell viability. We detected approximately 65 % of cells secreting at least one cytokine above the LOD in the LPS^-^sample, and around 88 % in the LPS^+^ sample (Figure 6E), proportions that are higher compared to those observed in other single cell platforms.^[26]^ This discrepancy may be due to an improved LOD of PiP-plex, an elevation of basal secretion by cell due to intrinsic stimulation by alginate, or a combination of both. We detected the expected changes in secreted cytokines (Figure 6, and Figure S27) from THP-1 cells upon LPS treatment that vastly exceeded the threshold statistical significance, along with distinct t-SNE clustering of IL-17A and MIP-1α secreting cells. These findings both validate and illustrate the potential of PiP-plex for single cell assay. We also found that TNF-α was not secreted standalone, but rather co-secreted with other cytokines, concordant with the literature on LPS stimulated THP-1 cells.^[2, 5^] In addition, these three cytokines overlap with the cluster associated to the LPS^+^ condition. Likewise, cells secreting IL-2, IL-5, and IFN-γ were predominantly mapped to the right of the t-SNE graph, but failed to form well defined clusters, reflecting overall low secretion of these cytokines, and weak association with either LPS^-^ and LPS^+^ conditions.

One of the main limitations of PiP-plex for single cell assays stems from the challenge of encapsulating the complete 7-plex protein panel within a single hydrogel particle. As previously noted, we achieved the full 7-plex coverage at the single cell level for only ∼35% of imaged cells. To increase the number of analyzable data points per condition, PiP-plex can be adapted to include PiPs containing fewer distinct BMPs. In this study, we focused on PiP with at least 5 distinct BMPs, which required imputation of up to 2 missing cytokine signals, as these missing values do not necessarily indicate true negatives. Currently, we impute missing data using the average signal measured across all beads for the corresponding cytokine within the same sample. While this approach has its limitations, the data obtained by PiP-plex was in good agreement with bulk assays using antibody microarrays capable of quantifying all cytokines across samples. Importantly, PiP-plex preserves cell viability throughout the entire assay and allows straightforward cell recovery (Figure S26). This capability enables iterative single cell analyses of secreted proteins, either by probing different targets in sequential rounds using newly encoded BMPs, or by assessing functions such as cytotoxicity and killing efficiency through alternative assay. In future work, we aim to incorporate a microscopy-compatible technology^[61]^, that will allow the selective isolation of individual hydrogel particles based on their assay signal. Overall, we envision that PiP-plex could open multiple new investigative avenues, such as characterizing the secretome signature of various cells, studying the effect of extracellular ligand binding on the secretome by changing the composition of the hydrogel or even co-encapsulating more than one cell types per PiPs to investigate the effect of cell-cell contact and paracrine signaling effects on secretion.^[38]^ Finally, PiP-plex could empower all of these investigations while preserving cell viability, allowing retrieval of selected cells for further analysis and clonal expansion.

## Materials and Methods

### Materials

A complete list of biological and chemical reagents is available in the supporting information (SI, Table S1, S2 and S3). All DNA sequences were acquired directly modified from Integrated DNA Technologies (IDT) with HPLC purification. THP-1 cells were a gift from Prof. Nicole Li-Jessen (McGill University).

### Cell culture

Monocytes (THP-1, ATCC TIB-202^TM^) were cultured in T75 flask in RPMI-1640 medium (Gibco) supplemented with 2 mM glutamine, 10% heat inactivated fetal bovine serum (HI-FBS, Gibco) and 1% penicillin/streptomycin (Gibco), referred to as completed RPMI-1640 medium. Cells were cultured in suspension following manufacturer’s recommended protocol at 37 °C and 5% CO_2_ at 100% humidity. When needed, THP-1 cells at 1 x 10^6^ cells mL^-1^ were differentiated into macrophage-like (Mφ) cells by supplementing 25 nM (15 μg mL^-1^) of phorbol-12-myristate-13-acetate (PMA; Sigma Aldrich) in completed RPMI-1640 and incubated for a period of 48 h to allow differentiations. Medium was then exchanged for completed RPMI and cells were allowed to rest for 24 h. Differentiation of cells was confirmed by light microscopy (Axiovert 40 CFL, Zeiss), where differentiated cells became adherent and exhibited important morphological changes. Cells were cultured up to passage 10 for the current study.

### Preparation of the precursor solutions

1 M EDDA solution was prepared by dissolving EDDA in 100 μL of 1 M NaOH with sonication, while 1 M Zn(CH_3_CO_2_)_2_ solution was prepared in 100 μL of MilliQ water. Solutions were prepared fresh immediately prior to use. The resulting solutions were then mixed in a 1:1 ratio by vortexing and incubated for 5 min to allow the generation of the Zn-EDDA complex. Then, 42 μL of the Zn-EDDA solution was mixed with 20 μL of 1 M MOPS pH 7.0, 167 μL of alginate 3 wt% in MilliQ, 5 μL of 1% v/v Tween-20 in MilliQ and 224 μL of either completed RPMI-1640, or MilliQ water. The solution was vortexed, resulting in a final solution composed of 42 mM Zn-EDDA, 40 mM MOPS pH 7.0, 1 wt% alginate, 0.01% v/v Tween-20 in RPMI-1640 of MilliQ water. A 200 mM stock solution of Ca-EDTA was prepared by dissolving an equimolar amount of CaCl_2_ and EDTA in MilliQ water, and pH adjusted to 7.4 with concentrated NaOH.

### DNA Antibody Conjugation, Purification and Quantification

To generate the DNA conjugated capture antibody (cAb_oligo_), 200 μL (100 μg, 0.67 nmol) of 0.5 mg mL^-1^ antibody was buffer exchanged into pH 7.4 phosphate buffer saline (10 mM Na_2_HPO_4_, 1.8 mM KH_2_PO_4_, 137 mM NaCl, 2.7 mM KCl, PBS; Gibco) and concentrated to a volume of ∼20 μL (∼5 mg mL^-1^) using a 50k MWCO Amicon centrifugal filter (0.5 mL, Millipore Sigma). The antibody was diluted with 28 μL of PBS and mixed with 12 μL of a 1.67 mM Sulfo-SMCC (Thermo Fisher Scientific, USA) solution in 16.7% v/v DMSO in PBS, corresponding to a molar excess of 30:1 Sulfo-SMCC to antibody. Here, 10 mM stock solutions of Sulfo-SMCC were prepared in DMSO (aliquots stored at -20 °C) and diluted to 1.67 mM with PBS. The mixture was incubated for two hours at room temperature. Separately, 6 nmol of a 5’ thiol modified DNA (Thiol Modifier C6 S-S; IDT) was incubated in 100 mM DTT solution in PBS and for 1 h 30 at room temperature to reduce the terminated thiol. The reduced DNA was then desalted twice by using two Zeba spin columns (7K MWCO, Thermos Fisher). Separately, the antibody-SMCC conjugate was also desalted by using a single Zeba spin column. Then, the antibody-SMCC conjugate (75 μL) was mixed with the reduced thiol DNA (70 μL) and incubated for 1 h at room temperature followed by an overnight incubation at 4 °C. The resulting cAb_oligo_ was purified using a 50k MWCO centrifugal filter to remove excess of unreacted DNA. The final antibody concentration was determined with a Pierce^TM^ BCA protein assay (Thermo Fisher Scientific, USA) by adapting the *Micro-assay protocol* from the manufacturer’s instruction. Briefly, the volume of samples and reagents were reduced to 4 μL, mixed with 80 μL of assay reagent and incubated at 60 °C for 5 min. Bovine gamma globulin standards (Thermo Fisher Scientific, USA) were employed as standards for the assay. A typical protein recovery of ∼60-85 % was measured for the various conjugates. Finally, the cAb_oligo_ were diluted to 1 mg/mL in PBS without further purification and stored at 4 °C.

### Generation of the intensity barcoded microparticles (BMPs)

The generation of the barcoded microparticles (BMP) was based on the assembly of double stranded DNA on 2.8 μm diameter streptavidin-coated magnetic microparticles (DynaBeads M-270, Thermo Fisher Scientific, USA) as reported elsewhere.^[29]^ First, fluorescently labeled double stranded encoder oligo (dsEO*_i_*) were prepared by annealing 5’ biotinylated oligos with the 3’ labeled oligos with encoder dye *i* (EO_i_). EO_0_ was unlabeled, while EO_1_ – EO*_i_* were labeled with dye **1-*i***. To generate antibody anchored BMPs (cAb-BMPs), two different biotinylated oligos were used: a 21-nt space oligo (SO), and a 42-nt capture oligo (CO) which incorporate additional nts serving as an anchor for the cAb_oligo_. COs and SOs were separately annealed at 10 μM with individual EO_i_ at 80 °C for 5 minutes and cooled at 25 °C for 15 min in pH 7.4 PBS supplemented with 300 mM NaCl and 0.05% v/v Tween-20 (PBST0.05 + 300 NaCl), resulting in the generation of double stranded capture (dsCO*_i_*) and double stranded space (dsSO_i_) encoder oligos. Each dsCO_i_ was then mixed with their respective dsSO_i_ in a 2:7 ratio resulting in dsEOs solutions, one for each *i* dye. The proportion of dsCO_i_ and dsSO_i_ allowed fine tuning of the surface density of the cAb_oligo_. A 25 μL volume of a given barcode was prepared by mixing the various dsEO*_i_* at various proportions, such as dsEO_0_ + dsEO_1_ +…+ dsEO_i_ = 90 pmol, in PBST0.05 + 300 NaCl. Then, 25 μL containing 3.25 X 10^6^ streptavidin coated microparticles in PBST0.05 + 300 NaCl was added and mixed rapidly by pipetting up-and-down to initiate the co-capture of the biotinylated dsEO_i_ onto the beads surface. Solutions were incubated for 90 min at room temperature on an orbital shaker at 950 rpm to prevent sedimentation of the microparticles. The resulting BMPs were washed three times with pH 7.4 PBS supplemented with 0.1% v/v Tween-20 (PBST0.1) through cycles of magnetic aggregation and resuspension to remove uncaptured DNA. At this point, the BMPs may be kept, protected from light, at 4 °C until needed up to a period of one week.

### Immobilization of the DNA-antibody conjugate on BMPs

BMPs were rinsed thrice with pH 7.4 PBS supplemented with 0.05% v/v Tween-20 and 450 mM NaCl (PBST0.05 + 450 NaCl) and resuspend in a final volume of 50 μL. Then, 12.4 pmol of cAb_oligo_ in PBS were added to the BMPs (2:1 molar excess cAb_oligo_ to anchors on BMPs) and incubated for 30 min at room temperature on an orbital shaker at 950 rpm. After the pull-down of the cAb_oligo_, the fully assembled cAb-BMPs were washed thrice with PBST0.01 and stored at 4 °C until needed for up to a week. Prior to the assay, the BMPs were blocked with pH 7.4 PBS supplemented with 0.05% v/v Tween-20, 1% w/v bovine serum albumin (BSA; Jackson ImmunoReseach, PA) and 1 mM biotin (PBST0.05 + 1% BSA + 1 mM biotin) for 1 h (or overnight) on an orbital shaker at 950 rpm and rinsed three times with PBST0.1 prior the assay.

### Flow Cytometry and analysis

BMPs were detected using a ZE5 cell analyzer (Bio-rad, Ontario) with 488, 561 and 640 nm lasers employing 509/24, 577/15 and 670/30 band passes, respectively. Five thousand beads were measured per condition. Data were analyzed in MATLAB (v2022b, Mathworks) using custom-written codes. Briefly, single beads were segregated from doublets and other particles via their forward and side-scatter signal (FSC/SSC) and gated automatically by fitting a gaussian mixture model.

### Confocal imaging

Fluorescent images of the BMPs, immobilized on surfaces or inside alginate hydrogel (e.g. PiPs), were acquired using a Nikon eclipse Ti2 confocal microscope (Nikon, Japan) with a 20X (NA = 0.75) Plan Apo λ air objective (Nikon, Japan). The pinhole aperture was set to one 1.2 Airy Unit, and unless stated otherwise, the experiments were performed at room temperature. Digital offset of 10 (12 Bits images) was added to each fluorescent channel. For assaying by CLSM, *Z*-stacks, composed of 512 x 512 px^2^ images with a 2X digital zoom, were acquired with a *Z*-step of 0.42 µm. Imaging was performed using a resonant *(bidirectional)* scanning mode and averaged 8 times with an overall integration time of 517.2 ms per frame (dwell time: 1.97 µs pixel^-1^). Lasers were warmed up for at least one hour prior to the experiment to insure minimal laser fluctuation throughout the measurements. Four lasers (405/488/561/640) were used, along with 450/50, 525/50, 595/50 and 700/75 band pass, respectively. Channels were imaged pairwise to accelerate imaging by pairing 488/640 and 405/561 channels. These combinations provided minimal bleed-through. Imaging was performed via NIS-element AR (v5.42.04; Nikon, Japan).

### 3D-based image analysis

All data analysis was performed in MATLAB using custom-written code available elsewhere (https://github.com/orgs/junckerlab/repositories/DropletAssay). BMPs were segmented and analyzed directly on three-dimensional data acquired by CLSM by adapting a 3D Hough transform developed by Xie et al.^[31]^ available elsewhere (https://www.mathworks.com/matlabcentral/fileexchange/48219-spherical-hough-transform-for-3d-images). Briefly, the algorithm employs the spatial gradient to detect BMPs edges and estimate the individual bead center. Then, spherical masks, centered on each individual BMPs with a diameter of 3 μm, were generated and further segmented by a 3D watershed segmentation algorithm. The segmented masks were applied to the 3D images to integrate fluorescence intensity for all monitored Ex/Em channels. BMPs exhibited a strong signal emission in the 595/50 channel upon excitation at 561 nm, herein referred to as the Bead_590_ channel, acted as a proxy of the scattering of individual BMPs. By employing this pseudo-scattering signal at 590 nm, BMPs were intensity-gated by imposing both a minimal and maximal intensity threshold corresponding to 85 % and 115 % of the median intensity distribution measured at 590 nm. This gating strategy eliminated false BMPs (i.e., false positive corresponding to bright detected individual pixels) and clustered BMPs resulting in poorly positioned spherical masks. Following intensity gating, the intensity barcode of individual BMPs was identified by normalizing the AF647 measured signal by the pseudo-scattering Bead_590_ signal. Assay signal measured in BV421 channel were also normalized by the pseudo-scattering.

### In silico design of BMPs

Due to the capability to generate independent barcodes through the usage of biotinylated oligos, barcode position and its resulting spectral bandwidth may be predicted and hence guide the development of spectral barcode exhibiting minimal overlaps (see Supplementary Note 2). Further details can be found elsewhere.^[29]^ Briefly, four single-color BMPs were generated per fluorescent dye, with an increase proportion of dsEO_Dye_:dsEO_unlabeld_, and measured by CLSM and FCM as described above. These standards enabled the extraction of the irradiance μ_i_ of each dye, corresponding to the slope of the calibration curve, and the average %CV across all standards of the same dye. Using these two metrics, intensity barcodes were modeled assuming a constant %CV value for a given dye and imposing a minimal distance in between intensity barcodes corresponding to 3.5 times the measured standard deviation (See Figure S8), hence expecting to include over 99 % of a given BMPs from a gaussian distribution. We limited the total amount of oligos to 90 pmol for all barcodes.

### Automatic decoding of BMPs

Mixture of BMPs were automatically decoded based on their 2D intensity (Bead_590_, AF647/Bead_590_) via 2D clustering, analogous to Dagher et al.^[29]^ Briefly, intensity data sets ***I*** were model with a gaussian mixture model (GMM) using the expectation-maximization (EM) algorithm^[62]^ to the probability distribution function given as:

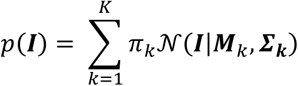

where **M**_k_ and 𝚺_k_ are means and covariances of the *k* gaussian given by 𝒩(𝑰|𝑴_k_, 𝜮_𝒌_), respectively, and 𝜋_k_ is the mixing coefficient, corresponding to a normalized metric denoting how well BMPs fit the k^th^ gaussian. *K* corresponds to the maximum number of unique barcodes to be decoded. Modeled intensity of each barcode was used as the initial value of the mean in the GMM, 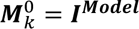. The initial covariance matrix value was a diagonal matrix with 10 %CV in each dimension 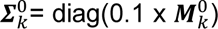, in accordance with our measured %CV values. Initial clusters probabilities were considered homogeneous (𝜋^O^ = 1/𝐾). When experimental data sets ***I*** are introduced, 𝑝(𝑰) measured the likelihood of that current data set to be fitted by the GMM clusters. Then, the probability 𝜙 of a given BMP 𝜓 belonging to the cluster *k*, also referred to as the posterior probability, is calculated during the expectation step of the EM search using:

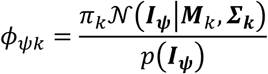

We then performed a maximization step, where values of the Gaussian components (𝑴_k_, 𝜮_𝒌_ and 𝝅_𝒌_) are updated to maximize the log-likelihood ln(𝑝(𝑰)). The process was repeated for up to 5,000 iterations or until condition for convergence, defined as ln(𝑝(𝑰_𝒊_)) – ln(𝑝(𝑰_𝒊-𝟏_)) < 1e^-7^, was reached. Finally, a posterior probability threshold, ranging from 60 to 80%, was applied, thereby eliminating BMPs with low 𝜙.

### Assessment of antibodies cross-reactivity in bulk

The cAb_oligo_ were anchored on their respective BMP as described above and combined at a concentration of 100 BMPs per barcode per microliter. Prior to the assay, cAb-BMPs were first blocked with PBST0.05 + 1% BSA + 1mM biotin for 1 h. Then, BMPs were rinsed thrice with PBST0.1 via cycle of magnetic aggregation and resuspension in 100 μL of PBST0.1 per cycle using a liquid dispenser (MultifloFX, BioTek). For all experiments, incubations were performed in V-bottom 96-well microplate (Greiner bio-one, Germany) at room temperature with horizontal shaking at 950 rpm. Incubations with cytokines were performed for 120 min at a specific concentration (1 ng mL^-1^ or 100 ng mL^-1^), rinsed thrice with 100 μL PBST0.1 and incubated with detection antibodies (dAbs) for 1 h at 5 μg mL^-1^ and 2 μg mL^-1^ in PBST0.05 + 1% BSA for individual and cocktail incubation, respectively. BMPs were rinsed thrice with PBST0.1 and immobilized onto poly-L-lysine coated imaging 96-well microplate (Greiner-Bio-one) by incubating the BMPs for 10 min. Wells were gently rinsed with PBST0.05 + 1% BSA and incubated for 60 min with 2 μg mL^-1^ BV421-labeled streptavidin (SA) and rinsed thrice with PBST0.1 prior to imaging. BMPs were imaged by CLSM while being submerged in PBST0.1. Signal-to-noise ratio (SNR_dAbs_) was calculated by subtracting cAb-BMP specific mean background in the absence of cytokine (*n = 3*) from the calculated MFI_BV421/590_ signal and normalized by the global standard deviation across all BMPs (*n* = 1*8*) of the SA background. Similarly, SNR_Ag_ was calculated by subtracting cAb-BMP specific mean assay background in presence of dAbs cocktail (*n = 5*) from the calculated MFI_BV421/590_ signal and normalized by the global standard deviation of the assay background across all BMPs (*n = 35*).

### Microfluidic Device Fabrication

Microfluidic designs were drawn using Fusion 360 (Autodesk, 2022) and printed on a high-definition transparency photomask (50,800 DPI, Fineline Imaging, Colorado Springs, CO). Microfluidics devices were fabricated with polydimethylsiloxane (PDMS) by standard soft lithography procedures.^[63]^ Briefly, 35 μm layer of SU-8 (GM 1060 negative resin, Gersteltec, Engineering Solution, Switzerland) was spin-coated onto a 6-inch silicon (Silicon Quest International, Santa Clara, CA) wafer. Following soft baking, the photomask was aligned onto the wafer and was exposed to UV light at 365 nm (Hg i-line). Subsequently, the SU-8 was post-baked and developed in propylene glycol monomethyl ether acetate (PGMEA; Sigma-Aldrich) for 5 min. Following hard baking, the silicon mold was ready for the subsequent steps. Note that all pre-bake, post-exposure bake, and hard bake steps were performed using the standard temperature and time ramps on a programmable hot plate (HS40A, Torrey Pines Scientific, CA). The resulting mold was used for downstream fabrication of PDMS replica without further functionalization. To generate devices, mold was inserted into a laser cut circular hollow plate made of PMMA and immobilized using extra tack Frisket film (Grafix, Amazon), similarly reported by Li et al. (Chips and Tips; A Simple and Economical Holder for Casting PDMS Chips). Then, a 10:1 ratio of PDMS and curing agent were vigorously mixed, degassed for one hour under vacuum, poured onto the mold and cured for at least three hours at 65 °C. PDMS was then cut and peeled off from the mold. Inlets and outlets holes were punched using a 1.2 mm biopsy punch (Harris Uni-Core^TM^, Ted-Pella, Inc). Glass slides (75 x 50 mm) were cleaned with isopropanol with sonication for 5 minutes and thoroughly dried using nitrogen gas. Both glass slides and PDMS devices were activated for 35 seconds with an air plasma (Plasma Etch, Inc., NV) with 300 mTorr pressure and 150 mW power and bonded together. Devices were incubated at 65 °C overnight to strengthen the glass and PDMS bonding efficiency. Prior their use, devices were treated with a 0.22 μm filtered solution of 2% v/v trichloro(1H,1H,2H,2H-perfluorooctyl)silane (Sigma-Aldrich) in Novec 7500 fluorinated oil (3M, Canada), rinsed with 0.22 filtered Novec 7500 and dried by pressured nitrogen blow.

### Alginate microgel- and PiPs generation

Alginate microgels were generated by adapting the competitive ligand exchange crosslinking (CLEX) reported by Håti et al.^[35]^ by droplet-based microfluidics. Briefly, two aqueous solutions were prepared: (**1**) a zinc-rich solution composed of 42 mM Zn-EDDA, 40 mM MOPS pH 7.0, 1% w/v alginate and 40 % v/v completed RPMI-1640 or MilliQ water, and (**2**) a calcium-rich solution composed of 42 mM Ca-EDTA, 40 mM MOPS pH 7.0 and 1% w/v alginate and 40% v/v completed RPMI-1640 or MilliQ water (see *Preparation of the precursor solutions* above). The oil phase was composed of 2% w/w fluorosurfactant (Ran Biotechnologies, MA) in Novec 7500. All solutions were filtered through a 0.22 μm pore filter and loaded in 1 mL luer-lock plastic syringe (BD) equipped with a ½ 23G blunt needle (McMaster Carr) with FEP tubing (1/16 OD, 1/32 ID; Cole Palmer) and introduced into the microfluidic device via neMESYS syringe pumps (Cetoni GmbH, Germany). Flow rates of the two aqueous solutions were set to 100 μl hr ^-1^ and the oil phase was set to 450 μL hr ^-1^. The produced W/O droplets were collected into a 1.5 mL Eppendorf on ice. To release the alginate hydrogels from the emulsion, the excess oil was removed, and 200 μL of releasing buffer, correspond to 25 mM MOPs pH 7.4 supplemented with 130 mM NaCl, 2 mM CaCl_2_ and 2% w/v BSA, was added on top of the emulsion. Then, the droplets were coalesced by introducing 150 μL of 20% v/v H,1H,2H,2H-Perfluoro-octanol (PFO; Sigma-Aldrich) in Novec 7500 and vigorously mixed by pipetting to fully saturate the droplet surface, while minimizing perturbation of the top medium layer. Droplets were incubated for 10 min for complete coalescence and release of the hydrogels, which relocated to the top aqueous phase. Released hydrogels were carefully resuspended by pipetting, collected and transferred into a new Eppendorf tube. Care was taken to prevent collection of oil. For the experiments with cells encapsulated inside the alginate hydrogels, the releasing buffer was exchanged to 400 μL of completed RPMI-1640 supplemented with 2 mM CaCl_2_. During droplet coalescence, the emulsion was incubated at 37 °C for 30 min. To generate PiPs, cells and/or BMPs were supplemented to one of the soluble alginate phases (e.g., BMPs in Zn-EDDA and Cells in Ca-EDTA). Cells and BMPs were not supplemented to the same alginate phase. When PiPs were formulated without cells, the completed RPMI-1640 was substituted by MilliQ water.

### Multiplex assays of cytokine in bulk and PiPs

The cAb-BMPs anchoring the cAb_oligo_ to their corresponding BMPs were prepared as described above, and combined at a concentration of 100 BMPs per barcode per microliter. Pooled cAb-BMPs were then blocked with PBST0.05 + 1% BSA + 1 mM biotin. For assay in PiPs, we first rinsed the cAb-BMPs thrice with 25 mM HEPES pH 7.4 supplemented with 0.1% v/v Tween-20 and 135 mM NaCl that emulate our PBST0.1. This washing step was used to remove phosphate-based buffer that may compete for Ca^2+^ ions. Rinsed cAb-BMPs were formulated into PiPs as described earlier in MilliQ at a final concentration of 100 BMPs per barcode per µL. Following PiPs generation, PiPs were isolated in 600 μL tube and rinsed thrice with an assay buffer, consisting of 25 mM MOPS pH 7.4 supplemented with 130 mM NaCl, 2 mM CaCl_2_, 2% w/v BSA and 0.05% v/v Tween-20, through cycles of centrifugation at 400 G for 2 min, resuspension in 200 μL of assay buffer and incubated for 10 min on a rotary shaker at 350 rpm to enable diffusion. All subsequent rinsing steps for PiPs involved the same procedure. After the last rinsing cycle, PiPs were centrifuged and resuspended in 200 μL of assay buffer containing various concentrations of cytokine standards. Incubations were performed in 600 μL tubes. For incubations in bulk, the blocked and pooled cAb-BMPs were pipetted into V-shaped 96 well plates and rinsed with assay buffer thrice via cycles of magnetic aggregation and resuspension in 200 μL of assay buffer. Then, cAb-BMPs were incubated with various concentrations of cytokine standards. All rinsing steps for bulk cAb-BMPs were performed using assay buffer, as described above. For all incubations, wells were sealed with adhesive PCR plate seals to minimize solvent evaporation (Thermo Fisher). To establish binding curves and LODs, cytokine mixtures were incubated for 120 min at the specified concentrations and rinsed. BMPs were incubated for 120 min with dAb cocktails at a concentration of 2 μg mL^-1^ per dAb and rinsed. Prior to labeling, assays using bulk cAb-BMPs were pipetted in a poly-L-lysine coated 96-well imaging microplates and incubated for 10 min to allow electrostatic immobilization of the BMPs. After carefully removing the supernatant, cAb-BMPs were incubated with 2 μg mL^-1^ BV421-labeled SA for 60 min and rinsed thrice with PBST0.1 by removing the supernatant and adding new PBST0.1. The well plate was gently agitated for 10 min per rinse using a rotary orbital shaker at 350 rpm. For PiPs assays, PiPs were centrifuged, resuspended in 200 μL of assay buffer containing 2 μg mL^-1^ BV421-labeled SA and incubated for 60 min in their corresponding 600 μL test tube. Following rinsing, PiPs were transferred into the same imaging well plates and both bulk cAb-BMPs and PiPs were imaged by CLSM using the same procedure as described above. SNRs were calculated by subtracting the specific cAb-BMP assay background MFI_BV421/590_ (*n = 5*) and normalizing by the global standard deviation across all barcodes (*n = 35*) of the assay background.

### Cell viability assessment

In a 6 wells plate, cells (1 x 10^6^ cells mL^-1^) were differentiated with 25 nM PMA as described above. Following differentiation, Mφ THP-1 cells were rinsed with PBS pH 7.4 and stained with Calcein-AM/Ethidium homodimer-1 (LIVE/DEAD Cell viability assay, ThermoFisher) based on manufacturer’s procedure. When cells were entrapped in hydrogels, 0.5 μL calcein-AM and 2 μL ethidium homodimer-1 stock solution (stock from manufacturers) were diluted into 1 mL of buffer composed of 25 mM MOPS, 130 mM NaCl and 2 mM CaCl_2_. The hydrogels were washed with 25 mM MOPS pH 7.4, 130 mM NaCl and 2 mM CaCl_2_, centrifuged at 200 G for 1 min and resuspend in 100 μL of the diluted solution of LIVE/DEAD stain in MOPS buffer. Cells were incubated at room temperature for 30 min, protected from light, prior to confocal imaging. It is noteworthy that DEAD stains, which typically stain duplexed DNA, will stain the BMPs in a homogeneous way, leading to a strong fluorescence background in the bead (561 nm excitation) channel. Fluorescence images were analyzed with Fiji.^[64]^ Each fluorescence channel was pseudo flat-field corrected employing BioVoxxel toolbox (https://doi.org/10.5281/zenodo.10050002), contrast enhanced (0.35), Otsu’s thresholded, features were then dilated, watershed segmented and analyzed via the Particle Analyzer function by adjusting the minimal- and maximal area value to the targeted feature size (cells, hydrogels, or satellite droplets). A circularity of 0.3 and 0.6 were fixed for cells and hydrogels/droplets, respectively.

### Droplet leakage assessment

To assess potential leakage or cross-contamination between droplets during incubation, 1 nM of 150 kDa FITC-dextran or 1 μg mL^-1^ of AF647-labeled BSA were individually encapsulated into 50 μm droplets with phenol-red free RPMI-1640 supplemented with 25 mM HEPES pH 7.4, 2 mM CaCl_2,_, 10% HI-FBS, 1% penicillin/streptomycin by droplet microfluidic. Droplets were produced using flow rates of 500 μL h^-1^ for the aqueous phase and 1000 μL h^-1^ for the oil phase. The oil phase was composed of 2% w/w fluorosurfactant in Novec 7500. After their generation, droplets were combined in a 1:1 volume ratio in a 1.5 mL Eppendorf tube. A droplet aliquot was directly imaged by CLSM to assess potential leakage at time *t* = 0, while the rest were incubated for 18 h at 37 °C, 5% CO_2_ prior to their assessment by CLSM.

### Cell encapsulation in PiPs and sandwich immunoassay

Prior to performing single cell immunoassays on THP-1 cells, Fc receptors on THP-1 cells were blocked for 10 min with 5 μL of Human TruStain FcX (Biolegend) based on manufacturer’s recommendation to reduce nonspecific adsorption of dAbs prior to their encapsulation in PiPs. Following blocking, Mφ THP-1 cells were rinsed once with PBS and detached from the culture flask with TrypLE^TM^ (Gibco) for 10 min at 37 °C. The suspended cells were diluted with completed RPMI-1640 and centrifuged at 300 G for 5 min. Cells were concentrated into 5 mL completed medium and counted. The desired number of cells were then centrifuged at 300 G for 5 min and resuspended 0.22 μm filtered Ca-EDTA rich alginate phase (42 mM Ca-EDTA, 40 mM MOPS pH 7.0, 1% w/v alginate, 40% v/v completed RPMI-1640) at a final concentration of 18 x 10^6^ cells mL^-1^. Independently, blocked cAb-BMPs were combined to a final concentration of 30 000 BMPs μL^-1^ per barcode and resuspended in Zn-EDDA rich phase (42 mM Zn-EDDA, 40 mM MOPS pH 7.0, 1% w/v alginate, 40% v/v RPMI-1640 and 0.01% v/v Tween-20). Each alginate phase was loaded into 200 μL sterile pipette tip pre-loaded with Novec 7500 as described elsewhere (see Figure S31).^[65]^ Water-in-oil droplets were generated with a flow focusing junction as described above. Droplets were collected on ice, excess of oil removed and coalesced with 20% v/v PFO in Novec 7500. Then, 400 μL of phenol-red free RPMI-1640 supplemented with 25 mM HEPES pH 7.4, 2 mM CaCl_2_, 10% HI-FBS and 1% penicillin/streptomycin, referred to as PiPs medium, was layered on top of the emulsion. Droplets were incubated at 37 °C during the droplet releasing process for a period of 30 min.

PiPs were centrifuged at 200 G for 1 min, resuspended in 400 μL of PiPs medium, and incubated for 5 min to enable diffusion in/out the hydrogels. These steps were repeated 3 times to ensure adequate washing. To stimulate cytokine secretion by Mφ THP-1 cells, the 2 last washing steps were performed by supplementing 2X LPS (200 ng mL^-1^; Invitrogen) solution in PiPs medium and incubated for 2 min each round. Then, the hydrogels were centrifuged at 200 G for 1 min and the supernatant removed. PiPs were re-emulsified by adding 150 μL of 2% w/w fluorosurfactant in Novec 7500, and vigorously mixed by pipetting up and down, leading to the confinement of individual PiP. The resulting *De novo* droplets, now entrapping cells and cAb-BMPs inside PiPs, were incubated for 6 h at 37 °C in a closed Eppendorf tube. Following incubation, PiPs were released by droplet coalescence with 20% v/v PFO in Novec 7500 and placed on ice. Cell laden PiPs were diluted with an assay buffer deprived from Tween-20 (25 mM MOPS pH 7.4, 130 mM NaCl, 2 mM CaCl_2_ and 2 % w/v BSA) to maximize cell viability, centrifuged at 4°C at 200 G for 1 min, washed with 200 μL of ice-cold assay buffer and incubated 5 min on ice. These washing steps were repeated thrice.

Isolated PiPs were centrifuged at 200 G for 1 min, rinsed with 200 μL of ice-cold assay buffer and incubated for 5 min on ice thrice. PiPs were diluted 1:1 on ice with dAbs cocktail for 120 min at a concentration of 4 μg mL^-1^ per dAb in assay buffer, rinsed thrice while incubating 10 min per wash with ice-cold assay buffer, diluted 1:1 with 4 μg mL^-1^ BV421-labeled SA for 60 min on ice, and finally rinsed thrice with ice-cold assay buffer with a 10 min incubation per wash cycle. PiPs were resuspended in assay buffer and loaded into an imaging 96-well plate and imaged by CLSM as described next.

### Fluorescence confocal imaging of cell-loaded PiPs

To image cells, BMPs, and PiPs by CLSM, PiPs suspension was diluted 1:5 in assay buffer supplemented with 1 nM 2MDa FITC-dextran (Sigma-Aldrich) and 4 μM calcein-AM. The solution was then transferred into a 96-wells imaging microplate which has been coated with 0.1% w/v poly-L-lysine for 30 min, rinsed with MilliQ water and dried prior to the addition of the PiPs. PiPs were incubated 5-10 min to allow sedimentation and immobilization, while enabling the increase of calcein signal inside viable cell’s cytosol. Cell-loaded PiPs were imaged as described above at room temperature.

### Fabrication of microfluidic traps by 3D printing-based replica molding

Microfluidic traps were used for permeability testing, but not for the assays. Microfluidic molds were designed in Fusion 360, exported as .STL file for slicing in the third-party software CHITUBOX (CBD technology Ltd., v1.9.5) at a layer thickness of 20 μm. Slices were then uploaded to an Elegoo Mars 3 Pro masked stereolithography LCD 3D printer (ELEGOO, Shenzhen, China) equipped with a 405 nm light source. The molds were printed in Miicraft BV-002A resin (Creative CADworks, Canada, ON) with the following settings: *Bottom Layer Count: 5, Anti-Aliasing: On, Grey-Level: 2, Base Exposure Time (BET): 25, Layer Height: 20 μm, Layer Exposure Time (LET): 2.3 seconds.* Following 3D printing, molds were washed on the build plate with isopropanol to remove excess of uncured resin and dried with nitrogen. Molds were then submerged in an isopropanol bath for 30 min with gentle agitation on a rotary shaker, dried with nitrogen followed by 4 min of exposure in a UV chamber (CureZone MKII, Creative CADworks), baked at 65 °C overnight and exposed to a second 4 min UV exposure. Degassed PDMS (10:1 ratio) was poured into the 3D-printed molds and cured overnight at 65 °C. PDMS was then cut and peeled off from the mold. Inlets and outlets holes were punched using a 1.2 mm biopsy punch. Glass slides (25 x 60 mm) were cleaned with isopropanol with sonication for 5 min and thoroughly dried using nitrogen gas. Both glass slides and PDMS devices were activated for 35 s with an air plasma with 300 mTorr pressure and 150 mW power and bonded together. Devices were incubated at 65 °C overnight to strengthen the glass and PDMS bonding efficiency. Prior to their usage, devices were plasma activated for 2 min and functionalized 0.1% w/v poly-L-lysine solution for 30 min. Devices were then rinsed with MilliQ water and dried by pressured nitrogen blow.

### Permeability assessment of PiPs

PiPs encapsulating BMPs (∼20 000 BMPs μL^-1^ per barcode) were formulated by droplet-based microfluidic as described above in the absence of RPMI-1640 medium. PiPs were isolated from the oil phase and washed thrice with diffusion buffer, containing 25 mM MOPS pH 7.4 supplemented with 130 mM NaCl and 2 mM CaCl_2_, using cycles of centrifugation and resuspension. PiPs were then resuspended in diffusion buffer supplemented with 1 nM 2MDa FITC-dextran and perfused into the microfluidic trap at 10 μL min^-1^. Once PiPs reached the trapping area (lower height of the microchannel), flow was stopped for 5 min to enable adhesion of PiPs to the microfluidic walls. CLSM images were acquired to assess baseline fluorescence signal of the BMPs using the same optical configuration as described above. Then, diffusion buffer supplemented with 2 μg mL^-1^ BV421-labeled SA, 4 μg mL^-1^ AF488-labeled Goat anti-rabbit antibody (Invitrogen) and 1 nM 2MDa FITC-dextran was perfused at a rate of 10 μL min^-1^ while imagining PiPs at an interval of 5 min for a period of 1 h. BMPs signals were extracted for each time point and normalized by the maximum fluorescence value across all BMPs for each fluorescence channel. Data were then model with a 1D accumulation model, defined as *f(t)* = *A(1-exp(-(t_0_-λ)/τ)* where *A*, *λ* and *τ* are fit parameters and *t_0_* is the time offset to account for the delayed perfusion of SA and Ab. The extracted value of *τ* corresponds to the characteristic diffusion constant.

### Cell retrieval from alginate-based PiPs

Cells (18x10^6^ cells mL^-1^) and BMPs (210 000 total BMPs μL^-1^) were encapsulated in alginate to produce the PiPs as described above. Following droplet coalescence and isolation of PiPs, cell laden PiPs were washed with assay buffer deprived from Tween-20, concentrated by centrifugation and resuspended in 300 μL of assay buffer and kept on ice for 3 h to emulate immunoassay incubations. Then, to isolate the cells from the PiPs, we adapted a protocol previously reported by Rowley *et al.^[47]^* Briefly, 100 μL of PiPs suspension was added to 400 μL of trypsin-EDTA (Gibco) for 5 min at 37 °C. Then, 1 mL of 2% w/v citrate (Sigma-Aldrish) at pH 7.4 was added. The solution was then incubated for 20 min at 37 °C. Following incubation, BMPs were removed by magnetic aggregation for 2 min, and the supernatant was transferred to a new Eppendorf tube. Cells were then isolated by centrifugation for 5 min at 180 G. Cell viability assessment was performed as described above on PiPs (prior to cell isolation) and following cell retrieval.

### Antibody microarray profiling of cytokines in cell supernatant

Antibody microarrays were employed to detect secreted cytokine by differentiated Mφ THP-1 cells in cell supernatants upon LPS stimulation. Briefly, antibody microarrays were patterned onto Aldehyde functionalized glass slides (PolyAn, Germany) using a sciFLEXARRAYER SX inkjet bioprinter (SCIENION GmbH, Germany) equipped with a single piezo dispense capillary nozzle with coating 1 (SCIENION). Prior to antibody immobilization, glass slides were inspected by fluorescence imaging with a InnoScan 1100 AL microarray scanner (Innopsys, France) at 532 nm and 100% gain to assess the quality of the aldehyde functionalized glass slide. Pristine capture antibodies (Table S3) were diluted to 100 μg mL^-1^ in 15% v/v 2,3-butanediol (Sigma-Aldrich) and 1 M betaine (Sigma-Aldrich) in pH 7.4 PBS. A Goat-anti rabbit antibody (Thermo Fisher Scientific, USA) was diluted and added to the antibody panel to serve as negative control. Antibody solutions were spotted in 16 (2 x 8) subarrays of 100 (10 x 10) 100 μm spots size using 500 μm pitch value. Patterned glass slides were incubated overnight at room temperature and 70% humidity, washed thrice with PBST0.01 (while incubating 5 min in between washes on a rotary shaker). Then, the glass slides were blocked with PBST0.1 supplemented with 3% w/v BSA (PBST0.1 + 3% BSA) at room temperature, washed thrice with PBST0.1, dried and inserted within a 16-well ProPlate multi-well chambers (Grace Bio-Labs, OR) to enclose the fabricated antibody microarrays. Cell supernatants (PMA^+^/LPS^-^ and PMA^+^/LPS^+^) were sampled out after 3 h of LPS stimulation and centrifuged at 1000 G for 5 minutes. The supernatant was then filtered using a 0.22 μm filter and incubated for 2 h on the antibody microarray at room temperature at 450 rpm on a rotary shaker. Blank and cytokine standards (1pg mL^-1^ to 100 ng mL^-1^) were prepared in phenol-red free RPMI-1640 supplemented with 10 % HI-FBS, 1% penicillin/streptomycin, 25 mM HEPES pH 7.4 and 2 mM CaCl_2_ to emulate cell conditions once entrapped in PiPs in *de novo* droplets. Wells were washed thrice with PBST0.1 + 3 % BSA with 5 min incubations. A detection antibody mixture (2 μg mL^-1^ per antibody; Table S3) in assay buffer deprived from Tween-20 (25 mM MOPS pH 7.4, 130 mM NaCl, 2 mM CaCl_2_, and 2% w/v BSA) was incubated for 2 h onto the wells at room temperature at 450 rpm. Wells were rinsed thrice with assay buffer, labeled by incubating with 1 μg mL^-1^ AF647 labeled streptavidin in assay buffer for 1 h, protected from light at room temperature at 450 rpm. The wells were washed thrice PBST0.1, rinsed once with MilliQ water, dried, and directly imaged on with an InnoScan 1100 AL microarray scanner. Images were exported as .TIFF files and quantified using ArrayPro software (Meyer Instruments, v4.5).

### Assessment of polyfunctionality and multivariate analysis

Single cell data was extracted, analyzed, and visualized using custom written codes in MATLAB. Multicolor fluorescence z-stacks for the different cell samples were acquired as described above. From the Calcein/FITC channel, we localized both PiPs and cells. PiPs were located by calculating the minimal *Z* intensity projection. The projection was binarized via thresholding the image, segmented using a watershed algorithm leading to the localization of individual PiPs. Potential alginate aggregates or odd-shaped objects, which exhibited a circularity value below 0.60, were eliminated from further analysis. Cells were located by calculating the maximum *Z* intensity projection, followed by thresholding and watershed segmentation. Cell-containing PiPs were identified by comparing each cell’s centroid to each detected PiP boundaries. PiPs containing more than two cells were excluded from further analysis. Then, BMPs were segmented based on the 561/590 nm channel to extract their *XY* coordinates. Each BMP was then associated to its corresponding PiP by comparing its *XY* coordinates to the extracted PiP boundary. The BMPs with XY coordinates located within the boundaries of a given PiP were then assigned to that PiP for further analysis, while those falling outside all PiP regions were excluded. Then, a spherical mask having a 3 μm dimater was generated and positioned on each segmented BMP centroid and subsequently used to integrate fluorescence intensity all fluorescence channels (BV421, calcein/FITC, 561/590 and AF647). The fluorescence intensities of the BV421 channel dedicated for immunoassay was corrected by subtracting the local background of the beads extracted from a 3D donut-shaped mask, analogeous to the spherical mask mentioned above. To prevent artefact due to bleed-through from the Calcein signal into the 561/590 nm fluorescence channel, BMPs located in *X, Y, Z* within area *X, Y* occupied by a cell (i.e. BMPs located above/below a cell in 3D) were excluded from the analysis.

Fluorescence intensity of the BV421 and AF647 channels for each BMPs were normalized by their respective 561/590 scattering signal. The normalized BMPs were gated by applying a lower and upper intensities thresholds based on the 561/590 signal corresponding, respectively, to 0.65 and 1.45 times the median value across all BMPs. Remining BMPs were decoded using a GMM as described above. Assay signal (BV421 channel) of BMPs encoding the same cytokine that were co-entrapped within the same PiPs were summed together to account for cytokine partitioning among these BMPs. Data of BMPs signals entrapped in PiPs without cell were used to assess assay background level of each cytokine. These data were used to generate cytokine abundance histograms which were then fitted by a normal distribution. The mean and standard deviation of the distribution were used to calculate assay LOD for each cytokine, defined as 3-fold the standard deviation above the mean background. These LODs were used to segregate cytokine-secreting cells and non secretor cells. Further downstream analyses were restricted to PiPs entrapping, at least, 5 distinct cytokine-encoded BMPs for each condition. This insured that a minimum of 5 cytokines per cells would be measured in multiplexed per conditions. Missing cytokine assay signals were imputed by the average cytokine value measured for the corresponding target across all BMPs in the same sample. These data were then used for further multivariate analysis and assessment of the PSI of the cells.

The PSI is defined as the total functional intensity contributed by all polyfunctional cells, that are cells co-secreting at least 2 cytokines above their respective assay LOD. The PSI of each condition was computed as the percentage (%) of polyfunctional cells relative to all measured cells, multiplied by the sum of the mean fluorescence intensity of each assayed cytokines secreted by those cells^[1]^:

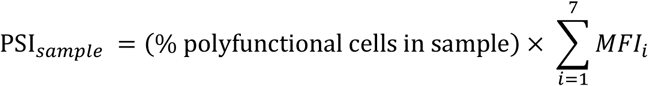

We presented PSI for each sample conditions as segmented bars highlighting contribution of each group of cytokines, namely effector, stimulatory, regulatory and inflammatory. The t-SNE analysis was conducted on 527 cells corresponding to PiPs encapsulating a single cell and at least 5 distinct cytokine-encoded BMPs. Due to the low number of variable (7 cytokines), dimension reduction by principal component analysis was not performed prior analysis. The t-SNE algorithm was run using default parameter value within the *tsne function* in MATLAB.

### Statistical Analysis

For single cell assay using PiPs, statistical analysis was performed using Mann-Whitney two-tailed test while unpaired, two-tailed *t*-test was used for bulk assay by using MATLAB. Data were presented as mean ± S.D. The sample size of figures requiring statistical analysis is stated in the corresponding figure caption. For all statistical tests, confidence levels of *\*P* < 0.05, *\*\*P <* 0.01, and *\*\*\*\*P* < 0.0001 were used as the threshold values.

## Acknowledgements

The funding was provided by the Cell and Gene Therapy Program, National Research Council Canada. We would like to thank Prof. Nicole Li-Jessen from McGill University for providing the THP-1 cells. We would also like to thank Hicham Saad and Michael Martin from Nikon (Ontario, Canada) for fruitful discussions and support for microscopy imaging. We thank Andreas Wallucks for help and discussion for the multivariate analysis.

## Authors contribution

F.L., D.J., L.M., T.V., conceptualized the experiments. F.L. performed, acquired, and curated all data. F.L. Designed the microfluidic devices. B.M., and M.J.P. prepared the photomask. L.L. prepared the microfluidic master wafer. F.L. developed the image processing pipeline. F.L., Y.M., designed and 3D printed microfluidic traps. M.S., prepared the Antibody microarray. F.S., optimized the hydrogel purification., D.J. L.M, A.N and T.V., acquired the fundings. D.J., L.M., and T.V. supervised the project. F.L. wrote the manuscript. All authors commented on the draft manuscript.

## Data Availability Statement

The data that support the findings of this study are available from the corresponding authors upon reasonable request.

