## Supporting Information for "PiP-plex: A Particle-in-Particle System for Multiplexed Quantification of Proteins Secreted by Single Cells"

**Table S1**: List and sequence of oligonucleotides used in this study

| Strand IDs | Modification | Sequence | (nt) |
|---|---|---|---|
| Capture Oligo (CO) | Biotin | 5'-Biotin - TTT TTT TTT GTG GCG GCG GTG ATT GGT TAT TGA GAG TTT ATG– 3' | **42** |
| Spacer Oligo (SO) | Biotin | 5'-Biotin - TTT TTT TTT GTG GCG GCG GTG – 3' | **21** |
| Encoder Oligo$_0$ ($E_0$) | | 5' – CAC CGC CGC CAC AAA AAA AAA – 3' | **21** |
| Encoder Oligo$_{FAM}$ ($E_{FAM}$) | FAM | 5' – CAC CGC CGC CAC AAA AAA AAA – FAM – 3' | **21** |
| Encoder Oligo$_{Cy3}$ ($E_{Cy3}$) | Cy3 | 5' – CAC CGC CGC CAC AAA AAA AAA – Cy3 – 3' | **21** |
| Encoder Oligo$_{Cy5}$ ($E_{Cy5}$) | Cy5 | 5' – CAC CGC CGC CAC AAA AAA AAA – Cy5 – 3' | **21** |
| Encoder Oligo$_{ATTO488}$ ($E_{ATTO488}$) | ATTO488 | 5' – CAC CGC CGC CAC AAA AAA AAA – ATTO488 – 3' | **21** |
| Encoder Oligo$_{ATTO550}$ ($E_{ATTO550}$) | ATTO550 | 5' – CAC CGC CGC CAC AAA AAA AAA – ATTO550 – 3' | **21** |
| Encoder Oligo$_{AF647}$ ($E_{AF647}$) | AF647 | 5' – CAC CGC CGC CAC AAA AAA AAA – AF647 – 3' | **21** |
| Anchor Oligo (AO) | 5' Thiol Modifier C6 S-S | 5'-Thiol - CTC ATT CGC CAT AAA CTC TCA ATA ACC AAT – 3' | **30** |

* (nt): Nucleotides

**All DNA sequences were acquired directly modified from Integrated DNA Technologies (IDT) with HPLC purification.

**Table S2**: List of bioreagents used in this study

| Target | Supplier | Cat.no. | Lot |
|---|---|---|---|
| **Antigen** | | | |
| IL-2 | RnD | 202-IL-010 | AE6421081 |
| TNF-α | RnD | 210-TA-020 | DDHB0422021 |
| IFN-γ | RnD | 285-IF-100 | RAX2422031 |
| IL-4 | Biolegend | 204-IL-010 | NCL0722051 |
| IL-17a | Biolegend | 570509 | NA |
| MIP-1α | RnD | 270-LD/CF | LG1322031 |
| IL-5 | Biolegend | 560701 | B297119 |
| **Capture Antibody** | | | |
| Rat anti human IL-2 | Biolegend | 500302 | B359106 |
| Mouse anti human TNF-α | Biolegend | 502802 | B362273 |
| Mouse anti human IFN-γ | Biolegend | 502402 | B322937 |
| Mouse anti human IL-4 | Biolegend | 500702 | B362398 |
| Mouse anti human IL-17a | Biolegend | 512702 | B342003 |
| Goat anti human MIP-1α | RnD | AF-270-NA | MP1122031 |
| Rat anti human IL-5 | Biolegend | 501002 | B330830 |
| Goat anti Rabbit IgG | Invitrogen | A16112 | 70-54-072919 |
| **detection Antibody/protein** | | | |
| Biotin Goat anti human IL-2 | Biolegend | 517605 | B369547 |
| Biotin Mouse anti human TNF-α | Biolegend | 502904 | B353934 |
| Biotin Mouse anti human IFN-γ | Biolegend | 502504 | B360065 |
| Biotin Rat anti human IL-4 | Biolegend | 500803 | B247643 |
| Biotin Goat anti human IL-17a | Biolegend | 518902 | B346127 |
| Biotin Goat anti human MIP-1α | RnD | BAF270 | UU1122021 |
| Biotin IL-5 | Biolegend | 501002 | B330830 |
| BV421 Goat anti mouse IgG | Biolegend | 405317 | NA |
| BV421 streptavidin | Biolegend | 405226 | B397508 |
| AF647 Streptavidin | Invitrogen | S21374 | 2145944 |

| | | | |
|---|---|---|---|
| TruStain FcX | Biolegend | 422301 | B385746 |
| Biotin Goat anti Rabbit IgG | Invitrogen | 31822 | WJ3418985 |
| AF488-labeled Goat anti-rabbit antibody | Invitrogen | A-11008 | NA |
| Mouse IgG | Biolegend | 400264 | B419942 |
| **Beads** | | | |
| Dynabeads M-270 Streptavidin | Invitrogen | 65305 | 01252313 |
| 3.0 µm Streptavidin Coated Microspheres | Bangs Laboratories, Inc | CP01005 | L210331C |

**Table S3** List of materials and reagents used in this study:

| **Name** | **Supplier** | **Cat.no.** |
|---|---|---|
| Alginate (UP-MVG) | Novamatrix | 4200101 |
| Sulfosuccinimidy 4-(N-maleimidomethyl) cyclohexane-1-carboxylate (Sulfo-SMCC) | Thermo Fisher Scientific | 22322 |
| DynaBeads M-270 Streptavidin | Thermo Fisher Scientific | 65305 |
| Ethylenediamine-N,N'-diacetic acid (EDDA) | TCI America | E0275 |
| Ethylenediaminetetraacetic acid (EDTA) | Sigma-Aldrich | E9884 |
| zinc acetate ($Zn(CH_3CO_2)_2$) | Sigma-Aldrich | 383317 |
| anhydrous calcium chloride ($CaCl_2$) | Sigma-Aldrich | C5670 |
| Trisodium citrate dihydrate, 99% | Thermo Fisher Scientific | A12274.0E |
| Biotin | Sigma-Aldrich | B4639 |
| (N-(2-hydroxyethylpiperazine-N-2-ethane sulfonic acid (HEPES) | Sigma-Aldrich | H4034 |
| 3-(N-Morpholino)propanesulfonic acid (MOPS) | Sigma-Aldrich | M3183 |
| trichloro(1H,1H,2H,2H-perfluorooctyl)silane | Sigma-Aldrich | 448931 |
| poly-L-lysine, 0.1% solution in water | Sigma-Aldrich | SLCP1100 |
| RPMI-1640, with glutamine | Gibco | 11875093 |
| Phenol-red free RPMI-1640, with glutamine | Gibco | 11835030 |
| TrypLE™ | Gibco | 12605-010 |

| | | |
|---|---|---|
| 0.25% Trypsin-EDTA (1X) | Gibco | 25200-056 |
| Heat inactivated fetal bovine serum (HI-FBS) | Gibco | 12484-028 |
| penicillin/streptomycin | Gibco | 15140122 |
| phorbol-12-myristate-13-acetate (PMA) | Sigma Aldrich | P8139 |
| Lipopolysaccharide (LPS) Solution (500X) | Invitrogen | 00-4976-93 |
| LIVE/DEAD Cell viability assay | Thermo Fisher Scientific | L3224 |
| FITC-Dextran (2MDa) | Sigma-Aldrich | FD2000S |
| adhesive PCR plate seals | Thermo Fisher Scientific | AB-0558 |
| V-bottom 96-well microplate | Greiner bio-one | 651201 |
| 96-well microplate, Sensoplate Microplate | Greiner-Bio-one | 655892 |
| Pierce™ BCA protein assay | Thermo Fisher Scientific | 23227 |
| Bovine gamma globulin standards | | 23212 |
| Bovine serum albumin (BSA) | Jackson ImmunoReseach | 001-000-162 |
| Neat-008 fluorosurfactant | RAN biotechnologies, Inc. | 008 FluoroSurfactant |
| 1H,1H,2H,2H-Perfluoro-octanol | Sigma -Aldrich | 370533 |
| 1 mL luer-lock plastic syringe | BD | 309628 |
| ½ 23G blunt needle | McMaster-Carr | 75165A255 |

### Supplementary note 1: Intensity Barcoded microparticles (BMPs)

To enable multiplexed detection of secreted proteins at the single cell level, we designed BMPs using fluorescently labelled DNA.[1] Herein, two DNA oligos, one bearing a 5'- biotin and a second a 3'- fluorophore (Fig. S1A), were annealed to generate sets of double-stranded encoder oligos ($dsEO_i$) where *i* correspond to the desired fluorophores. We also included a non-fluorescent oligo, referred to as encoder 0 ($EO_o$, Fig.S1B). The usage of $dsEO_i$ aims to provide similar affinity toward streptavidin (SA) through the biotinylated strand, while also normalizing the molecular footprint of the overall dsEOs on the surface of microparticles, hence mitigating the impact of the size of the fluorophore on the binding onto beads. In addition, specific protein capture can be encoded within the BMPs by supplementing biotinylated cAbs to the dsEOs mixture prior affinity binding onto the SA-coated microparticles (Fig.S1B-C). By mixing $dsEO_i$ and $dsEO_0$ at various proportion, where $dsEO_i + dsEO_0 = 90$ pmol, BMPs exhibiting discrete fluorescence intensity can be generated, hence supporting the synthesis of various cAb-BMPs and empowering high multiplex detection of protein.

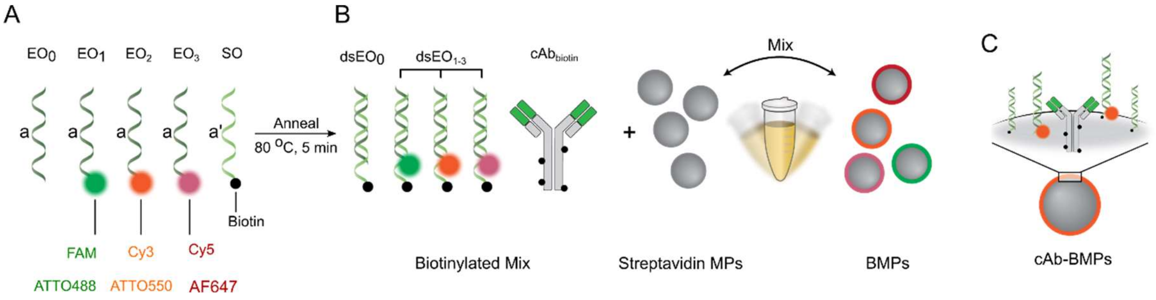

**Figure S1**: Assembly of intensity barcoded microparticles (BMPs). (**A**) Fluorescently labelled encoder oligos ($EO_i$) are individually mixed with a biotinylated oligo (SO) possessing a complementary sequence and annealed to generate sets of double stranded encoder oligos ($dsEO_i$) harboring the fluorescent dye *i*, and a biotin. (**B**) Single-color BMPs are produce by combining, at various mixing ratio of $dsEO_i$:$dsEO_0$, biotinylated reagent and streptavidin-coated microparticles. (**C**) Due to the high affinity of biotin to streptavidin, $cAb_{biotin}$ and dsEOs bind onto the SA-coated microparticles (MPs) to generate capture antibody BMPs (cAb-BMPs).

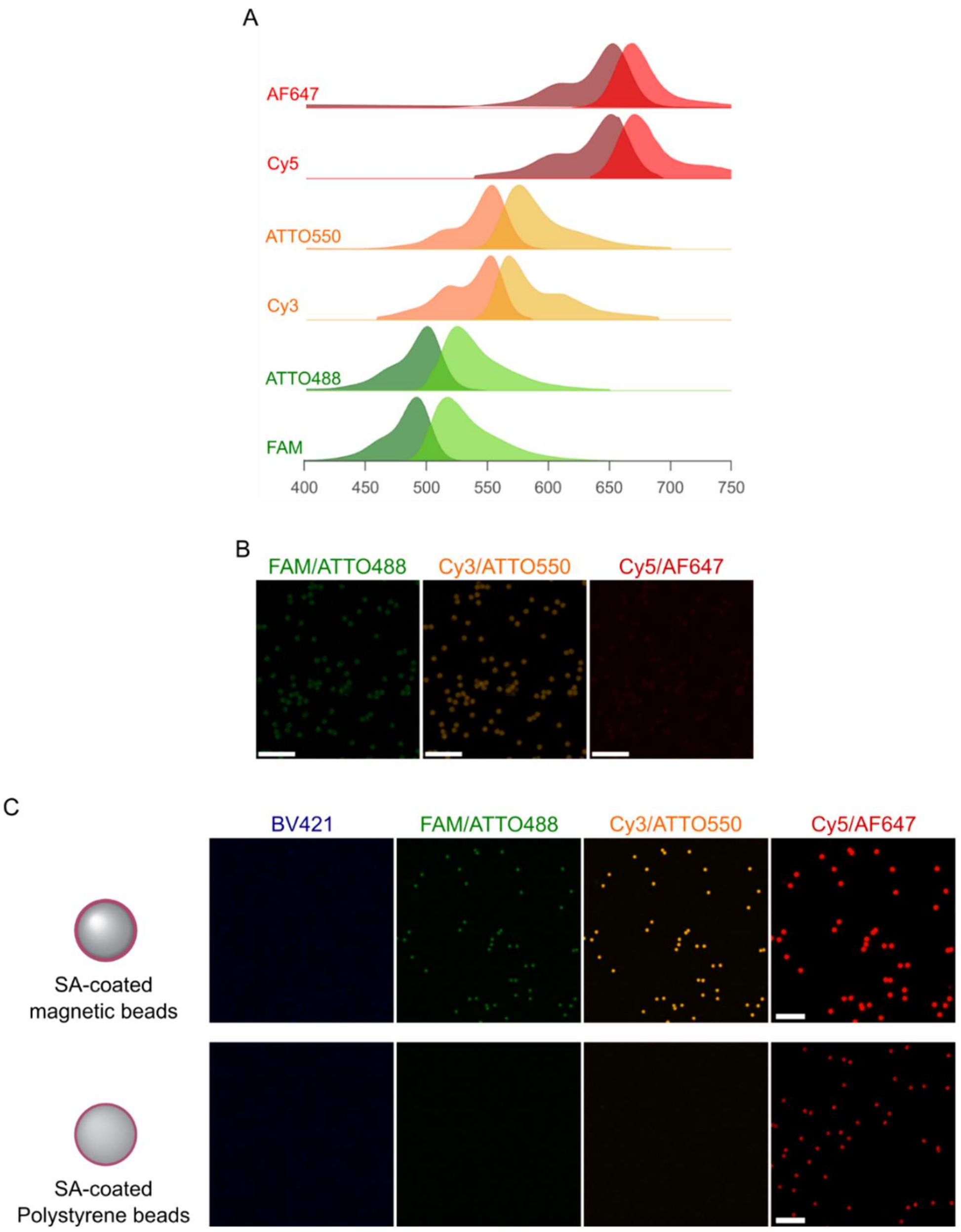

**Figure S2**: (**A**) Excitation and emission spectra of the various fluorescent dyes employed to benchmark the cAb-BMPs whereas FAM, Cy3 and Cy5 correspond to common flow cytometry dyes, while ATTO488-550 and AF647 are dyes exhibiting higher photostability which are favored for microscopy. (**B**) CLSM images of blank cAb-BMPs functionalized with only unlabelled oligos ($dsEO_0$ = 90 pmol) and biotinylated IgG to assess BMPs background in the various Ex/Em channels. Scale bars: 20 µm. (**C**) CLSM images of AF647 labeled-BMPs generated from Streptavidin (SA)-coated superparamagnetic microparticles (e.g. SA-Dynabeads®; Top) or SA-coated polystyrene beads (e.g. Bangs Laboratories, Inc.; Down). Scale bars: 25 µm.

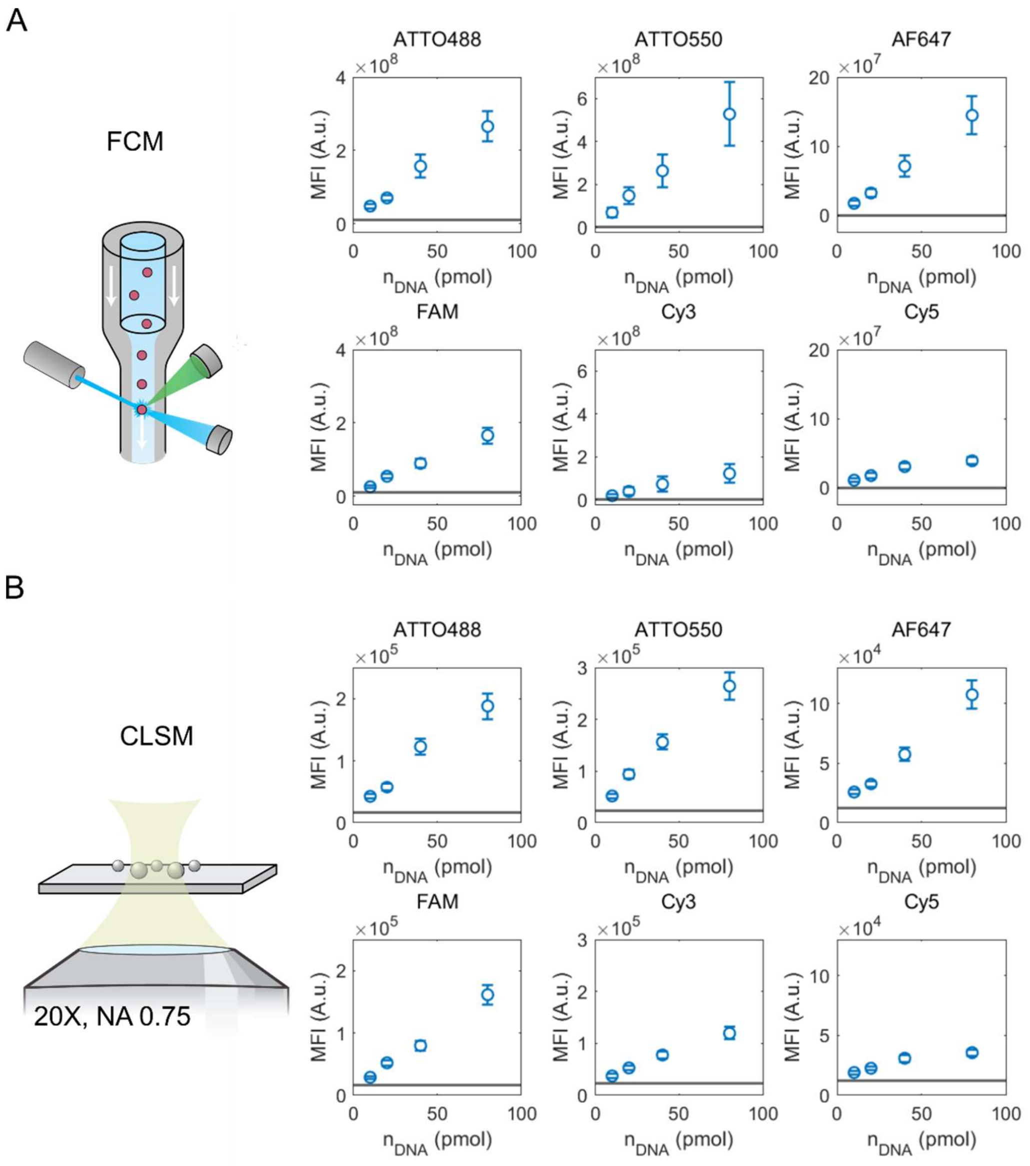

**Figure S3**: Evaluation of various dyes employed on single color intensity barcoded microparticles (BMPs) measured as single particle by (**A**) flow cytometry and (**B**) confocal laser scanning microscopy (CLSM). For CLSM, beads were segmented using signals in the 561/590 Ex/Em channel. Mean ± S.D. of population are presented (*n* = 3). Solid lines correspond to the signal detected on BMPs without labeled oligo, whereas $dsEO_0$ = 90 pmol.

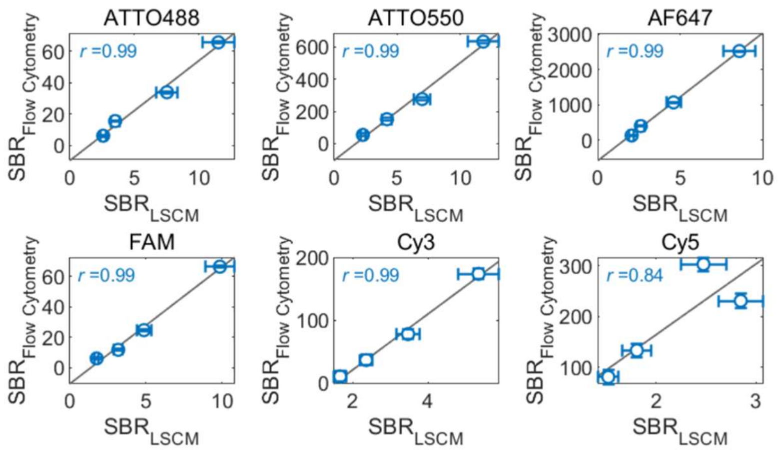

**Figure S4**: Correlation of BMPs signal-to-background ratio (SBR) measured in FCM (**Fig. S3A**) and CLSM (**Fig. S3B**). Fits correspond to a least-square regression. Pearson's r values are shown for each correlation. Mean ± S.D. ($n$ = 3) are presented.

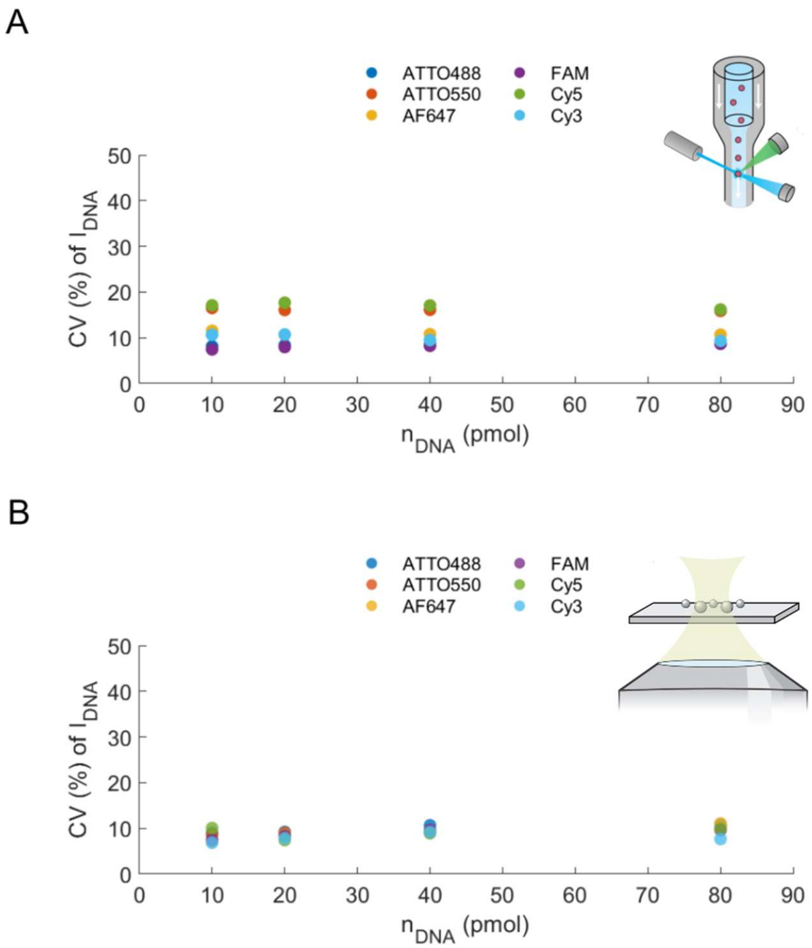

**Figure S5**: Coefficient of variation (CV) of cAb-BMPs generated with various $dsEO_i$ ratios detected by both FCM (**A**) and CLSM (**B**). In all cases, CV were calculated based on the mean and S.D. of triplicate.

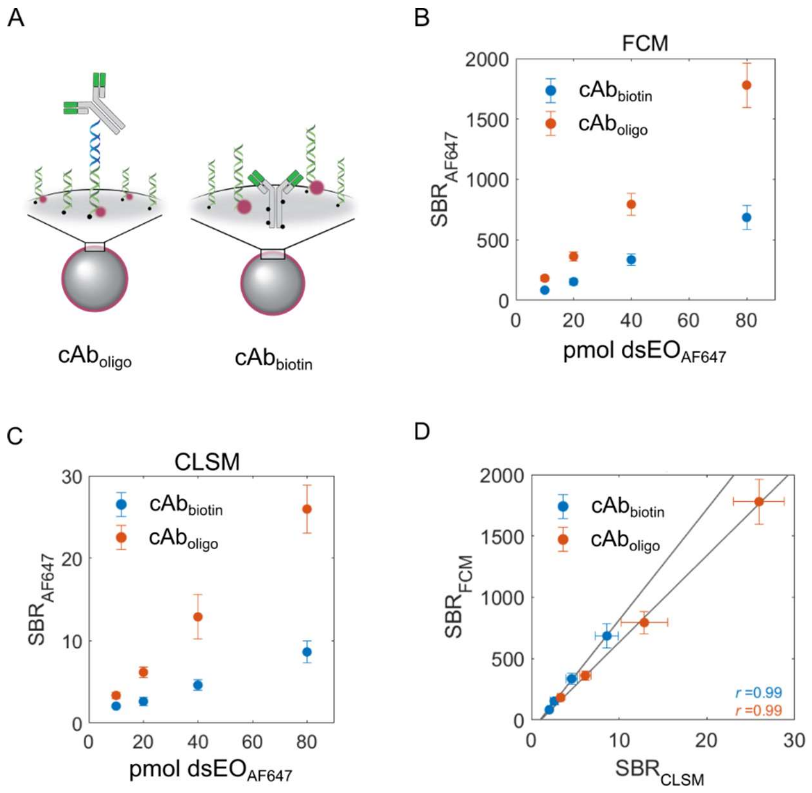

**Figure S6**: Impact of BMPs construct on intensity-based barcoding. (**A**) Scheme presenting the novel ($cAb_{oligo}$) and previously reported construct[1] ($cAb_{biotin}$) relying on the co-adsorption of biotinylated cAb and $dsEO_s$. (**B-C**) Signal-to-background ratio (SBR) of single color (AF647) cAb-BMPs measured by (**B**) FCM and (**C**) CLSM. (**D**) SBR correlation in between cAb-BMPs measured by both analytical methods for the two types of constructs. Lines correspond to linear least-square regression. The presented *r* values correspond to Pearson's coefficient. In all cases, mean ± S.D. ($N$ = 3, n ≥ 5000 BMPs) are presented.

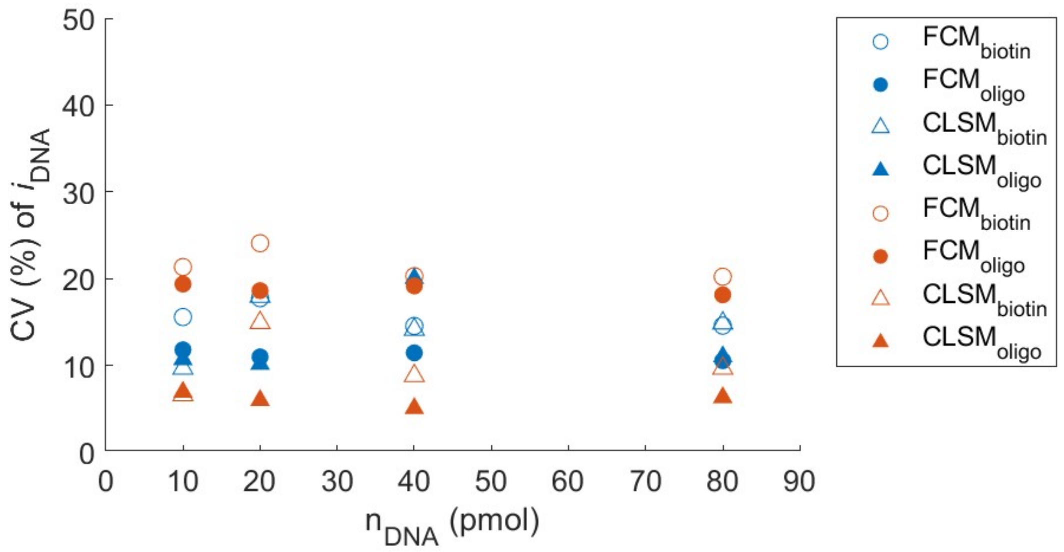

**Figure S7:** Coefficient of variation (CV) of the two cAb-BMPs constructs generated referred to as *biotin* and *oligo* corresponding to the co-adsorption of biotinylated cAb and immobilization of DNA-cAb conjugates via a DNA anchor, respectively. Blue: $MFI_{AF647}$ signal intensity; Orange: ratio $MFI_{AF647}/MFI_{590}$; Open symbol: *Biotin* construct; Filled symbol: *oligo* construct.

**Supplementary note 2: Modeling BMPs position and distribution to guide BMPs design**

By using the single-color cAb-BMPs as standards, we extracted the irradiance (sensitivity) of a dye, corresponding to the slope of the calibration curve providing the relationship between the measured fluorescence intensity and number of labeled $dsEO_i$ adsorbed onto the surface of the MPs. For each BMPs color, we measured an average %CV across all barcodes. By using the averaged %CV, we modeled *in silico* the mean intensity value of a given BMPs and it's expected distribution (~3 folds standard deviation (S.D.)) by using $\%CV = 100^* \sigma/\mu$, where σ corresponds to the standard deviation and μ is the mean of the distribution. This approach helped guide the design of minimally overlapping BMPs by constraining BMPs design to exhibit at least a distance of 3.5-fold their respective S.D. (Fig. S8). In all cases, a maximal number of 90 pmol of total dsEOs was allowed. Using this approach, we model a series of $dsEO_{AF647}$: $dsEO_0$ exhibting minimal overlap for the various construct ($cAb_{oligo}$ and $cAb_{biotin}$) for FCM and CLSM.

Importantly, two metrics will drastically impact the barcoding capacity (i.e. theoretical number of none-overlapping possible barcodes): (i) a high irradiance for a given BMPs (i.e. the slope) and (ii) low %CV across all BMPs. Due to the heteroskedasticity, we opted to apply $1/Y^2$ weighted least-square regression to extract the irradiance of the BMPs on each calibration curve, whereas the error of the fit will ultimately impact the number of potential BMPs. In the case of the %CV, high values will further limit the number of potential BMPs due to large distribution, hence requiring BMPs with large difference in $dsEO_{AF647}$ amount, while been limited to a total amount of 90 pmol of dsEOs.

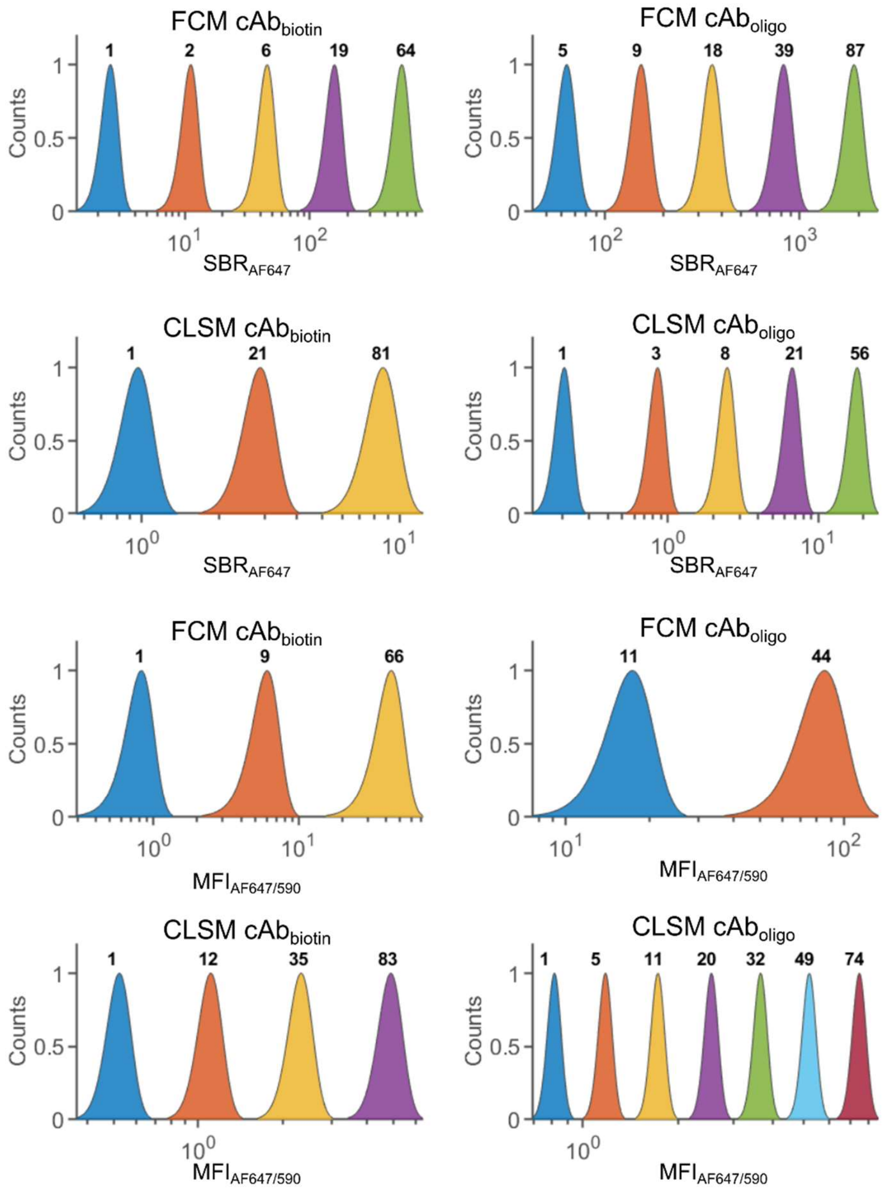

**Figure S8**: Barcode spectral position and bandwidth can be modeled *in silico* to guide barcode optimization. Single color (AF647) barcode position, in either signal-to-background ratio (SBR) or normalized by the scattering signal at 590 nm ($MFI_{AF647/590}$) were predicted by assuming a minimal distance equal to 3.5 folds the expected standard deviation of the barcode position, based on a coefficient of variation (%CV) experimentally calculated on single color standard (Fig. S6). Two structures, wherein biotinylated capture antibody ($cAb_{biotin}$) was co-adsorbed along with the labeled oligos dsEOs onto streptavidin-coated beads, or through the anchoring of the DNA-cAb onto beads ($cAb_{oligo}$) where evaluated. Numbers, located above the predicted Gaussian peak, correspond to the predicted amount, in picomole, of labeled dsEOs to employ during synthesis.

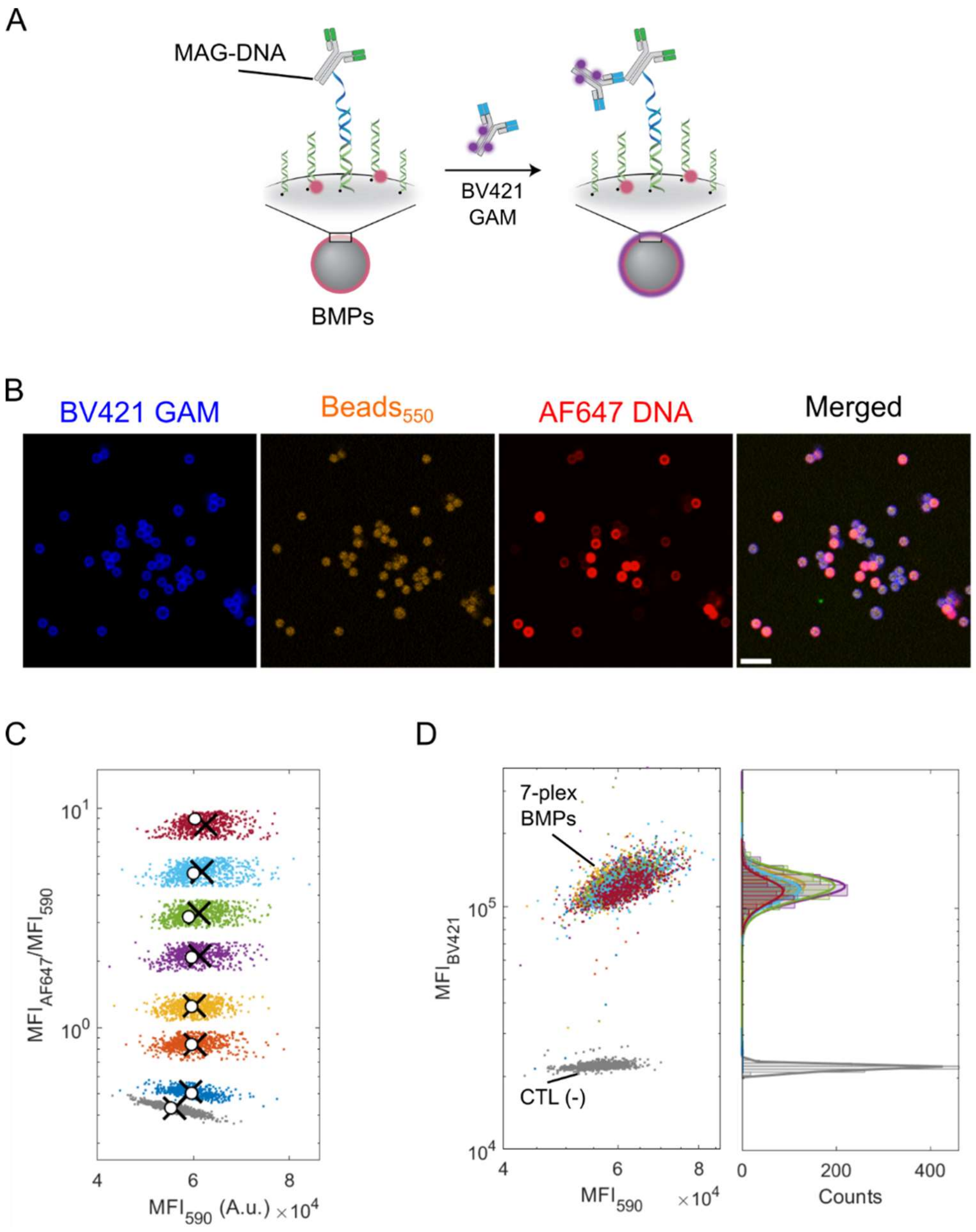

**Figure S9**: DNA-IgG conjugate hybridized onto barcoded microparticles. (**A**) Following BMPs incubation with a single mouse anti-goat DNA-IgG conjugate, the resulting cAb-BMPs are exposed to fluorescently labeled B421 goat anti-mouse IgG detection antibody (dAb) to validate cAb immobilizing onto BMPs. (**B**) Fluorescent CLSM images of the BMPs immobilized on the glass slide. Scale bar: 10 µm (**C**). cAbs-BMPs were analyzed and decoded by Gaussian Mixture Model (GMM). (**D**) BV421 labeled dAb signal across all BMPs possessing 0, 1, 5, 11, 20, 32, 49 and 74 pmol of $dsEO_{AF647}$, respectively. Here, BMPs presenting 0 pmol of AF647 DNA (CTL (-)) were deprived of anchored cAb, acting as negative control.

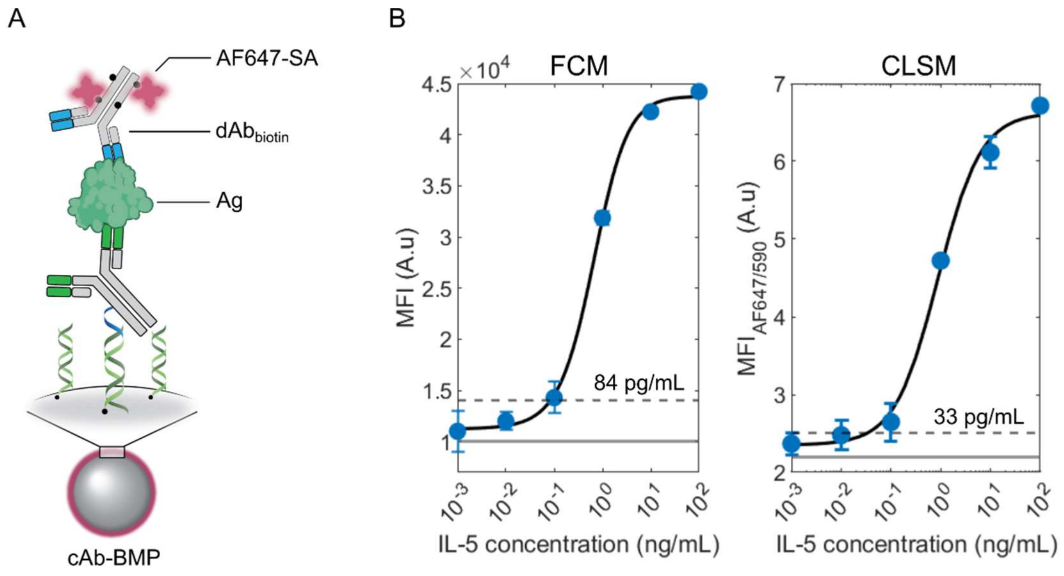

**Figure S10**: Sandwich immunoassay on BMPs with $cAb_{oligo}$ constructs can be read-out by confocal microscopy. (**A**) Scheme of the cAb-BMPs for antigen capture and detection via biotinylated detection antibody ($dAb_{biotin}$) and labeled streptavidin (AF647-SA). (**B**) Binding curves of recombinant IL-5 captured by cAb-BMPs measured by FCM (left) and CLSM (right). Error bars are mean ± S.D. (*n* = 3). Solid grey lines are the mean of a blank and dashed lines are the calculated limit of detection of IL-5. Fits correspond to a four-point logistic regression.

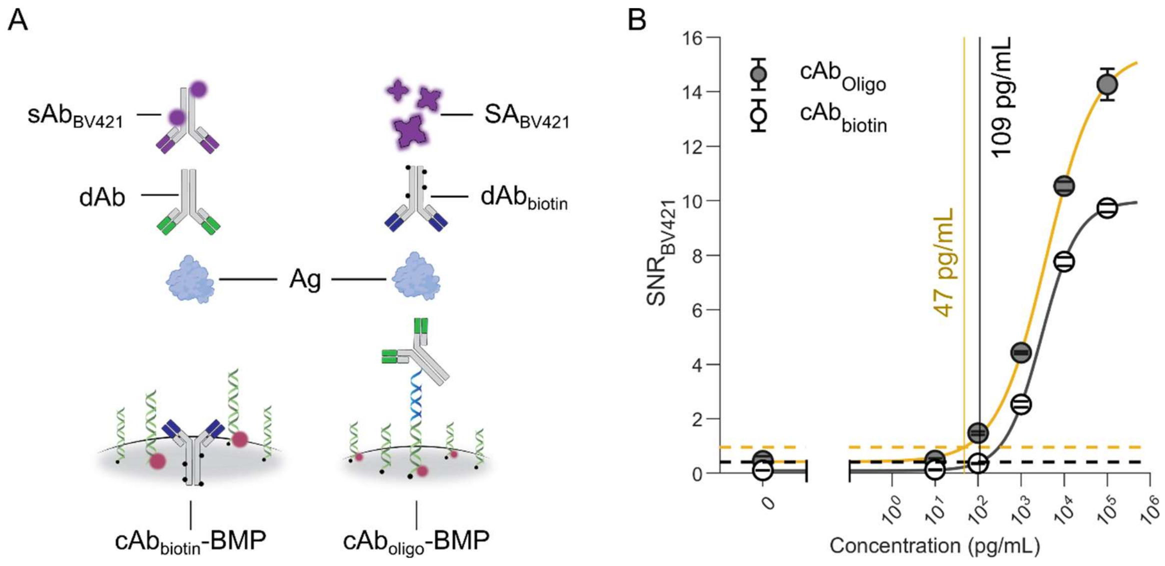

**Figure S11:** Antibody encoding of BMPs using $cAb_{oligo}$ improved assay performance compared to $cAb_{biotin}$ using the same antibody pairs. (**A**) Schematics presenting the key differences between the two BMP constructs: (Left) $cAb_{biotin}$ rely on biotinylated cAb adsorbed onto BMPs during self-assembly to enable antigen (Ag) capture. A single dAb (mouse IgG) is used for detection and assay signal is generated using BV421-labeled goat anti-mouse IgG as secondary antibody ($sAb_{BV421}$). (Right) Conversely, the $cAb_{oligo}$ rely on the anchoring of a oligo-conjugated cAb for Ag capture, followed by the usage of the biotinylated dAb for detection. Assay signal is generated via the usage of BV421-labeled streptavidin ($SA_{BV421}$). (**B**) Binding curves of $cAb_{oligo}$ and $cAb_{biotin}$ detecting IFN-γ in buffer. Mean ± S.D. are presented ($n = 3$). Fit curves correspond to a 4-point logistic regression. The horizontal dashed lines correspond to the calculated signal at LOD, calculated as 3-fold the standard deviation of the blank sample above background. Vertical lines mark the calculated LOD.

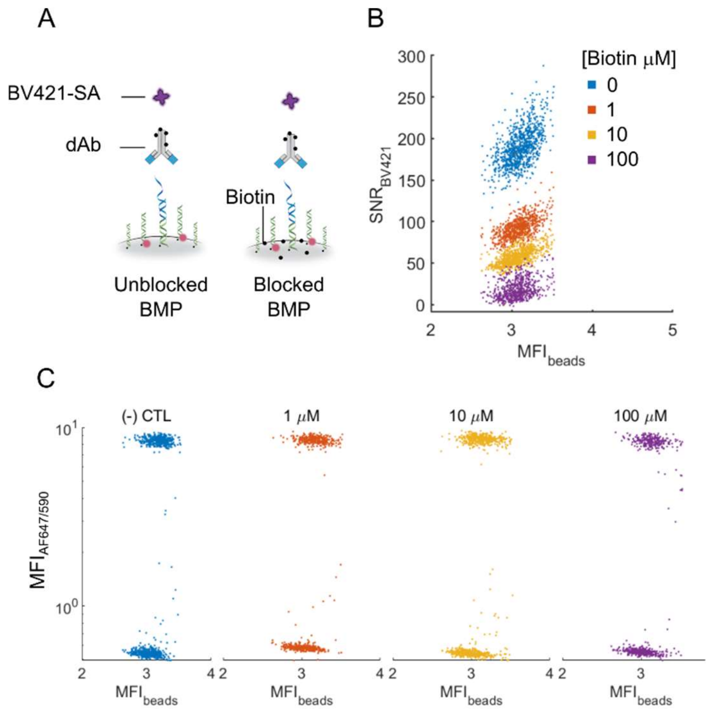

**Figure S12:** Blocking beads with biotin reduces undesired binding of biotinylated detection antibody (dAb) on streptavidin coated beads in a sandwich immunoassay. (**A**) Scheme presenting the surface of BMPs coated by streptavidin. 'Unblocked' BMPs are sequentially incubated with dAb and detected by BV421 labeled streptavidin (BV421-SA). 'Blocked' BMPs are first exposed to free biotin that could bind unreacted streptavidin prior to incubation with dAb and BV421-SA. (**B**) Signal-to-noise ratio (SNR) of BV421-SA measured by CLSM on BMPs incubated with 0, 1, 10 or 100 μM of free biotin supplemented into the blocking buffer (PBST0.1% + 1% wt/vol BSA). (**C**) Scatter plots showing the fluorescent barcodes are unaffected by incubation with free biotin between 0 to 100 μM, revealing that barcodes are robust in upon biotin incubation for all ratios of $dsEO_{AF647}$:$dsEO_0$.

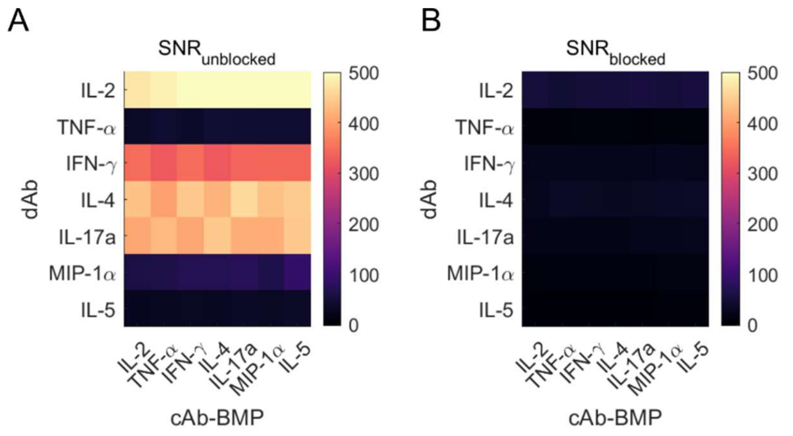

**Figure S13:** Blocking remaining streptavidin binding sites on BMPs reduces undesired assay background in an antibody dependant fashion. Signal-to-noise ratio (SNRs) of BMPs anchoring a single capture antibody (cAb-BMP) per barcode without blocking (**A**) and with blocking (**B**) with biotin prior assay measured by CLSM. A mixture of BMPs, each with their respective cAb, were exposed to a single biotinylated dAb (rows). The mean MFI for each decoded cAb-BMPs is then reported (column).

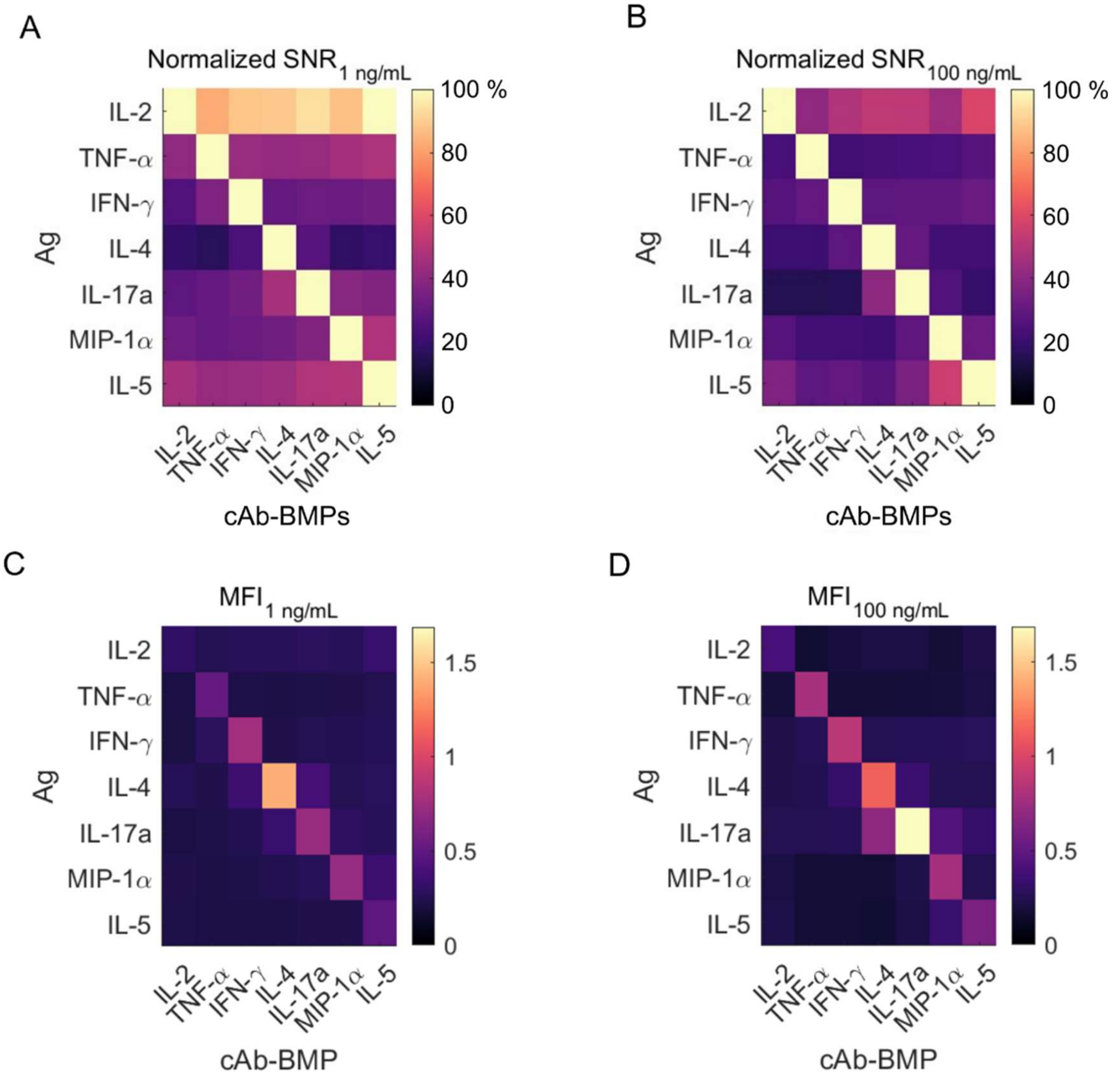

**Figure S14**: Specificity matrix of specific (diagonal) and non-specific (off-diagonal) normalized assay $SNR_{Ag}$ (A, B) and assay signal $MFI_{Ag}$ (C, D) following individual cytokines (Ags; rows) at (**Left**) 1 ng/mL and (**Right**) 100 ng/mL for each cAb-BMP (columns). The $SNR_{Ag}$ were calculated by correcting the mean $MFI_{Ag}$ presented in C and D by the mean assay background ($n$ = 3), normalized by the global standard deviation ($n$ = 18) and then by the $SNR_{Ag}$ value of the corresponding Ag on the diagonal. $MFI_{Ag}$ were calculated by correcting the mean MFI by the mean assay background ($n$ = 3).

We incubated all BMPs with a single recombinant antigen, followed by a cocktail containing all dAbs, and then BV421-labeled SA. We observed low cross reactivity for both low (1 ng/mL; Fig. S14A, C) and high (100 ng/mL; Fig. S14B, D) antigen concentration, with the exception of IL-2 at low concentration, thus validating this panel of antibody pairs for MSA. The apparent cross reactivity (in normalized SNR) of recombinant IL-2 is due in part to the low signal at low concentration (Fig. S14A,C) compared to the other Ab pairs employed within our antibody panel, and a low SNR and comparatively high background signal normalized by specific (diagonal) assay signals. At higher concentration, the SNR is higher (Fig. S14B, D), suggesting its functionality within its assay range. Future studies may seek to identify reagent combinations that yield higher signals to improve the sensitivity against IL-2.

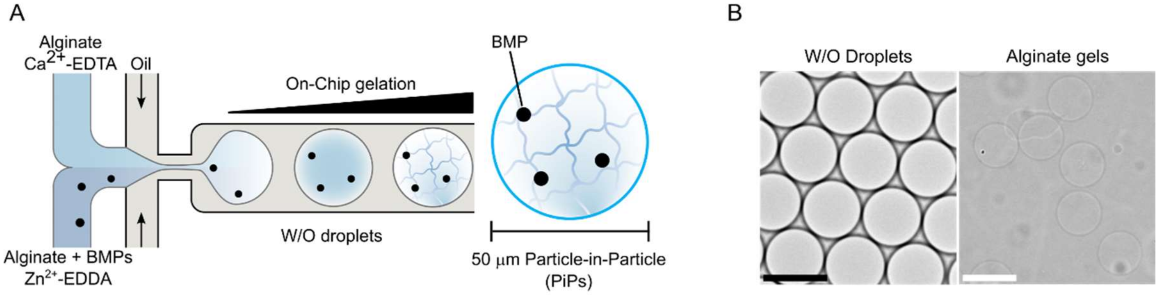

**Figure S15**: Alginate hydrogel-based PiPs generation. (**A**) W/O droplets encapsulating soluble alginate are generated by microfluidics. Alginate is ionically cross-linked on-chip through competitive ligand exchange,[2] following droplet generation. Generated PiPs are collected and resuspended in buffer after droplet coalescence. (**B**) Brightfield images of W/O droplets and alginate hydrogels. Scale bars: 50 μm.

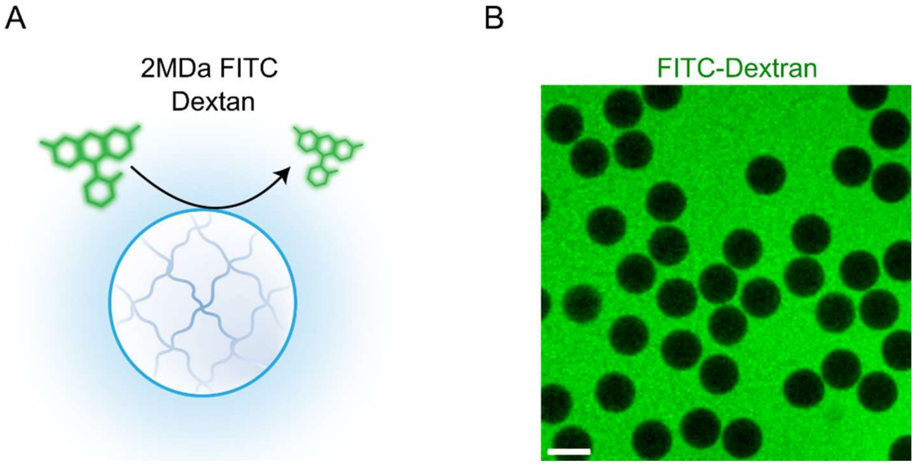

**Figure S16**: Alginate hydrogel can be localized by labeling the outside aqueous phase using large molecular weight dextran. (**A**) Schematic describing the minimal diffusion of 2MDa FITC-labeled dextran inside the alginate gels or PiPs. (**B**) CLSM image of PIPs 2MDa FITC-labeled buffer. The middle plan is presented. Scale bar: 50 μm.

**Supplementary note 3: PiPs are permeable to commonly used immunoassay reagents**

To assess PiPs permeability, we produced 50-μm-diameter PiPs and trapped them in a microchannel possessing a step height reduction from 60 μm to 20 μm, that was manufactured by PDMS replica molding of 3D-printed molds (Fig. S17A). We perfused 2MDa FITC-dextran that is too large to diffuse throught the PiPs and thus remained outside, which increased the contrast in between the boundaries of the hydrogels, and the surrounding medium. This contrast-enhanced image aided to localize PiPs by fluorescence microscopy as fluorescent-negative objects by automatic image processing. However, PiPs deformed upon trapping and could even squeeze through the 20-μm-deep channels. We coated the microchannels with positively charged poly-L-lysine to improve the immobilization of the negatively charged alginate, but it remained ineffective. Suprisingly, we were able to immobilize them by supplementing 2 mM of $Ca^{2+}$ ions in the perfusion buffer and incubating them for 5 min inside the trap, in absence of flow. Following this short incubation, PiPs remained immobilized upon perfusion at various flow rates (100 μL/hr and 1000 μL/hr; Movie S1) and also through the perfusion of various subsequent buffers supplemented with $Ca^{2+}$ (Movie S2). We hypothesized that $Ca^{2+}$ ions may leach out from the alginate matrix upon buffer perfusion, fragilizing the outermost layer of the hydrogels causing it to peel-off the surface subjected to flow in absence of $Ca^{2+}$. Further experiments may be undertaken in the future to better characterize this behavior but are currently outside the scope of this study.

We opted to test the permeability of the PiPs for biomolecules with varying molecular weight typically used in immunoassay. We used MPs functionalized with goat anti mouse Ab to generate PiPs and entrapped them within a microchannel (Fig. S17B). Following the immobilization of PiPs, we perfused a $Ca^{2+}$ supplemented buffer containing BV421-labeled SA, AF488-labeled chicken anti-goat antibody and 2MDa FITC-dextran (Fig. S17C). These dyes were selected to confirm diffusivity of large polymer-based dye such as Brilliant violet® (BV421) and smaller organic dye (AF488) inside the alginate. We observed a rapid increase in fluorescence of the MPs for both BV421 and AF488/FITC channel consistent with diffusion of both BV421-SA and AF488-Ab into the alginate matrix and binding to the MPs (Fig. S17D), while excluding large 2MDA FITC-dextran from the PiPs. We measured a higher flux of ~150 kDa Ab onto the beads compared to the 340 kDA SA (Fig. S17E) via time course analysis of the fluorescence signal of the beads, which is consistent with the decreasing diffusion constant of larger molecules. We calculated the characteristic diffusion time *t* for reaching half of the maximal fluorescence for AF488-labeled Ab and BV421-labeled SA as of 3.6 and 10.1 min, respectively. Importantly, we noted selective binding of both Ab and SA on the MPs with minimal background within the hydrogels and an overall higher irradiance for BV421-SA compared AF488-Ab. These observations confirmed that the PiPs are permeable to fluorescently-labelled Ab and SA and could be used for immunoassays.

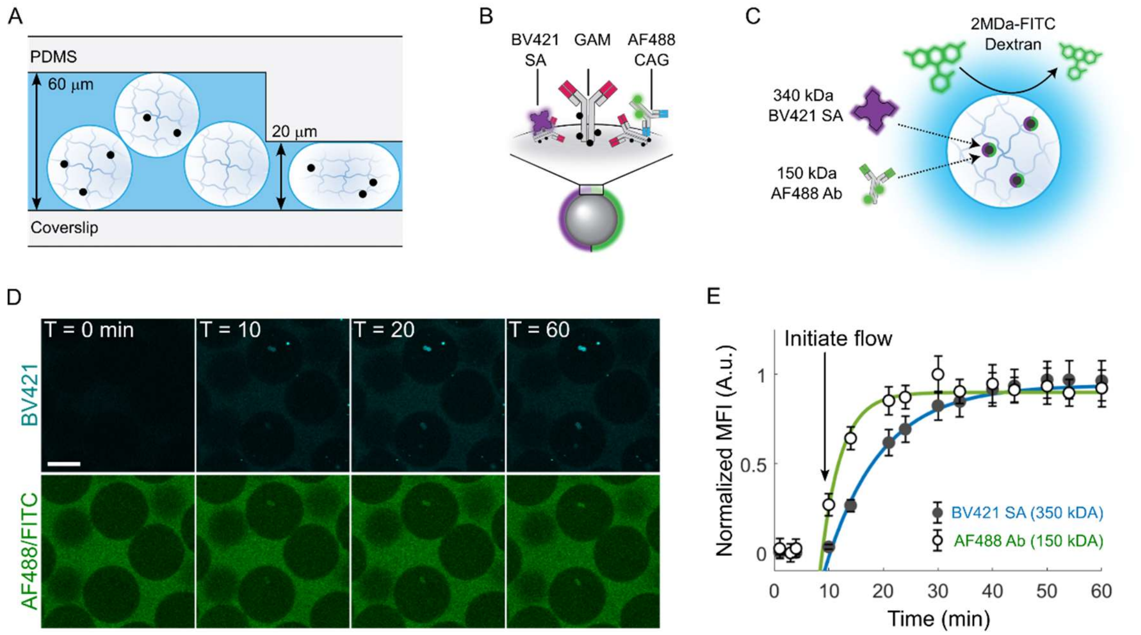

**Figure S17**: Alginate-based PiPs are selectively permeable for immunoassay reagents but not for biomolecules having MW ≥ 2MDa. (**A**) Schematic presenting the entrapment of PiPs in microfluidic traps (60 µm height) by an abrupt reduction in channel height (20 µm) (**B**) Schematic presenting the adsorption of biotinylated Goat anti-Mouse (GAM) antibodies on SA-coated MPs and detected by of BV421 labeled streptavidin (SA) and AF488 labeled Chicken anti-Goat (CAG) antibodies. (**C**) Schematic of size selective diffusion of biomolecules throughout PiPs. (**D**) CLSM images of PiPs exposed to 2 µg/mL BV421-SA, 4 µg/mL AF488-CAG and 1 nM 2MDa-FITC dextran. Scale bar: 50 µm. (**E**) Normalized mean fluorescence intensity (MFI) of labeled SA and CAG measured on MPs upon perfusion. Mean ± S.D. (*n* = 5, > 200 MPs measured per replicate). Fit corresponds to a 1D accumulation model, defined as $f(t) = A(1-exp(-(t_0-\lambda)/\tau)$ where $A$, $\lambda$ and $\tau$ are fit parameters and $t_0$ is the time offset to account for the delayed perfusion of SA and Ab. The fitted value of $\tau$ corresponds to the characteristic diffusion constant.

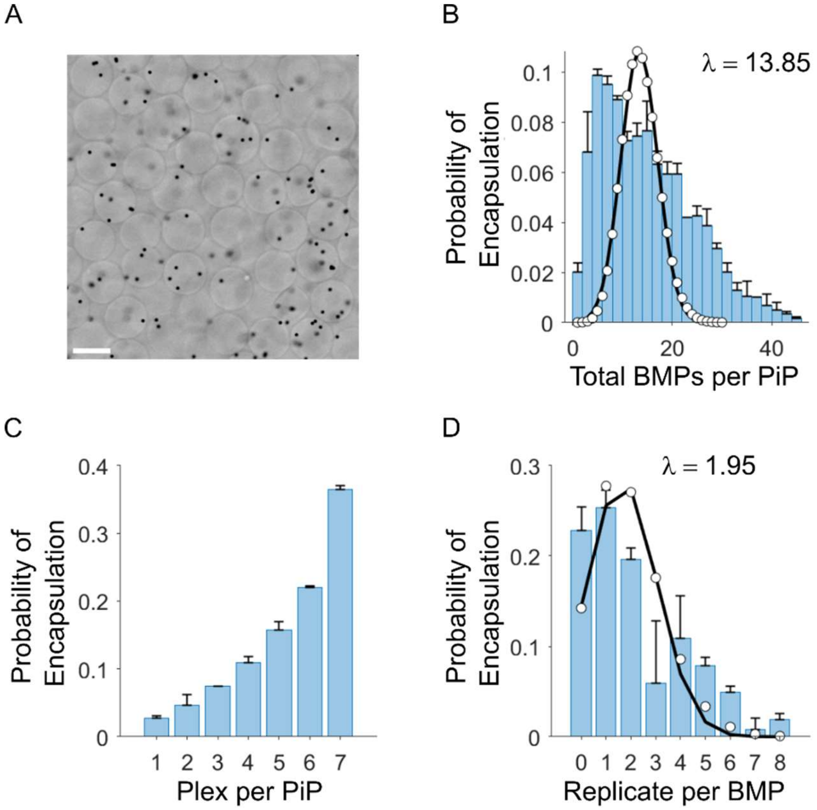

**Figure S18**: Statistic of antibody-encoded barcoded microparticles (cAb-BMPs) encapsulation in alginate hydrogel to generate the particles-in-particle system (PiPs) via droplet-based microfluidics. (**A**) Brightfield microscope image of PiPs entrapping cAb-BMPs (30 000 BMPs/µL per barcode, 210 000 BMPs/µL overvall). Scale bar: 50 µm. (**B**) Probability of encapsulation of given (total) number of BMPs per PiP for an average number of BMPs per bead λ = 13.85. Lambda was calculated based on 210x10$^6$ beads per mL partitioned into 50 µm droplets (~65 pL). Scatter points correspond to the predicted Poisson statistic. Mean ± S.D. (N = 2, *n = 5*) are presented. (**C**) Probability of detecting unique cAb-BMP (i.e. plex) per PiP. Mean ± S.D. (N = 2, *n* = *5*) are presented. (**D**) Mean number of BMPs with the same barcode (i.e. technical replicate) co-entrap within a single PiP. Scatter points correspond to the Poisson statistic based on the experimental λ value of 1.95, calculated from an individual cAb-BMP concentration of 30x10$^6$ BMPs per mL encapsulated into 50 µm droplet (~65 pL). Mean ± S.D. (N = 2, *n* = *7*).

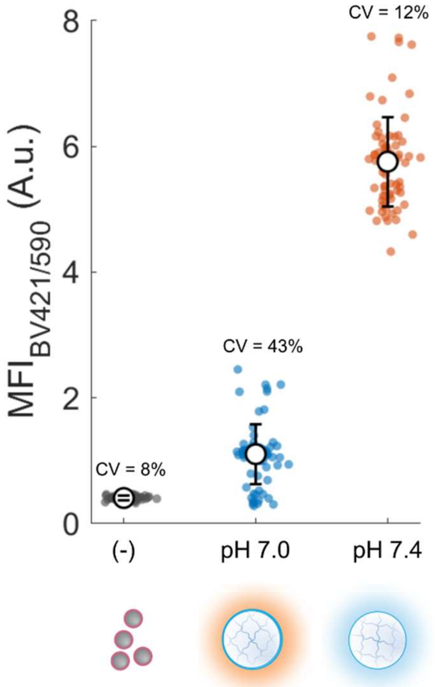

**Figure S19**: pH affects the antigen capture on the cAb-BMPs. Scatter plot presenting assay signal (MFI$_{BV421/590}$) of BV421-labeled SA detection of anti-human IL-17a exposed to beads in entrapped in PiPs at various pH. The negative control (-) corresponds to the assay signal of beads suspended in solution in absence of antigens at pH 7.4 (grey points). Recombinant IL-17A was encapsulated at 100 ng/mL along with anti-human IL-17A BMPs and soluble alginate at pH 7.0 and incubated for 2 h for both antigen capture and alginate gelation by ionic cross-linking prior droplet coalescence, and detection by sandwich immunoassay with dAb at pH 7.4 (blue points). PiPs entrapping anti-human IL-17A BMPs were buffer exchange at pH 7.4 prior antigen capture, incubated with 100 ng/mL IL-17A at pH 7.4 for 2 h and detected by sandwich immunoassay using dAb at pH 7.4. Antigen concentration was adjusted to account for both the dilution by hydrogels, and the dilution during droplet generation.

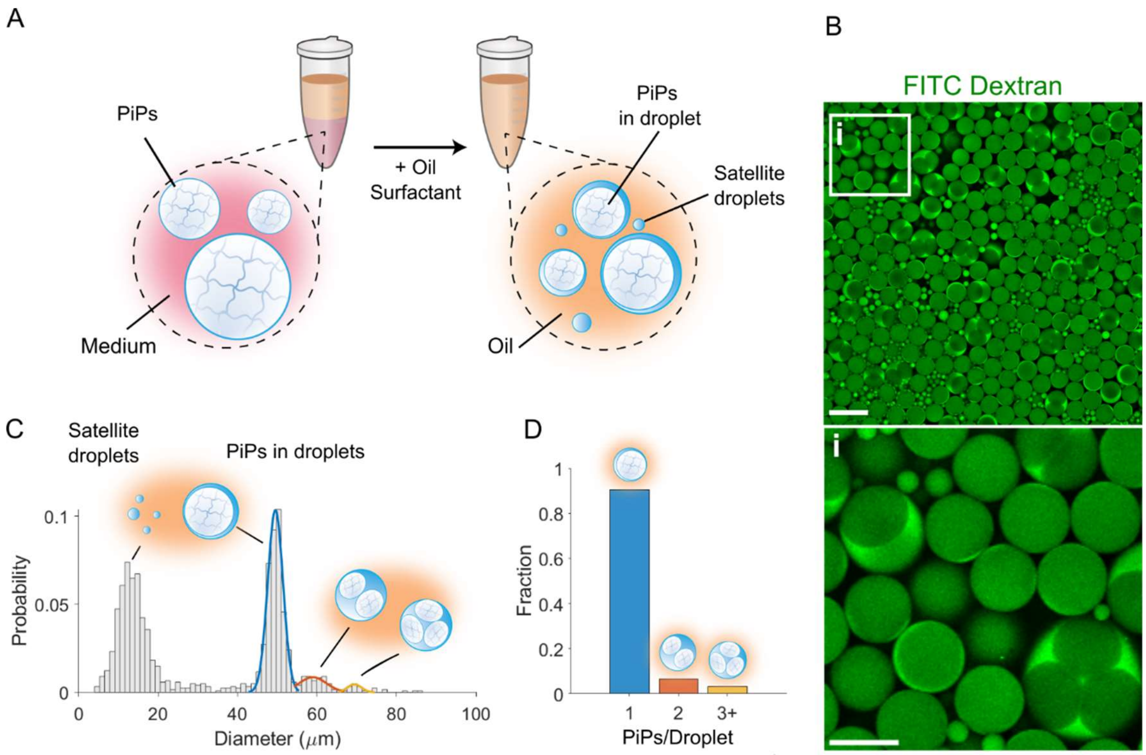

**Figure S20:** PiPs can be washed and re-entrapped into W/O droplet for downstream incubation. Upon adding oil and surfactant to a suspension of PiPs and rapidly pipetting up and down for 1 min, PiPs can be segregated into aqueous droplets. (**A**) PiPs loaded with excessive crosslinking reagents are washed with buffer or medium for removing reagent and promoting nutrient perfusion. PiPs are re-emulsified by pipetting via the addition of a fluorinated oil supplemented with 2% w/w fluorosurfactant. (**B**) CLSM images of PiPs in droplets. For ease of visualisation, 500 kDa FITC-dextran was supplemented to the aqueous phase. Scale bars are (top) 50 and (inlet *i*) 100 μm, respectively. (**C**) Size distribution of satellite droplets, single, and multiplex PiPs-loaded droplets. Fits represent gaussian distributions of the gated droplet size. (**D**) Fraction of PiPs in droplet containing 1, 2 or more PiPs per droplet.

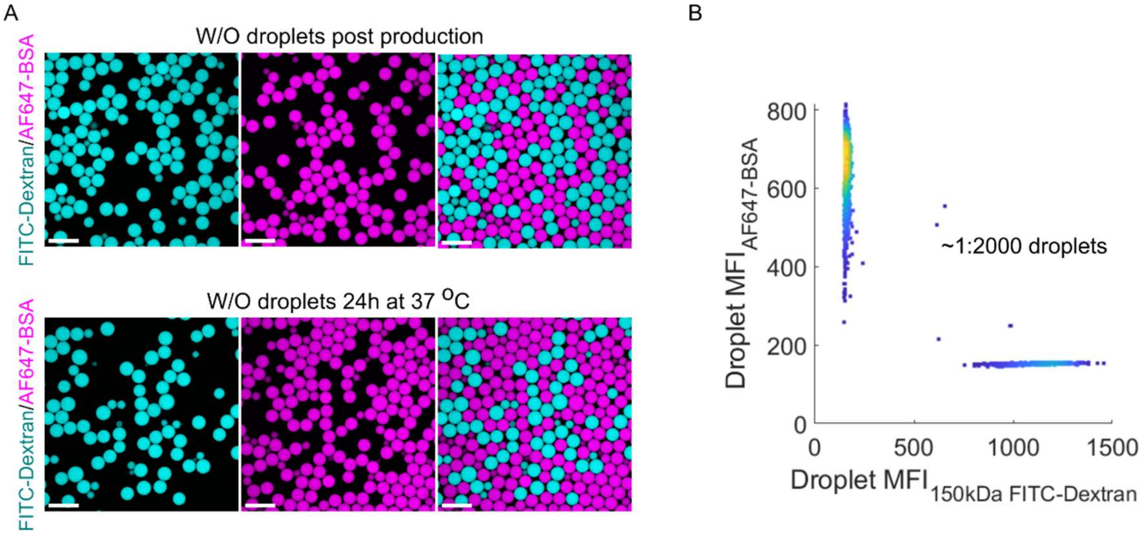

**Figure S21**: Assessment of droplet leakage. (A) CLSM images of water-in-oil (W/O) droplets encapsulating either 150kDa FITC-dextran, or AF647-labeled BSA in assay buffer composed of phenol-red free RPMI 1640 supplemented with 10% (v/v) heat inactivated FBS, 1% (v/v) penicillin/streptomycin, 25 mM HEPES pH 7.4 and 2 mM $CaCl_2$ directly after droplet production (Top) or following incubation at 37 °C for 18 h (Bottom). Scale bars, 100 µm. (B) Fluorescent droplet signals. Droplet signals were normalized by the droplet area. Out of focus droplets were removed by applying a threshold on the detected area. Around 1:2000 droplets showed significant signal in both fluorescent channel (FITC/AF647).

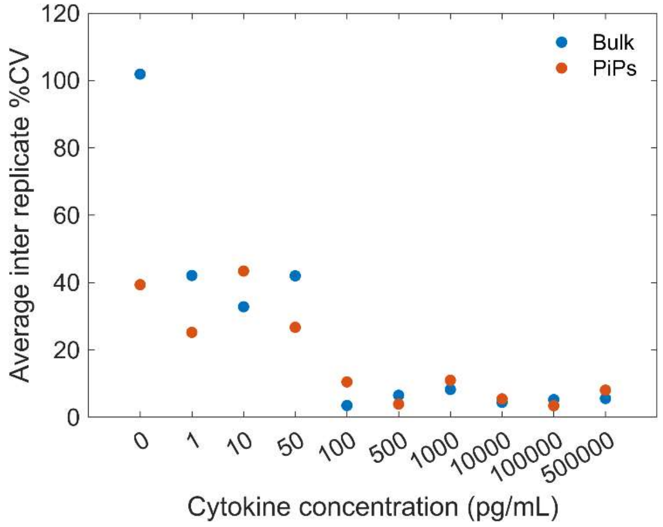

**Figure S22**: Average %CV between replicates for multiplex sandwich immunoassay performed using bulk BMPs or PiPs. The %CV was calculated from the standard deviation and mean from each assay results (*n* = *5*).

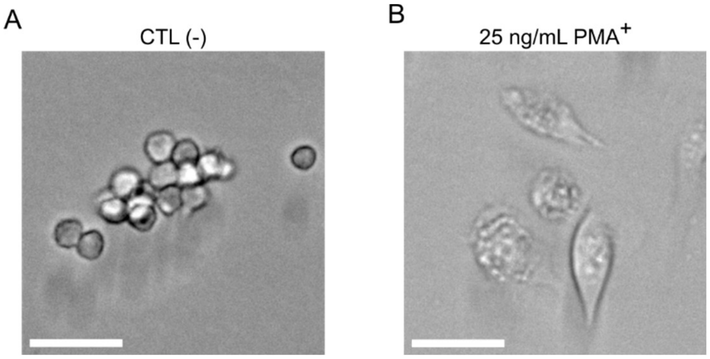

**Figure S23**: Differentiation of THP-1 cells by via Phorbol 12-myristate 13-acetate (PMA). (**A**) THP-1 cells prior and (**B**) after exposure to 25 ng/mL PMA for 48 h. Scale bars: 50 µm.

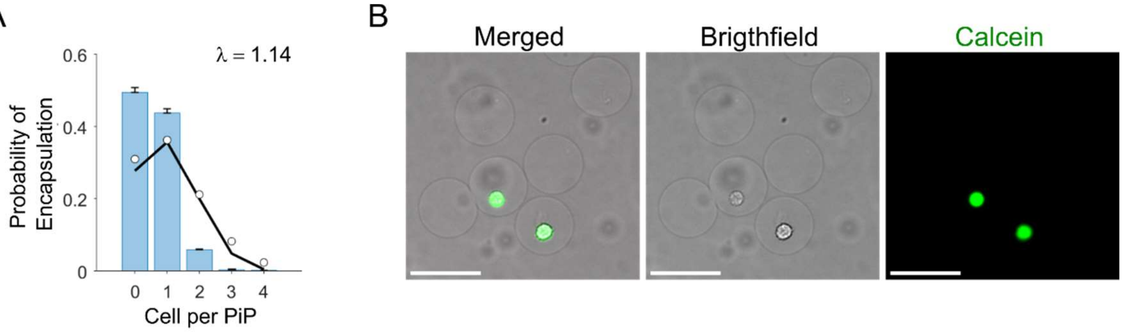

**Figure S24**: Single cell encapsulation into alginate hydrogels by droplet-based microfluidics. (**A**) Probability of encapsulation of none, single and multiple cells per gel imaged by CLSM. Scatter points correspond to the predicted Poisson statistic from experimental λ value of 1.14 whereas $18x10^6$ cells per mL were segregated into 50 µm (~65 pL) droplets. Mean ± S.D. (*N* = 2, *n* = 5) are presented. (**B**) Widefield microscope images of calcein-AM stained viable THP-1 cells. Scale bar: 50 µm.

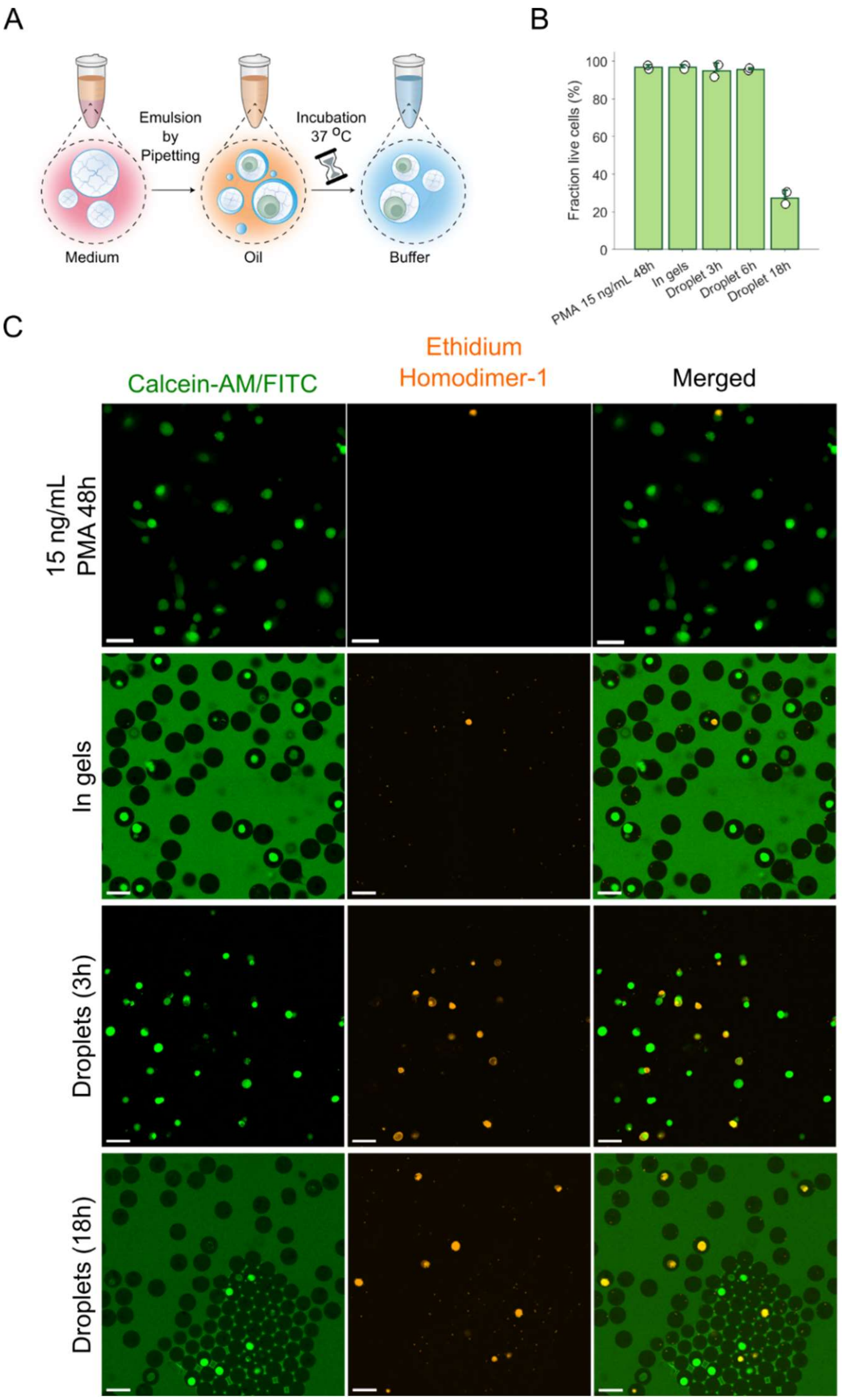

**Figure S25**: Cell viability of differentiated THP-1 cells entrapped in PiPs (**A**) schematic presenting PiPs, rinsed with medium and re-emulsified into W/O droplet to generate the PiPs-in-droplet array for incubation at 37 °C Following incubation, droplets were coalesced, and cell stained with calcein-AM (Live) and ethidium homodimer-1 (Dead). (**B**) Cell viability of THP-1 entrapped in PiPs. Cell exhibiting both signal in calcein (Live) and

Ethidium Homodimer-1 (Dead) were considered as viable. Mean ± S.D. are presented (*N* = 2). (**C**) CLSM images PMA differentiated THP-1 cell prior being formulated into PiP, and PiPs imaged after being incubated into W/O droplets for 0 (in gel), 3h and 18h at 37 $^{o}$C. Scale bars: 50 µm. Note that small points in the ethidium homodimer-1 (Dead) channel correspond to the BMPs. As dead stain typically labels DNA, the BMPs will exhibit a significant rise in fluorescence in the 561/590 nm channel.

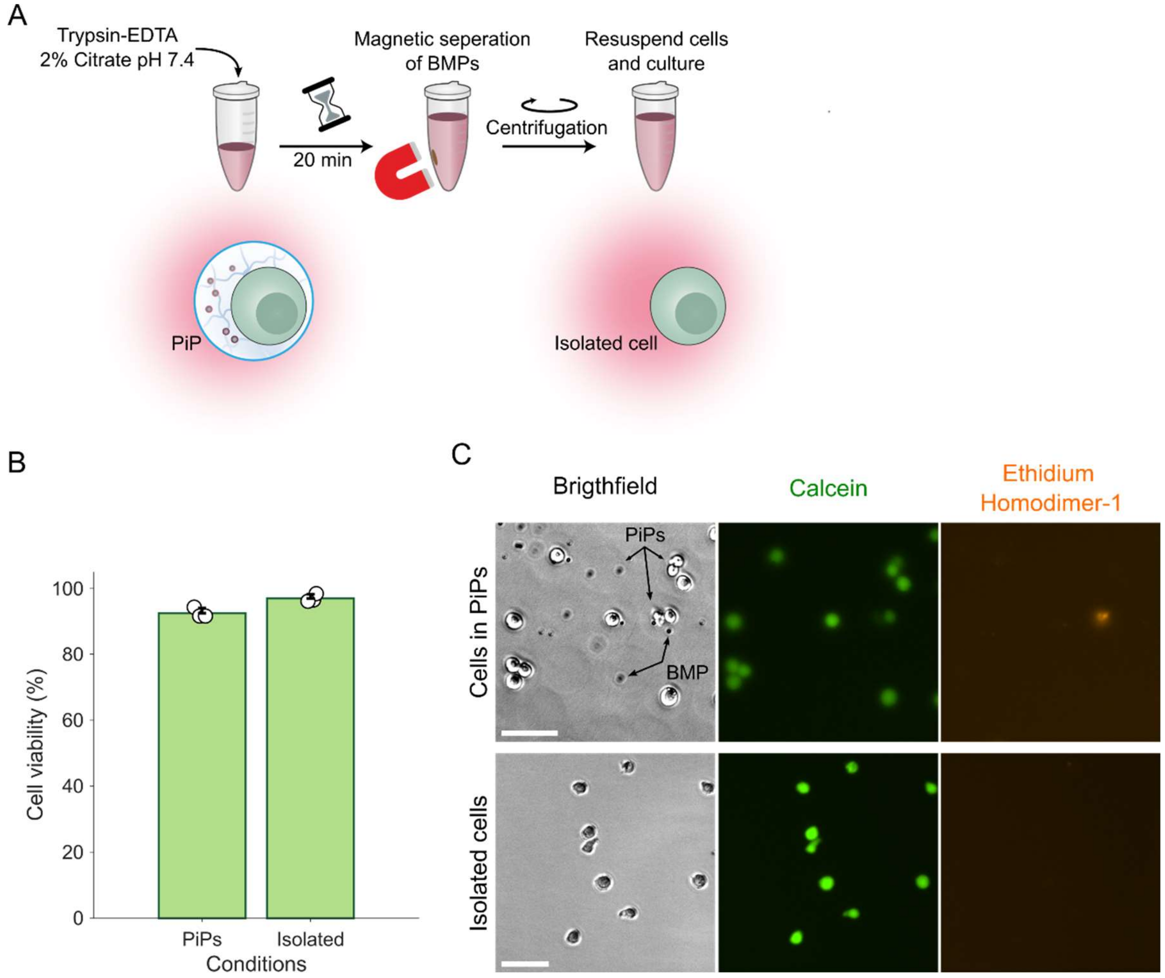

**Figure S26**: Live cells can be extracted from PiPs and cultured. (**A**) General pipeline to extract cells from alginate-based PiPs. First, potential cell adhesion to alginate is cleaved by suspending hydrogels into trypsin-EDTA. Ionically cross-linked alginate hydrogels are degraded by adding 2% citrate to chelate the $Ca^{2+}$ ions. Then, BMPs are magnetically separated from the mixture, and cells isolated via centrifugation. (**B**) Viability of THP-1 cells within PiPs and after being extracted from the PiPs. (**C**) Fluorescence microscopy images of PiPs entrapping BMPs and THP-1 cells (top) and isolated from PiPs using the procedure described in A (Bottom). Scale bars: 50 μm.

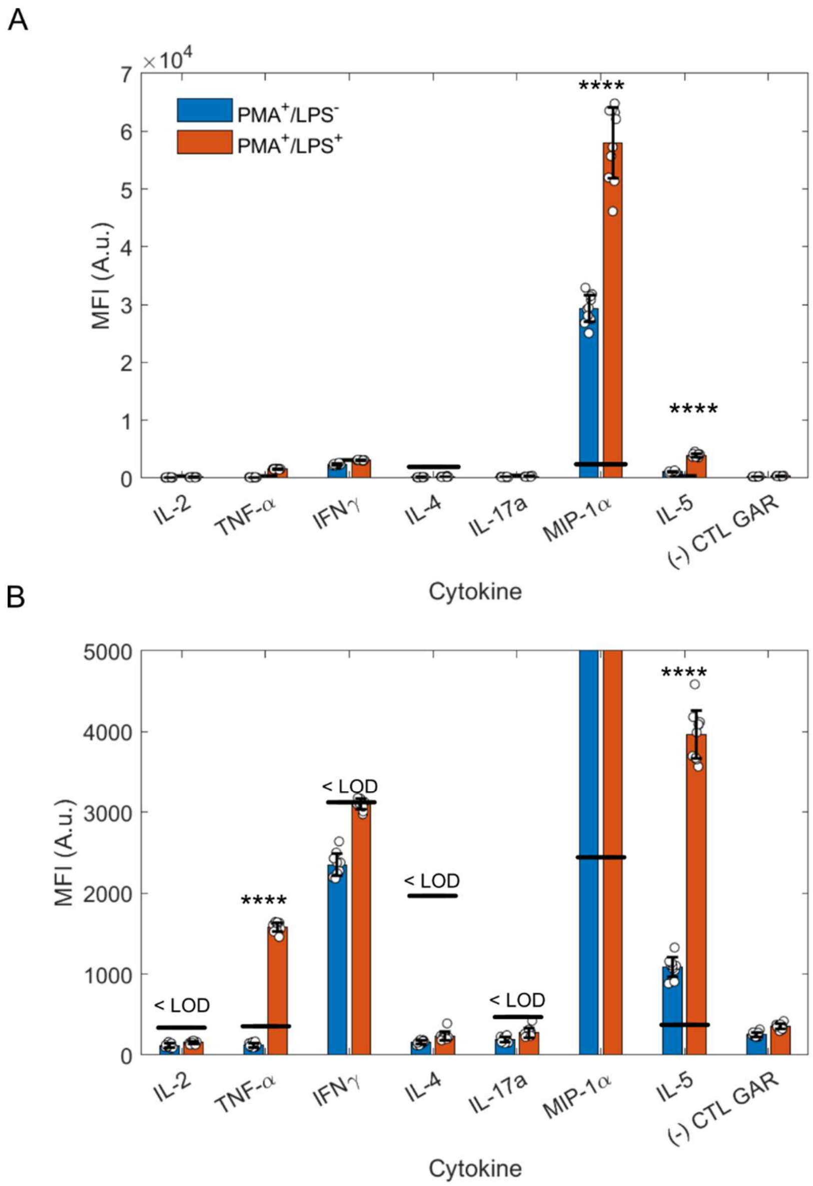

**Figure S27**: (**A**) Multiplexed cytokine detection by antibody microarray of differentiated THP-1 cell supernatant with and without 100 ng/mL LPS stimulation. (-) CTL GAR: Goat-Anti Rabbit capture IgG. Mean ± S.D. ($n$ = 10) are presented. (**B**) Zoomed bar graph from (**A**) highlighting signals measured between 0 and 5000 (A.U.). Solid black lines represent the calculated LOD for each cytokine from their corresponding binding curve in assay buffer (25 mM MOPS pH 7.4, 130 mM NaCl, 2 mM $CaCl_2$, 2% BSA). Statistical test represents unpaired, two-tailed $t$-test in between $LPS^-$ and $LPS^+$. <LOD: lower than limit of detection; ****: $P<0.0001$.

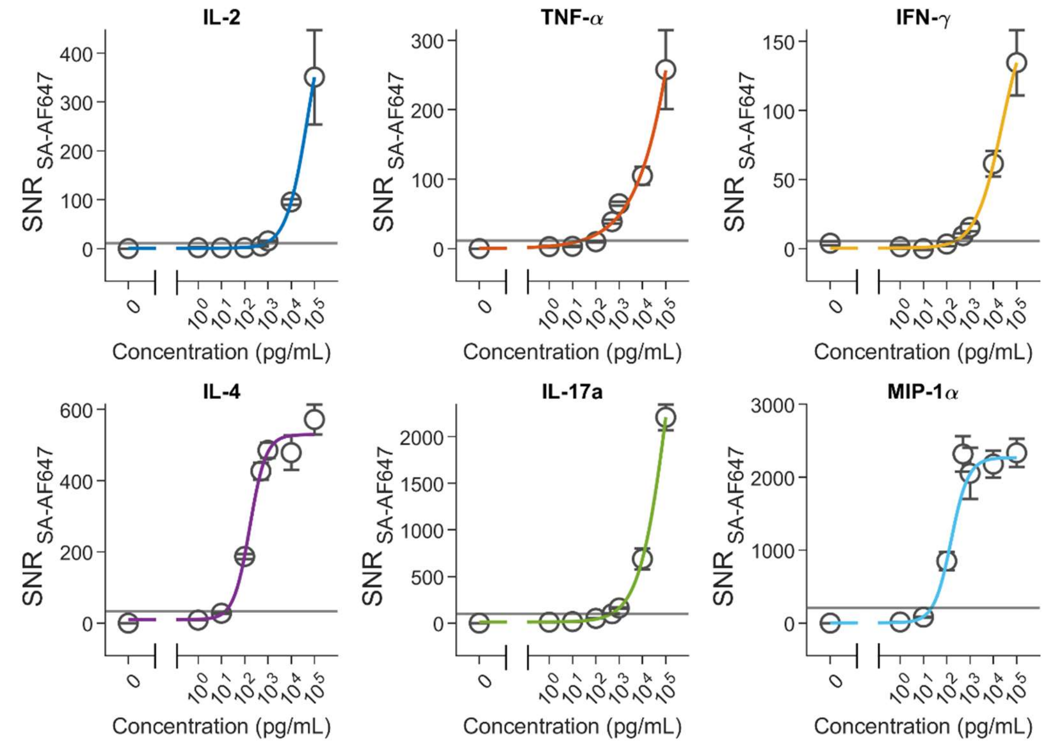

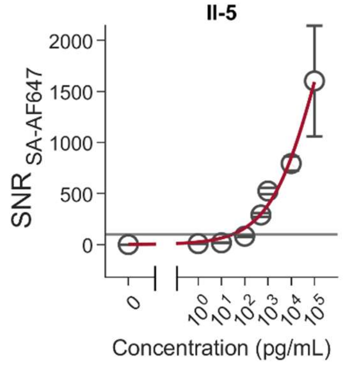

| LODs in multiplexed assay (pg/mL) | |
|---|---|
| **Cytokine** | **Bulk** |
| IL-2 | $8.01 \times 10^2$ |
| TNF-α | $2.19 \times 10^1$ |
| IFN-γ | $2.23 \times 10^2$ |
| IL-4 | $1.46 \times 10^1$ |
| IL-17a | $4.50 \times 10^2$ |
| MIP-1α | $2.23 \times 10^1$ |
| IL-5 | $2.84 \times 10^1$ |

**Figure S28**: Binding curves of cytokines measured by antibody microarray. Grey lines are calculated LOD, based on 3-fold the standard deviation of the assay blank above background. LOD for each cytokine is summarize in the inset table. Fits are four-points logistic regressions. Mean ± S.D. ($n$ = 10) are presented.

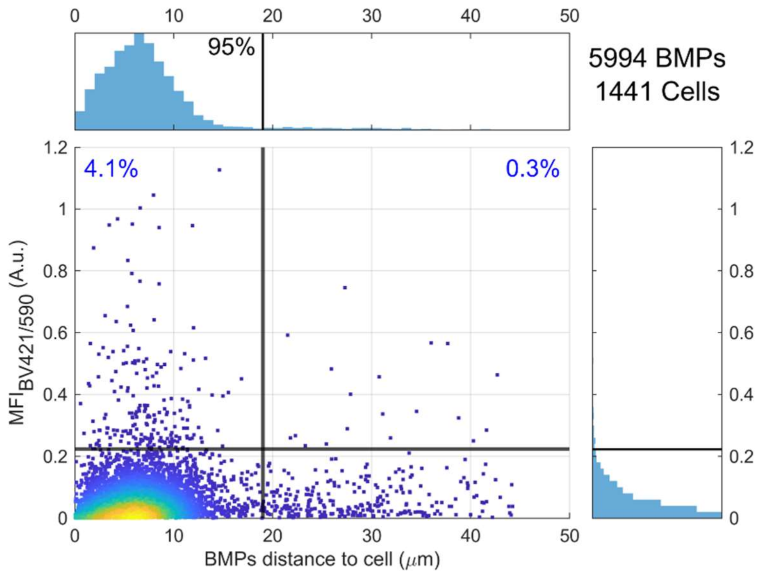

**Figure S29:** BMPs inside PiPs remained close to their co-encapsulated cell. Scatter plots presenting the assay signal intensity as a function of the distance between the cell boundary and the center of the BMP. The vertical line highlights the distance of 19 μm, which correspond to the limit at which 95% of the BMPs population is contained. Conversely, the horizontal line located around 0.22 marks the average assay LOD across all cytokines assayed in PiPs. Numbers in quadrant indicate percentage of BMPs population above average assay LOD prior accounting for technical replicate within the PiPs.

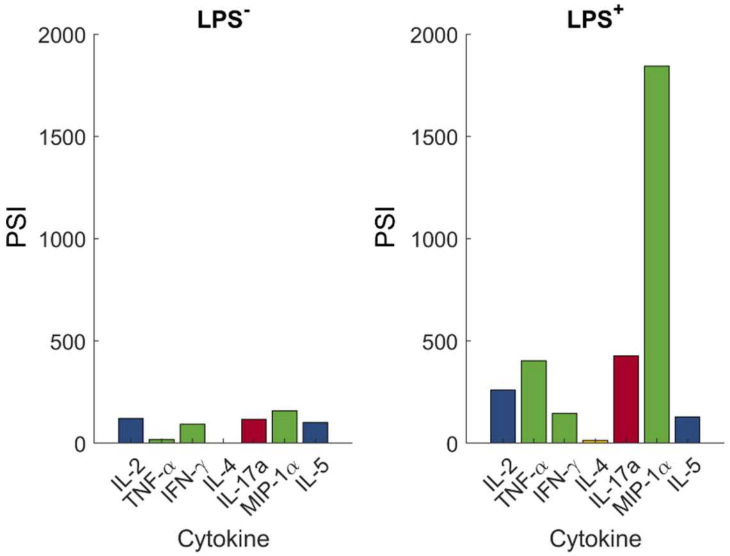

**Figure S30** Differentiated-THP-1 cell polyfunctional strength index (PSI) profile measured by PiP-plex across all samples separated into individual cytokines contribution, without and upon stimulation by 100 ng/mL LPS. Only cells secreting at least 2 cytokines were considered.

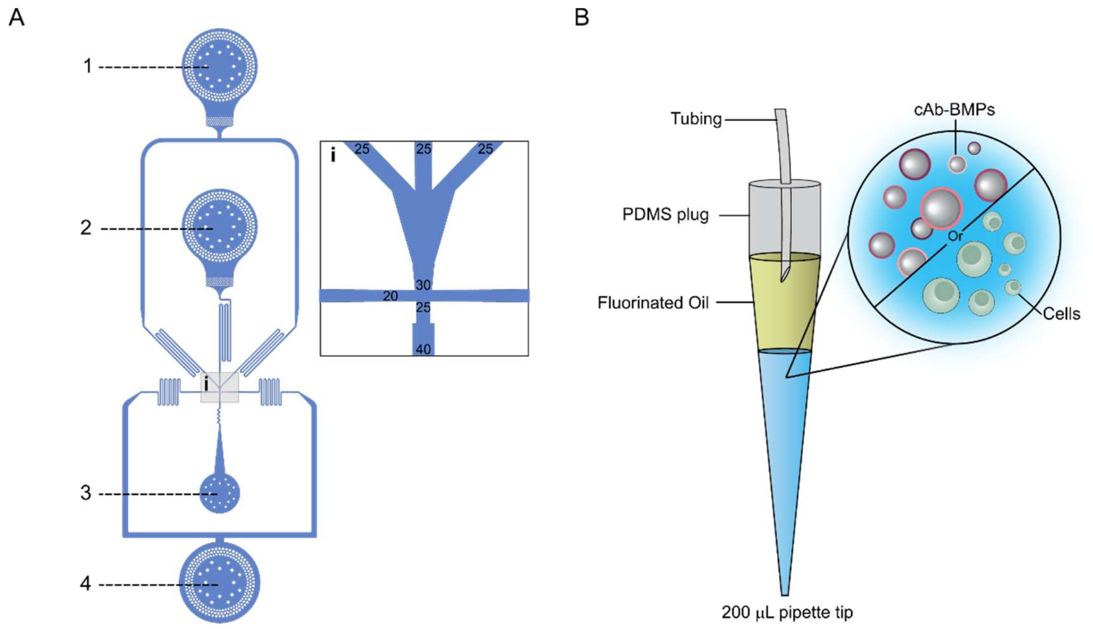

**Figure S31**: Encapsulation of cells and/or BMPs with soluble alginate by a microfluidic droplet generator. (**A**) Schematic presenting the geometry of the droplet generator used to generate the PiPs and (**B**) the strategy for loading ~30 µL volume reagents (soluble alginate, BMPs and cells) for W/O droplet generation by microfluidics using 200 µL pipette tips. By loading and storing reagents vertically above the microfluidic chip, sedimentation of BMPs and cells happened on chip rather than inside connecting tubing or syringes. This strategy improved yield of encapsulation of cells and BMPs during droplet generation. (**A**) Droplet generator: (1) and (2) are aqueous inlets, (3) correspond to the outlet and (4) to the oil inlet. Inset *i* presents a zoomed schematic describing key sizes of the droplet generator. Values are in micrometers. Height of the channels is 35 µm. (**B**) cAb-BMPs or live cells in soluble alginate are loaded into 200 µL pipette tips, pre-loaded with fluorinated oil. The tubing is directly connected to a ½27G blunt needle and mounted on a syringe pump for infusion of the reagents into the microfluidic chip to generate the PiPs.
